# Recruitment of the m^6^A/Am demethylase FTO to target RNAs by the telomeric zinc finger protein ZBTB48

**DOI:** 10.1101/2024.01.15.575768

**Authors:** Syed Nabeel-Shah, Shuye Pu, Giovanni L. Burke, Nujhat Ahmed, Ulrich Braunschweig, Shaghayegh Farhangmehr, Hyunmin Lee, Mingkun Wu, Zuyao Ni, Hua Tang, Guoqing Zhong, Edyta Marcon, Zhaolei Zhang, Benjamin J. Blencowe, Jack F. Greenblatt

**Affiliations:** Donnelly Centre, University of Toronto, Toronto, M5S 3E1, Canada; Department of Molecular Genetics, University of Toronto, Toronto, M5S 1A8, Canada; Department of Computer Sciences, University of Toronto, Toronto, M5S 1A8, Canada

**Keywords:** TZAP, ZBTB48, mRNA modification, m6A/m6Am, FTO, iCLIP, Telomeres, TERRA, RNA-binding

## Abstract

N6-methyladenosine (m6A), the most abundant internal modification on eukaryotic mRNA, and N6, 2′-O-dimethyladenosine (m6Am), are epitranscriptomic marks that function in multiple aspects of posttranscriptional regulation. Fat mass and obesity-associated protein (FTO) can remove both m^6^A and m6Am; however, little is known about how FTO achieves its substrate selectivity. Here, we demonstrate that ZBTB48, a C2H2-zinc finger protein that functions in telomere maintenance, associates with FTO and binds both mRNA and the telomere-associated regulatory RNA TERRA to regulate the functional interactions of FTO with target transcripts. Specifically, depletion of ZBTB48 affects targeting of FTO to sites of m6A/m6Am modification, changes cellular m6A/m6Am levels and, consequently, alters decay rates of target RNAs. ZBTB48 ablation also accelerates growth of HCT-116 colorectal cancer cells and modulates FTO-dependent regulation of Metastasis-associated protein 1 (MTA1) transcripts by controlling the binding to MTA1 mRNA of the m6A reader IGF2BP2. Our findings thus uncover a previously unknown mechanism of posttranscriptional regulation in which ZBTB48 co-ordinates RNA-binding of the m6A/m6Am demethylase FTO to control expression of its target RNAs.

## Background

Among more than 100 different chemical modifications in RNA, m6A is the most abundant in mRNA and has been linked to multiple posttranscriptional regulatory pathways, including mRNA splicing, mRNA export, mRNA decay, and translation [1–5]. It is incorporated in mRNA by a multi-component methyltransferase complex, which includes a Mettl3-Mettl14 core complex, WTAP, VIRMA, HAKAI, RBM15, and ZC3H13. Reversal of m6A is mediated by FTO and ALKBH5 [6–10]. Recently, it was shown that telomeric repeat-containing RNAs (TERRA) contain m6A modifications that play an important role in telomere maintenance [11]. Consistent with a role for m6A in multiple posttranscriptional pathways, links are increasingly being found between dysregulation of m6A and various human diseases, including cancer [12,13].

Transcriptome-wide studies have shown that m6A sites are enriched in 3’UTRs and near stop codons and have identified a DRACH (D = A, G, U; R = A, G; H = A, C, U) motif as the major m6A consensus sequence [14,15]. It has been estimated that ∼0.1–0.5% of total adenosine in polyadenylated RNA is methylated [8]. Many of the above-noted m6A functions are mediated by “reader” proteins that specifically recognize m6A in mRNAs. YTH-domain proteins, including YTHDF1-3, YTHDC1 and YTHDC2, are considered the primary m6A readers [10], as they can bind directly to m6A [16]. In addition to these direct m6A binders, certain context-specific readers, including heterogeneous nuclear ribonucleoproteins (HNRNPs) [17,18] and insulin-like growth factor 2 mRNA-binding proteins (IGF2BPs) [19], have also been identified. m6A can alter the local structure in mRNAs and long non-coding RNAs (lncRNA), which can facilitate recruitment of certain context-specific reader proteins, e.g., hnRNP C [20]. The effects on target mRNAs can vary depending on the function(s) of the specific m6A-reader protein; for example, the binding of YTHDF2 destabilizes its target mRNAs [21], whereas IGF2BP2 enhances the stability of its bound mRNAs [19].

When the first transcribed nucleotide at the mRNA 5’-end is adenosine, it can be methylated at the 2′-hydroxyl group and further methylated at the N6 position, thus giving rise to m6Am [22,23] at ∼50–80% of mRNAs that start with adenosine [22]. Although the biological role(s) of m6Am has remained somewhat enigmatic, it is thought to regulate both mRNA stability and translation [24–26]. Recently, it was shown that m6Am is important for stem-like properties in colorectal cancer cells [27]. m6Am is installed at the first transcribed adenosine, and a BCA consensus sequence (where B denotes G or C or T, and A is the methylated adenosine) has been identified as the most prominent motif [28]. It was recently shown that the phosphorylated CTD interacting factor 1 (PCIF1) is the sole methyltransferase that installs m6Am on nascent transcripts [26,29–31]. FTO, which can remove internal m6A modifications, also functions as the m6Am demethylase [25], and m6Am appears to be its preferred target *in vitro* [25,32]. Nevertheless, since internal m6A is ten-fold more abundant than m6Am in cells, it has been suggested that more internal m6A than m6Am sites are demethylated by FTO [32].

The precise mechanism through which FTO achieves its substrate selectivity and binds to m6A (and/or m6Am)-containing mRNAs has remained largely ambiguous [33,34]. While structural context has been suggested to be important for the catalytic activity of FTO [35], the demethylation activities of FTO and ALKBH5 are not strictly dependent on DRACH sequence motifs [36]. Consistently, a previously reported FTO UV crosslinking and immunoprecipitation followed by sequencing (CLIP-seq) analysis failed to identify DRACH motifs [37], suggesting that the targeting of FTO to m6A might not be strictly sequence-dependent. Moreover, FTO was not identified when RNA probes containing four repeats of the m6A consensus sequence ‘GGACU’ were used to pull down cellular proteins that bind to DRACH motifs [38]. The subcellular localization of FTO, which appears to differ between tissue- and/or cell-types, has been suggested to play a role in determining its target preference [34]: for example, FTO is nuclear in HEK293 cells [37], whereas it localizes to both cytoplasm and nucleus in colon cancer cells, where it appears to regulate m6Am in the cytoplasm [27].

ZBTB48 (also known as telomeric zinc finger-associated protein (TZAP) or human Krüppel-related 3 (HKR3)), a Cys2-His2 zinc finger protein (C2H2-ZFP) which contains a POZ domain at its N-terminus and 11 C2H2 zinc finger domains towards its C-terminus [39], preferentially binds to the telomeric DNA repeats of long telomeres and replaces telomeric repeat–binding factors 1 and 2 (TRF1 and TRF2). ZBTB48 overexpression results in progressive telomere shortening, whereas its silencing in mouse embryonic stem cells causes telomere elongation [40,41]. Moreover, ZBTB48 can also function as a transcriptional activator for a small set of genes [40]. Furthermore, ZBTB48 has been shown to function as an inhibitor of the self-renewal of porcine mesenchymal stromal cells, and it has also been implicated in the regulation of premature senescence [41]. Interestingly, ZBTB48 has been localized to chromosome 1p36, a region that is commonly rearranged or deleted in many cancers [42]. Although ZBTB48 has been suggested to function as a tumor suppressor [43,44], its precise function in tumor cells remains unknown.

Here, we show that ZBTB48 interacts physically with the m6A/m6Am demethylase FTO and modulates its binding to target mRNAs. By utilizing individual nucleotide resolution CLIP-seq (iCLIP-seq), we demonstrate that ZBTB48 binds directly to mRNA in cells, independently of its DNA-binding activity, and that its RNA-binding sites are enriched around those of FTO. Moreover, ZBTB48 ablation affects binding of FTO to m6A/m6Am sites, and its overexpression reduces cellular m6A/m6Am levels. We further demonstrate by knockdown followed by RNA-sequencing that ZBTB48 depletion results in an average downregulation of the expression of m6A-containing mRNAs. These results delineate a new role for ZBTB48 in posttranscriptional regulation and implicate it as a regulator of m6A/m6Am.

## Results

### ZBTB48 interacts with the m^6^A/m6Am demethylase enzyme FTO

We first investigated the function (s) of ZBTB48 by exploring its protein-protein interaction (PPI) partners in our previously published affinity-purification followed by mass spectrometry (AP-MS) data in HEK293 cells [45]. Consistent with its known function(s) as a DNA-binding factor, proteins involved in multiple DNA-related processes, including DNA repair, DNA recombination, and chromosome organization, were identified as interaction partners of ZBTB48 (Figure S1A). Unexpectedly, we also identified the m6A/m6Am demethylase FTO as an interaction partner, suggesting a potential role for ZBTB48 in m6A/m6Am metabolism. We assessed the specificity of its interaction with FTO by filtering ZBTB48 AP-MS data against similar data for numerous GFP purifications and several DNA-binding transcription factor purifications (total controls: n=218) using the “Significance Analysis of INTeractome” (SAINTexpress) algorithm [46]. While the number of ZBTB48 interaction partners was then reduced to only 12 proteins that passed our statistical threshold of a Bayesian false discovery rate (FDR) ≤ 0.01 (Figure 1A; Table S1), FTO remained one of the highest-confidence interaction partners, indicating that FTO association with ZBTB48 is robust. Moreover, our analysis of the ‘Contaminant Repository for Affinity Purification’ (CRAPome) database did not identify FTO as a ‘frequent flyer’ protein in AP-MS experiments (Figure S1B) [47], excluding the possibility that FTO might be a general non-specific background protein in proteomic studies.

**Figure 1:**
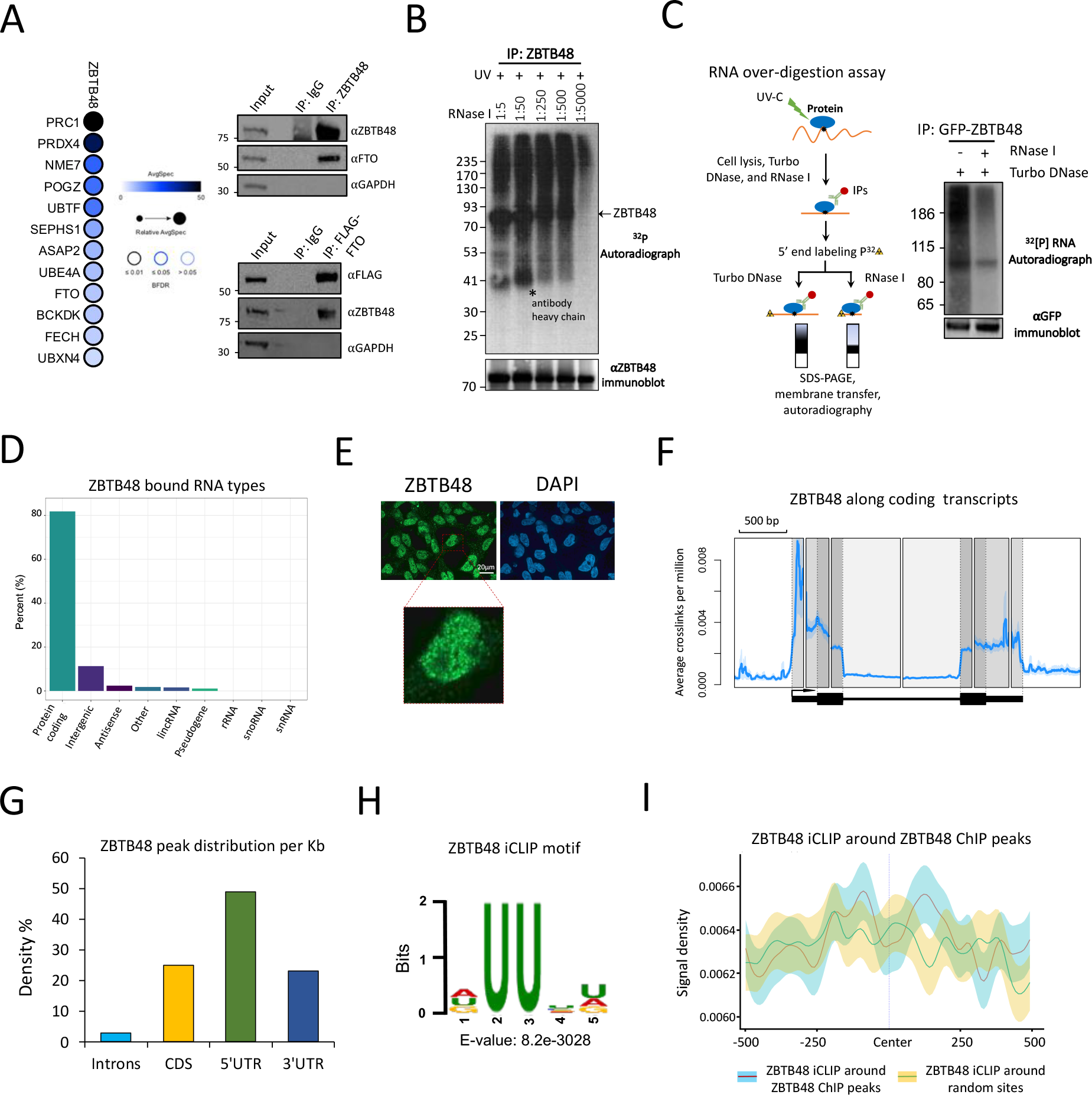
ZBTB48 interacts with FTO and binds to RNA in cells. A: Left, Dot plot representation of ZBTB48 protein-protein interactions. Right, Interaction of ZBTB48 with FTO detected by co-immunoprecipitation. Cell lysates were treated with Benzonase prior to IPs. Note: The inputs and IPs were loaded twice on 4-12% Bis-Tris SDS polyacrylamide gels and probed with the indicated antibodies. **B:** CLIP autoradiography of ^32^P-labeled ZBTB48-RNA complexes was performed after RNase I treatment at various dilution. The Western blot in the bottom panel shows the recovery of ZBTB48. **C:** Left, Schematic representation of CLIP RNA over-digestion assay. Right, Autoradiograph of immunopurified ^32^P-labeled ZBTB48-RNA complexes after RNase I and/or DNase I over-digestion. The bottom panel Western blot indicates the recovery of GFP-ZBTB48 protein. **D:** Bar chart representing the distribution (%) of ZBTB48-bound RNA types. **E:** Immunofluorescence (IF) analysis to examine the localization of ZBTB48 in HEK293 cells. Experiment was performed using an anti- ZBTB48 antibody. For nuclear counterstaining, DAPI was used. **F:** Standardized metaplot profile showing the density of ZBTB48 as average crosslinks per million. Black arrow indicates transcription start site (TSS). **G:** Bar chart representing the distribution of ZBTB48 mRNA CITS peaks per Kb of mRNA (FDR ≤ 0.01). **H:** Enriched sequence motif found in ZBTB48 iCLIP-seq peaks. E-value represents the significance of the motif against randomly assorted sequences. **I:** iCLIP-seq signal density (CITS) of ZBTB48 around either ZBTB48 ChIP-seq peaks or random sites. Shaded area indicates a standard error of mean (SEM). See also Figures S1, S2 and Tables S1, S2, and S3.

To validate the interaction between ZBTB48 and FTO, co-immunoprecipitation (co-IP) experiments were performed. Consistent with our AP-MS analysis, FTO co-immunoprecipitated with GFP-ZBTB48 (Figure S1C). To further confirm that endogenous ZBTB48 interacts with FTO, co-IPs were performed using antibodies against the endogenous ZBTB48. To test for possible indirect associations mediated by DNA or RNA, cell extracts were treated with a promiscuous nuclease (Benzonase) prior to IPs. In these experiments endogenous FTO co-immunoprecipitated with ZBTB48 (Figure 1A). Conversely, endogenous ZBTB48 was pulled down with FLAG-FTO (Figure 1A). To roughly map the FTO-interaction region of ZBTB48, we engineered HEK293 cells expressing ZBTB48 truncation mutants and subjected them to co-IPs. Both the full-length protein and the N-terminal region of ZBTB48 containing its BTB domain immunoprecipitated FTO, whereas the C-terminal region harbouring its zinc fingers (ZnFs) did not (Figure S1D). These results suggested that the BTB domain or an unstructured region of ZBTB48 might participate in its interaction with FTO. We conclude that ZBTB48 forms a direct or indirect PPI with FTO. This interaction is highly specific, since FTO did not co-purify with 120 other C2H2-ZFPs [45].

### ZBTB48 binds to mRNAs in cells

Because ZBTB48 directly or indirectly interacted with the RNA demethylase FTO, we examined whether ZBTB48 binds to RNA in cells. By utilizing CLIP followed by radio-labeling of RNA, SDS-PAGE, and autoradiography, we observed that ZBTB48 crosslinks robustly to RNA in cells, specifically upon UV exposure. Importantly, the bulk of the radioactive signal migrated above the position of ZBTB48 when a low concentration of RNase I was used and increasingly migrated at the position of ZBTB48 as the RNase I concentration was increased (Figure 1B). Moreover, the radiolabelled RNA that crosslinked to ZBTB48 was substantially reduced when CLIP samples were over-digested with RNase I, but not DNase I, indicating that ZBTB48 binds directly to RNA in cells (Figure 1C). To further confirm that the RNA signal emerged from crosslinking of RNA to ZBTB48 and was not due to RNA-binding proteins or RNA interacting with ZBTB48 in extracts from UV-treated cells, we mixed UV-untreated cells expressing GFP-ZBTB48 with UV-treated wildtype cells before lysis and used them as a control in CLIP assays. Consistently, radioactive signal was observed only when the GFP-ZBTB48-expressing cells were treated with UV (Figure S1E). In contrast, neither GFP-alone nor the ‘mixed’ control samples displayed any appreciable radiolabeled RNA signal (Figure S1E). We then subjected ZBTB48 truncation mutants to CLIP assays and found that the C-terminal region containing its ZnFs strongly crosslinked to RNA, whereas the N-terminal region yielded a much weaker signal (Figure S1F). These results suggested that, like its DNA binding, ZBTB48 binds to RNA through its ZnFs.

To identify ZBTB48-bound RNAs in cells, we performed iCLIP-seq experiments using a previously validated anti-ZBTB48 antibody [40] (Figure S1G), in parallel with sequencing of size-matched input (SMI) samples. Biological replicates highly correlated with each other, indicating the reproducibility of our data (Figure S1H). In addition to the SMI samples, iCLIP-seq data from GFP-alone samples were employed to control for background signal. We defined ZBTB48 RNA-binding sites by identifying ‘crosslinking-induced truncation sites’ (CITS, FDR ≤ 0.01; Methods). The majority (∼80%) of the identified CITS peaks fall within protein-coding transcripts, indicating that ZBTB48 predominantly targets mRNAs in cells (Figure 1D). Without length normalization across various transcript features, ∼70% of crosslinking peaks were identified as intronic (Figure S1I; Table S2), consistent with introns being much longer than exons. The peaks within introns suggest binding to pre-mRNA and, consistently, immunofluorescence analysis showed that ZBTB48 is predominantly nuclear (Figure 1E). When we examined the average RNA-binding density after normalizing to the lengths of various features along target mRNAs using a composite analysis, most ZBTB48 crosslinks accumulated within 5’ and 3’UTRs (Figure 1F), whereas ∼24% of the peaks reside within coding sequences (CDS) (Figure 1G), suggesting that ZBTB48 might preferentially bind to the untranslated regions of target mRNAs. To examine if ZBTB48 has any sequence preferences, we searched for enriched motifs and identified a U-rich consensus sequence (_UU_ _) as the most significantly enriched motif. GGGAGG and _ _CCAGC were identified as the second and third most over- represented motifs, respectively (Figure 1H; Figure S1J).

Since ZBTB48 binds to telomeric DNA repeats and can also function as a transcription factor for a few genes [40,41], we examined whether its DNA- and RNA-binding sites are correlated. ChIP-seq experiments were performed with two biological replicates to identify ZBTB48 DNA-binding sites (Figure S2A; Table S3). Even though our ChIP-seq data were generated using GFP-ZBTB48 in HEK293 cells, and we employed enrichment over input to identify significant peaks, the ChIP peaks we identified were enriched around previously published ChIP sites that were detected for endogenous ZBTB48 in U2OS cells using ZBTB48 knockout cells as a control (Figure S2B) [40]. Conversely, previously published ChIP sites were enriched around our identified ZBTBTB sites (Figure S2B). These observations indicate that our data captured the reported ZBTB48 DNA-binding profile. When the RNA-crosslink sites were plotted around the ChIP-seq peaks, ZBTB48 RNA-binding density was not enriched around its DNA-binding sites (Figure 1I), and only ∼5% of ChIP-seq peaks overlapped with the iCLIP-seq peaks when the peaks were extended by 50 nucleotides on each side (Figure S2C). Thus, ZBTB48 appears to have distinct, mostly non-overlapping, DNA- and RNA-binding sites, although there is a small degree of enrichment specifically in 3’ and 5’ UTRs (Figure S2D).

In addition to our use of a previously validated antibody [40], as well as our additional validations of ZBTB48 antibody using ZBTB48 knockdown cells (see below), we independently confirmed RNA-binding by ZBTB48 by performing RNA-immunoprecipitation (RIP) followed by qRT-PCR experiments utilizing GFP-ZBTB48-expressing cells. As shown in Figure S2E, precipitation of GFP- ZBTB48 successfully pulled down 8/8 of the target transcripts, as defined by our iCLIP-seq experiments, that we examined, whereas precipitation of GFP-alone did not. In contrast GFP- ZBTB48 did not show significant binding to two non-targets (Figure S2E). We conclude that ZBTB48 binds directly to mRNAs in cells and that binding is mostly distinct from its DNA-binding activity.

### ZBTB48 and FTO RNA-binding sites coincide

Since ZBTB48 associated with FTO, we next examined whether their RNA-binding sites coincide on mRNAs. We performed FTO iCLIP-seq experiments, with four biological replicates, using inducible Flp-In T-REx HEK293 cells expressing FLAG-tagged FTO (Figure S2F; Table S4). Consistent with previous studies [37], we found that most FTO CITS peaks (∼80%) fall within introns (Figure 2A, left), whereas, after normalizing to the lengths of various transcript regions, ∼28%, ∼34%, and ∼28% of FTO peaks map within 5’UTRs, 3’UTRs, and CDS, respectively (Figure 2A, right) [33]. To examine whether ZBTB48 crosslink sites coincide with FTO binding sites, we plotted the ZBTB48 CITS around the FTO RNA-binding positions. Importantly, ZBTB48 iCLIP signal was significantly enriched in the vicinity of FTO-binding sites (Figure 2B, p ≤ 0.001, Wilcoxon test; Figure S3A, left). Furthermore, ZBTB48 signal was also enriched around FTO peaks when previously published CLIP-seq data was used to map FTO RNA-binding sites (Figure S3B) [37].

**Figure 2:**
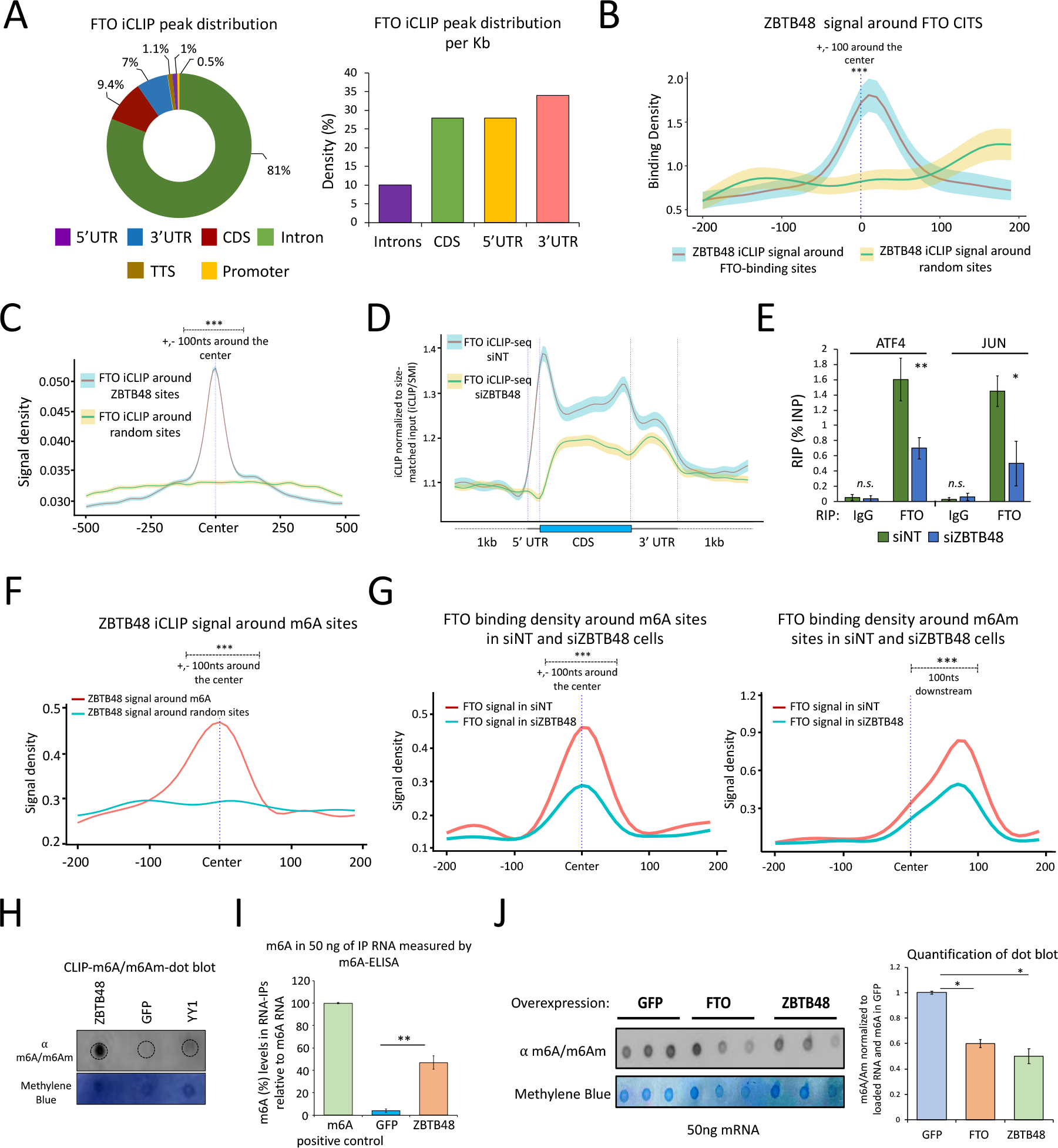
RNA-binding sites of ZBTB48 and FTO coincide on target transcripts. A: Left, FTO iCLIP-seq CITS distribution. Absolute numbers of peaks are used. Right, Bar chart showing the distribution of FTO iCLIP-seq peaks per Kb along mRNA. **B:** Metagene profile showing ZBTB48 iCLIP-seq signal density around either FTO RNA-binding sites or random sites. Quantification of signal densities was performed for +/-100bp around the center (∗∗∗p ≤ 0.001, Wilcoxon (Mann- Whitney) test). Shaded area: SEM. **C:** FTO iCLIP-seq signal density around either ZBTB48 CITS or random sites (∗∗∗p ≤ 0.001, Wilcoxon test). **D:** Standardized metaplot profiles showing the binding density (iCLIP/size-matched input) of FTO in either siZBTB48 or siNT cells. **E:** Bar graph showing FTO RIP-qPCR experiments after knocking down ZBTB48 (biological replicates (n) = 4, student’s t test, ∗∗p ≤ 0.01, ∗p ≤ 0.05, error bars: SEM). **F:** ZBTB48 iCLIP-seq signal density around either m6A sites or random sites. **G:** FTO signal density around m6A (left) or m6Am (right) sites in cells treated with either siZBTB48 or siNT. Note: For F and G, quantification of signal densities was performed for +/-100bp around the center; ∗∗∗p ≤ 0.001, Wilcoxon test. **H:** ZBTB48 CLIP dot blot showing the enrichment of m6A/m6Am modified RNA. Top panel: anti-m6A, bottom panel: Methylene blue. **I:** Bar plot showing the enrichment of m6A. RIP was performed using either GFP- ZBTB48 or GFP-alone cells, and RNA was analyzed using an m6A ELIZA kit. Synthetic RNA containing m6A was used as a positive control (error bars: SEM, n =3, student’s t test, ∗∗p ≤ 0.01). **J:** m6A dot blot using polyadenylated RNA purified from either GFP-alone, or ZBTB48- overexpressing, or FTO-overexpressing cells (top panel: anti-m6A, bottom panel: Methylene blue, error bars: SEM, ∗p ≤ 0.05; student’s t-test; n = 3). See also Figure S3, S4 and Table S4.

Consistently, when examining the RNA-binding profile of FTO around ZBTB48 crosslink sites, we found that FTO signal was also significantly enriched around ZBTB48 RNA-binding sites (Figure 2C), and this enrichment was observed for 5’UTR, CDS, and 3’UTR regions (Figure S3C, p ≤ 0.001, Wilcoxon test). Consistent with their overlapping RNA-binding sites, the majority of the identified target transcripts were also shared between the two proteins (Figure S3A, right). To further examine the specificity of FTO enrichment around ZBTB48 sites, we then utilized our previously reported iCLIP-seq data for SP1 [48]. Consistently, SP1, which we have previously shown to function in alternative polyadenylation [48], was not significantly enriched around ZBTB48 sites (Figure S3D). Moreover, FTO peaks were depleted around ZBTB48 DNA-binding sites (Figure S3E), perhaps because transcription factor-binding regions in promoters or elsewhere are not transcribed, indicating that the ZBTB48-FTO interaction may not normally occur in the context of the DNA-related functions of ZBTB48. We conclude that the ZBTB48 and FTO RNA-binding sites are coincident across the transcriptome, consistent with our observation of a direct or indirect physical association between these factors. Collectively these observations suggest possible co-regulatory potential for these proteins on their common target mRNAs.

### ZBTB48 modulates FTO RNA-binding

To address their possible functional interaction, we assessed whether ZBTB48 affects the RNA-binding activity of FTO by performing FTO iCLIP-seq in HEK293 cells, in 2 biological replicates, after knockdown (KD) with a ZBTB48 siRNA or non-targeting siRNAs (siNT) (Figure S4A). Western blotting analysis using anti-ZBTB48 antibody confirmed that the ZBTB48 protein levels were specifically depleted in siZBTB48-treated cells (Figure S4B). The expression of FTO remained unaffected upon ZBTB48 KD at the levels of both mRNA and protein (Figure S4B). Moreover, the subcellular localization of FTO did not change in HEK293 cells following ZBTB48 KD (Figure S4C). In siNT treated cells, FTO’s RNA-binding density was particularly enriched in the 5’UTRs (Figure 2D), consistent with previous studies [33,37] and results described above. In contrast, the RNA-binding density of FTO was substantially reduced all along target transcripts in siZBTB48 cells, and especially in the 5’UTR, even though the number of uniquely mapping reads was comparable across both conditions (Figures 2D and S4D; Table S5). These results suggested that ZBTB48 recruits FTO to target transcripts in cells.

Since RNA-binding sites of FTO often coincide with those of ZBTB48, we also examined whether depleting ZBTB48 affects FTO binding to these sites. When we plotted FTO iCLIP-seq signal around ZBTB48-binding sites, we observed a reduction in the RNA-binding density of FTO in siZBTB48 cells in comparison with control cells (Figure S4E, p ≤ 0.001, Wilcoxon test). Furthermore, FTO RIP-qPCR experiments in cells treated with a different siRNA against ZBTB48 (siZBTB48 # 2) (Figure S4A and S4F) showed a significant reduction in FTO signal for the two FTO targets we examined in ZBTB48-depleted cells (Figure 2E, p ≤ 0.01, student’s t-test). In contrast, RNA-binding by ZBTB48 in FTO depleted cells remained unaffected for the examined target transcripts (Figure S4G). We conclude that ZBTB48 helps recruit FTO to target mRNAs in cells.

### ZBTB48 affects FTO RNA-binding around m6A/m6Am sites

We next utilized published m6A-iCLIP-seq (miCLIP-seq) data [28] to ask whether ZBTB48 affects binding of FTO around m6A/m6Am sites in cells. FTO showed an enrichment around m6A and m6Am sites, as assessed using both our FTO iCLIP-seq and previously reported FTO CLIP-seq data (Figures S5A-C) [37]. Consistently, ZBTB48 also exhibited strong enrichments around m6A and m6Am sites (Figures 2F and S5D). In contrast to FTO and ZBTB48, however, none of the other RNA-binding proteins we examined, including U2AF1, PTBP1, MKRN2 [49], and SP1 [48], showed any enrichment around m6A sites (Figure S5E; see Methods). We then utilized our FTO iCLIP-seq data generated in siZBTB48 or siNT cells and observed that FTO binding around m6A/m6Am sites was reduced in ZBTB48-depleted cells in comparison with the control cells (Figure 2G, p ≤ 0.001, Wilcoxon test). These results suggested that ZBTB48 might reduce cellular m6A/m6Am levels by recruiting FTO around m6A/m6Am sites on target mRNAs.

### ZBTB48 binds m6A/m6Am-containing RNAs and regulates cellular m6A/m6Am via FTO

Since the above results indicated that ZBTB48 binds near m6A/m6Am sites and co-ordinates the targeting of FTO to at least a subset of methylated sites, we examined whether ZBTB48 preferentially associates with m6A/m6Am-modified RNAs. We first carried out additional validations on a previously characterized anti-m6A antibody through a dot-blot assay using total RNA derived from cells treated either with siMETTL3 or control siRNAs (Figures S5F and S5G) [14,15]. Note that the anti-m6A antibody does not distinguish between internal m6A and 5’ cap-associated m6Am. We then used this antibody to probe RNA isolated in ZBTB48 CLIP experiments. Consistently, GFP-ZBTB48 immunoprecipitated RNAs with far more m6A/m6Am-modificationscompared to GFP-alone (Figure 2H). YY1, another C2H2-ZFP that is known to bind RNA [50], was used as an additional specificity control in these experiments and failed to show any appreciable m6A/m6Am signal (Figure 2H). To further examine the ability of ZBTB48 to target m6A/m6Am-modified RNAs, we performed GFP- ZBTB48 RIP experiments with anti-GFP followed by ELISA-based assays for m6A. Consistently, we observed that, in comparison with the GFP control, RNA immunoprecipitated with GFP-ZBTB48 was m6A/m6Am modified (Figure 2I). These results support the conclusion that ZBTB48 binds near m6A/m6Am residues on its target RNAs.

Since ZBTB48 targeted methylated transcripts and modulated the mRNA-binding activity of FTO, we examined if ZBTB48 influences total m6A/Am levels in mRNA. We isolated PolyA-enriched mRNAs from cells overexpressing GFP-ZBTB48, GFP-FTO, or GFP (Figure S6A) and subjected these mRNAs to m6A/m6Am dot-blot assays. As expected, FTO overexpression resulted in a significant reduction in global m6A/m6Am levels in mRNAs (Figure 2J). Notably, in comparison with control cells expressing GFP, m6A/m6Am levels in cellular mRNAs were also significantly reduced upon GFP-ZBTB48 overexpression (Figure 2J, p ≤ 0.05, student’s t-test). These results are consistent with the idea that ZBTB48 co-ordinates FTO-binding to target RNAs, thus indirectly regulating cellular m6A/Am levels.

### ZBTB48 modulates FTO-mediated transcriptome-wide demethylation of mRNAs

To further assess the potential impact of ZBTB48 on cellular m6A/m6Am at a transcriptome level, we carried out miCLIP-seq experiments, in two biological replicates, using HEK293 cells overexpressing either GFP-ZBTB48, or FTO, or GFP-alone (Figures 3A and S6B; Tables S6, S7, and S8). Since ZBTB48 and FTO both appear to target pre-mRNAs, we isolated total cellular RNA in these experiments, instead of using poly-A enriched processed mRNAs. The majority of identified CITS (FDR ≤ 0.01) fell within protein-coding genes with a strong enrichment near the stop codons and 3’UTR, as well as a smaller enrichment in the 5’UTR, and the DRACH sequence was found as the most significantly enriched motif in each dataset (Figure S6B), consistent with previous studies [28].

**Figure 3:**
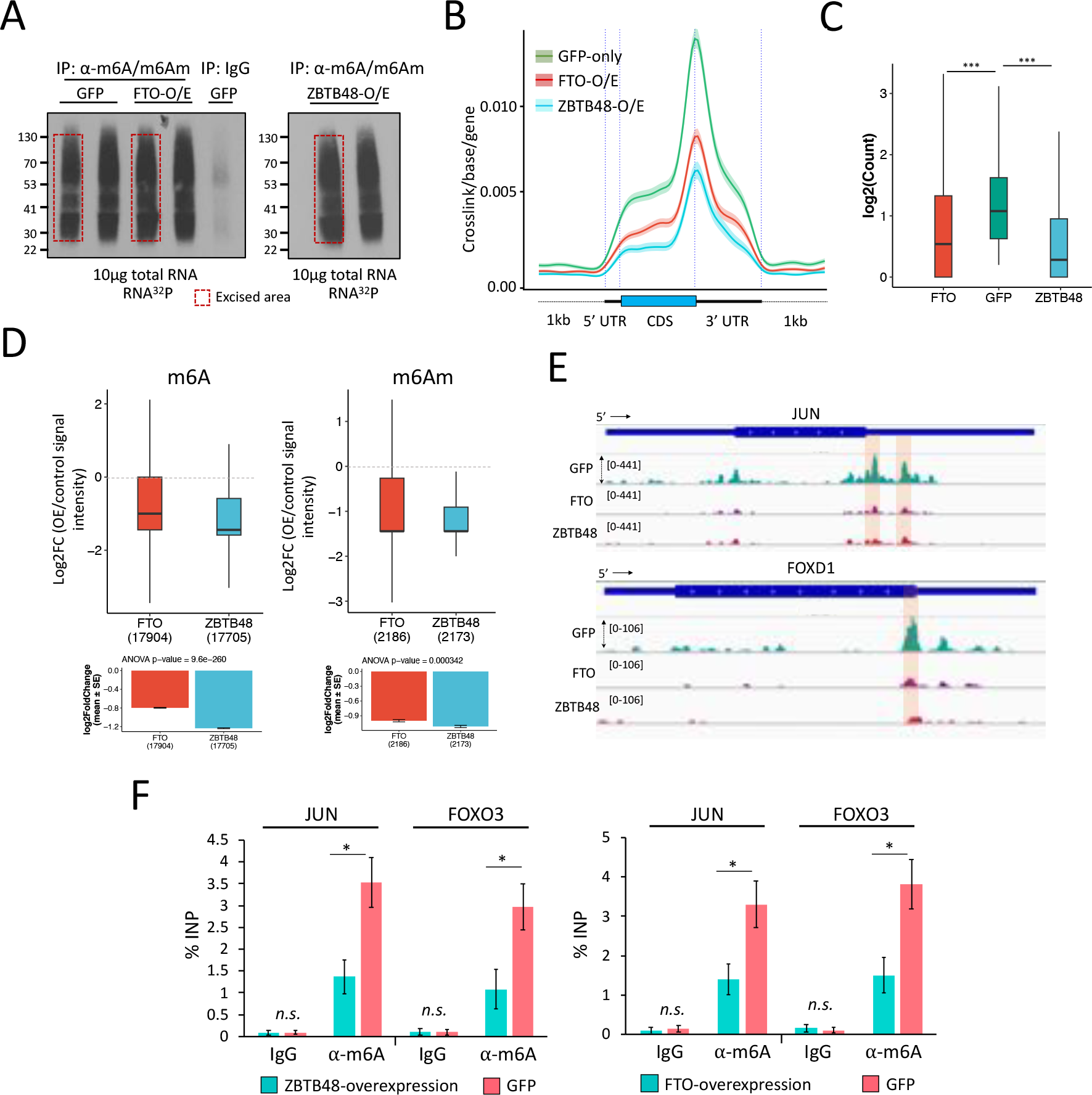
ZBTB48 affects cellular m6A/m6Am levels. A: Autoradiographs of ^32^P-labeled immunopurified (IP) anti-m6A-RNA complexes. Red boxes indicate the excised areaS for Standardized metagene plots showing miCLIP-seq signal density for RNA purified from cells overexpressing GFP-alone, or GFP-ZBTB48, or GFP-FTO. **C:** Boxplot representation of read counts for DRACH motif containing m6A sites across different samples, as indicated (Mann–Whitney test, ∗∗∗p ≤ 0.001; outliers not shown in the plot). **D:** Box plot showing differential methylation analysis using miCLIP data generated from either GFP-ZBTB48-overexpressing or GFP-FTO-overexpressing cells versus GFP control cells. Analysis was performed with the DESeq2 package, which accounts for the read depth in each sample, using reads corresponding to internal m6A (left) and terminal m6Am (right) sites. Outliers are not shown in the box plots. Dotted lines indicate the base line signal (i.e., m6A or m6Am in GFP cells). Bottom bar plots indicate the mean values for the indicated groups along with p-values (ANOVA, post hoc Tukey HSD). **E:** Genome browser snapshots of JUN and FOXD1 gene tracks show the location and coverage of miCLIP in GFP- or GFP-ZBTB48- or FTO-overexpressing cells. Locations of the putative m6A sites are highlighted. **F:** Bar plots showing the m6A levels for JUN and FOXO3 transcripts estimated using m6A-RIP-qPCR on RNA purified from either GFP-ZBTB48-overexpressing or GFP-only cells (left) or from either FTO-overexpressing or GFP-only cells (right). Data are represented as % input. Note, for qPCRs: biological replicates n = 5, student’s t-test, ∗∗∗p ≤ 0.001, ∗∗p ≤ 0.01, ∗p ≤ 0.05, n.s.: non-significant, error bars denote SEM. See also Figures S5, S6 and Tables S5-S8.

By comparing their signal densities, we found that, on average, m6A/m6Am intensities were substantially reduced in FTO- or ZBTB48-overexpression samples when compared with GFP controls (Figure 3B). We next considered only DRACH-motif-containing peaks as the high-confidence m6A sites for further analyses (Figure S6C). Moreover, we also identified potential m6Am sites (see Methods for details) (Figure S6D). Consistently, we observed a significant reduction in both DRACH-motif-containing m6A sites in the 3’UTR, CDS, and 3’UTR, as well as m6Am, upon FTO- or ZBTB48-overexpression, in comparison with control samples (Figure 3C and Figures S6E and S6F, p ≤ 0.001, Mann–Whitney). Differential analysis revealed that both FTO- and ZBTB48- overexpression samples had substantially reduced RNA methylation levels relative to control GFP samples (Figures 3D, 3E, S6G and S7A), suggesting a global reduction in transcriptome methylation levels. When we overlapped the m6A-containing genes, ∼2200 genes that were present after GFP-overexpression appeared to lack a defined m6A CITS peak in FTO- or ZBTB48-overexpression samples (Figure S7B). We validated this change in m6A levels for two targets, JUN and FOXO3, using m6A-RIP-qPCR experiments in cells overexpressing either ZBTB48, or FTO, or GFP. Consistently, m6A levels were significantly reduced for both examined transcripts in ZBTB48- or FTO-overexpression samples in comparison with control GFP cells (Figure 3F, p ≤ 0.001, t-test).

We initially utilized ZBTB48- and FTO-overexpressing cells in our experiments for consistency, because elevated FTO levels have previously been shown to significantly decrease cellular m6A levels [32]. Although the expression of known Mettl3/14-complex subunits and the m6Am writer enzyme PCIF1, as well as YTH-family proteins, was not significantly changed (see below), we noted that the overexpression of ZBTB48 elevated FTO expression levels (Figure S7C). The observed increase in FTO expression does not appear to have been caused directly by the transcription factor-related functions of ZBTB48, since ZBTB48 did not target the FTO promoter in our ChIP-seq data (Table S3). ZBTB48 and FTO mRNA expression levels were comparable in wildtype cells (Figure S7D), suggesting that the observed increase in FTO levels upon ZBTB48 overexpression could simply reflect a feedback mechanism in which cells sense the need for additional FTO protein. However, this elevation of FTO could at least partly explain why overexpression of ZBTB48 causes reductions in the levels of RNA methylation.

Because of this issue, we also examined m6A/m6Am levels in siZBTB48- or siFTO-treated cells. Importantly, ZBTB48 knockdown did not significantly alter the expression level of FTO (Figure S4B), and vice versa (Figure S7E). Moreover, the expression of METTL3 and PCIF1 also remained unaffected in siZBTB48 or siFTO cells (Figure S7E). When we analyzed m6A-to-A and m6Am-to-A ratios in total RNA using quantification by LC-MS/MS, knockdown of ZBTB48 led to a significant increase in cellular m6A and m6Am levels in comparison with control knockdown samples (Figure 4A, p ≤ 0.05; t-test), and consistent with previous reports [32], FTO depletion also resulted in an increase in both m6A and m6Am. Conversely, LC-MS/MS confirmed that overexpression of ZBTB48 or FTO decreased m6A and m6Am in purified polyadenylated mRNAs as well as in total RNA samples (Figures S7F, S7G), consistent with the findings in Figure 2J. We validated these findings for two individual targets, JUN and FOXO3 by performing m6A-RIP followed by qRT-PCR experiments after knocking down ZBTB48 or FTO using different siRNAs (siZBTB48 # 2 and siFTO # 2) (Figure 4B, p ≤ 0.001; t-test).

**Figure 4:**
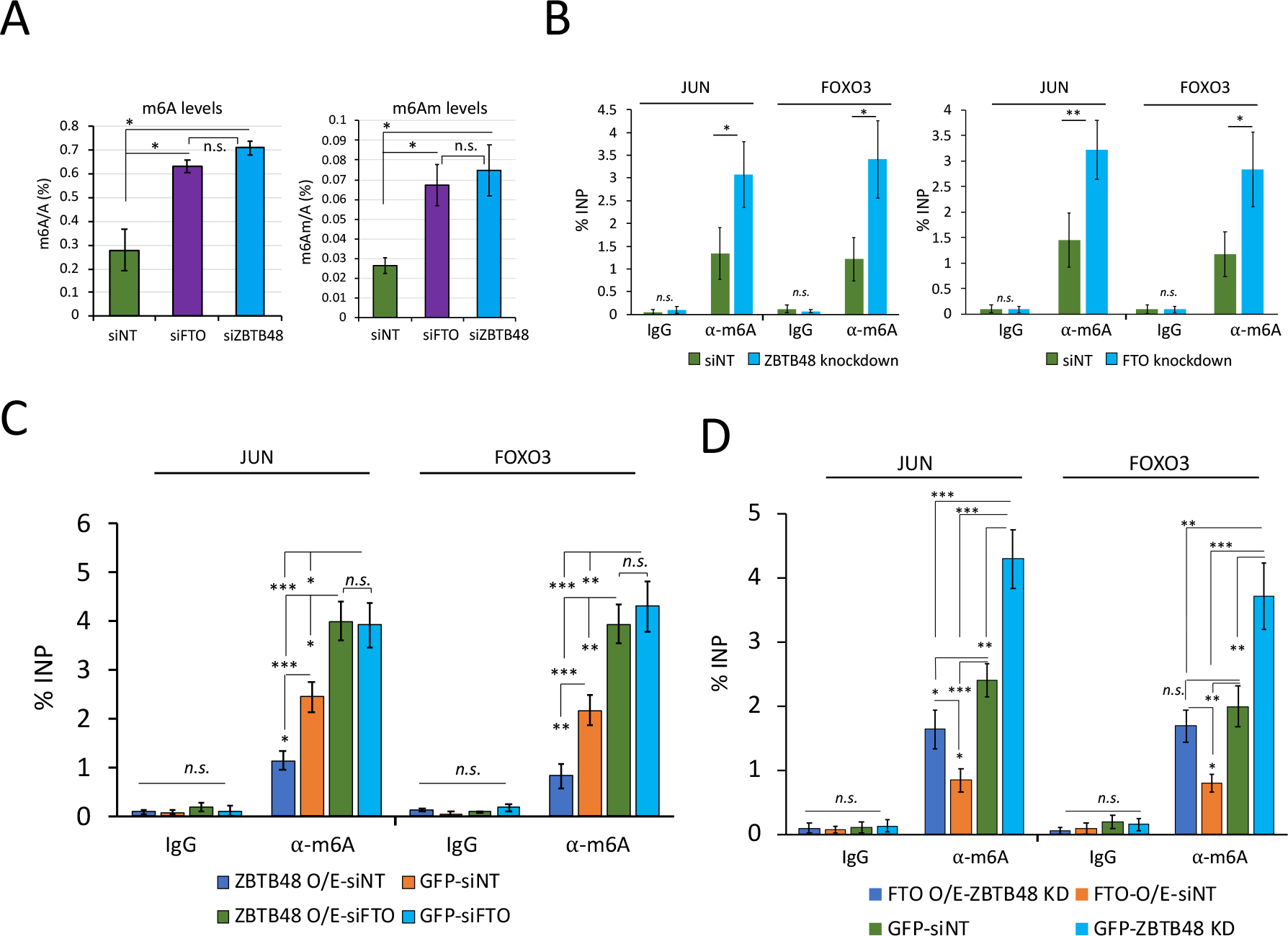
ZBTB48 knockdown increases cellular m6A/m6Am. A: LC-MS/MS shows that knockdown of ZBTB48 or FTO leads to an increase in m6A/A (left) and m6Am/A (right) in total RNA in comparison with the controls (i.e., siNT). Error bars show SEM (∗p ≤ 0.05; student’s t-test). **B:** Bar plot showing the m6A levels for JUN and FOXO3 transcripts estimated using m6A-RIP-qPCR on RNA purified from either siZBTB48- or siNT-treated cells (left) or from either siFTO- or siNT-treated cells (right). Data are represented as % input. **C-D:** Bar plots showing the m6A levels for JUN and FOXO3 transcripts estimated using m6A-RIP-qPCR on RNA purified from cells treated in the indicated ways. Data are represented as % input. Note, qPCRs in B-D: biological replicates n = 5, student’s t-test, ∗∗∗p ≤ 0.001, ∗∗p ≤ 0.01, ∗p ≤ 0.05, n.s.: non-significant, error bars denote SEM. See also Figures S6, S7 and Tables S5-S8.

To investigate whether ZBTB48-mediated modulation of m6A requires FTO, m6A-RIP-qPCR experiments were performed in GFP-ZBTB48-overexpressing cells treated either with siFTO or siNT. The results showed that neither JUN nor FOXO3 displays a significant decrease in m6A levels in ZBTB48-overexpressing cells when FTO is depleted (Figure 4C, Figure S7H). In fact, we observed a significant increase in the methylation levels for both JUN and FOXO3 transcripts, consistent with results described above indicating that FTO knockdown increases cellular m6A levels (Figure 4C). Expectedly, GFP-ZBTB48 overexpression in siNT-treated cells caused a significant reduction in m6A levels in comparison with GFP-expressing cells treated with siNT (Figure 4C). Conversely, m6A-RIP-qPCR experiments after knocking down ZBTB48 in FTO-overexpressing cells showed a slight reduction in m6A levels for both examined targets, although to a lesser extent than siNT-treated FTO-overexpressing cells (Figure 4D, Figure S7I). Furthermore, FTO overexpression in siNT-treated cells resulted in a significant reduction in m6A in comparison with GFP cells treated with siNT (Figure 4D). The observation that FTO-overexpression alone is sufficient to cause a decrease in cellular m6A levels in ZBTB48-depleted cells is consistent with previous CLIP-seq studies indicating that exogenously overexpressed FTO binds to RNA and targets m6A motif-containing sites more effectively [33]. In effect, overexpression of FTO can partially bypass the targeting effect of ZBTB48. We conclude that ZBTB48 regulates cellular m6A/m6Am levels by facilitating binding of FTO to target mRNAs.

### ZBTB48 impacts m6A in TERRA

Recently, it was shown that telomeric repeat-containing RNA (TERRA), which is transcribed from telomeres and functions in telomere maintenance through R-loop formation, is m6A-modified in a METTL3-dependent manner [11,51]. Considering that ZBTB48 regulates telomere metabolism [40,41], we examined whether ZBTB48 modulates TERRA m6A in an FTO-dependent manner. We utilized U2OS cells, since these cells have long recombinant telomeres [52], and ZBTB48 has been shown to localize to telomeres in these cells [40,41]. Moreover, we found that FTO is predominantly nuclear in these cells (Figure S8A). To examine the interaction between ZBTB48 and FTO in U2OS cells, co-IP experiments were performed. In these experiments, endogenous FTO co-precipitated with endogenous ZBTB48, consistent with their direct or indirect interaction in HEK293 cells (Figure S8B). We then performed RIP-qPCR and examined the binding of ZBTB48 to TERRA transcribed from 15q and 6q. Both 15q- and 6q-TERRA were significantly enriched in ZBTB48 samples in comparison with IgG controls (Figure 5A, p ≤ 0.001; t-test). Although we cannot exclude the possibility that ZBTB48 is binding R-loops rather than RNA, these experiments indicate that TERRA associates with ZBTB48. FTO binding to both 15q- and 6q-TERRA was significantly reduced when ZBTB48 was knocked down, in comparison with control cells (Figure 5A, p ≤ 0.01; t-test), even though knockdown of ZBTB48 did not significantly alter FTO expression levels (Figures S8C-S8E). Overexpression of FTO significantly decreased m6A levels for both 15q- and 6q-TERRA (Figure 5B), consistent with previous studies indicating that targeting of FTO to TERRA removes m6A [51]. Importantly, m6A levels were significantly increased for both 15q- and 6q-TERRA in siZBTB48- treated cells (Figures 5B and S8F). These results are consistent with the idea that ZBTB48 often regulates m6A by recruiting FTO and suggest that ZBTB48 might function in telomere maintenance by regulating m6A modification of TERRA.

**Figure 5:**
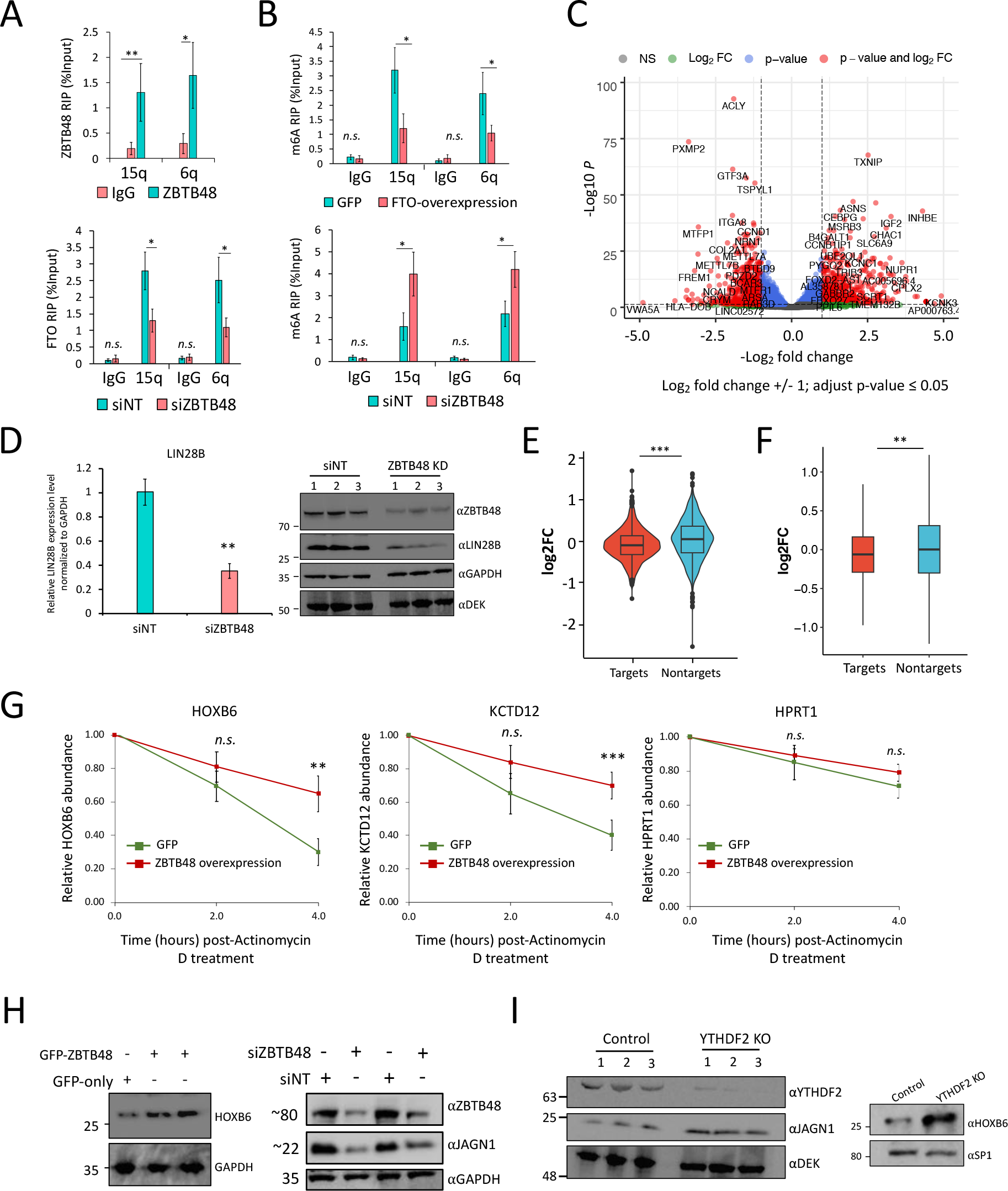
ZBTB48 knockdown alters gene expression and affects mRNA stability. A: Top: RIP-qRT-PCR to examine binding of ZBTB48 to TERRA transcripts in U2OS cells. Bottom: FTO RIP-qPCR bar graph showing effects of siNT- or siZBTB48-treatment on binding of FTO to TERRA in U2OS cells. TERRA was detected using specific primers against the 15q and 6q chromosomes. **B:** Bar plots showing the m6A levels for TERRA transcripts (15q and 6q) in U2OS cells estimated using m6A-RIP-qPCR in GFP- and FTO-overexpressing cells (top), or ZBTB48 knockdown and siNT-treated cells (bottom). Data are represented as % input. **C:** Volcano plot representation of genes differentially expressed in ZBTB48 knockdown HEK293 cells in comparison with control siNT-treated cells. Each dot represents a single gene. Vertical dotted lines represent log2fold change of 1, and highly significant genes are shown as red dots with labels indicating some of the gene names. **D:** left, qRT-PCR results to examine the differential expression of LIN28B in ZBTB48 knockdown cells. Right, Western blotting analysis in whole cell lysates prepared from either siZBTB48 (siZBTB48 #1 + #2) or siNT-treated cells. Blots were probed with the indicated antibodies. Note: DEK and GAPDH were used as loading controls. **E:** Relative abundance of ZBTB48 mRNA targets (based on iCLIP-seq) and non-targets after ZBTB48 KD in HEK293 cells. Violin plots show mean log2 fold change values for the indicated groups (p ≤ 0.01 ANOVA, post hoc Tukey HSD). **F:** Box plot shows mean log2 fold change values for the abundance of mRNA from m6A genes (targets) and non-m6A genes (non-targets) after ZBTB48 KD in HEK293 cells (p ≤ 0.01 ANOVA, post hoc Tukey HSD; Outliers are not depicted). **G:** qRT-PCR quantification of selected m6A-containing mRNAs targeted by ZBTB48 after treating GFP or GFP-ZBTB48 overexpressing cells with Actinomycin D for the indicated times. HPRT1 serves as negative control. **H:** Western blotting analysis in whole cell lysates prepared from cells treated in the indicated ways. Note: ZBTB48 KD (right) was carried out using two different siRNAs, i.e., siZBTB48 #1 and #2 (the second and fourth lanes, respectively). Blots were probed with the indicated antibodies. **I:** Western blotting analysis in whole cell lysates prepared from either YTHDF2 KO or control cells. Blots were probed with the indicated antibodies. Note: For qPCRs (including RIPs): biological replicates n = 5, student’s t-test, ∗∗∗p ≤ 0.001, ∗∗p ≤ 0.01, ∗p ≤ 0.05, n.s.: non-significant. Error bars represent SEM. See also Figures S9, S10 and Table S9.

### ZBTB48 depletion alters expression levels of m6A-containing transcripts

The effects of ZBTB48 on m6A/m6Am levels suggested that it would alter mRNA expression levels. Therefore, we carried out deep RNA-seq experiments, in biological replicates, and identified 1246 genes that were significantly differentially expressed in siZBTB48-treated cells in comparison with control knockdown cells (log2fold change ± 1, p-value (adjusted) ≤ 0.05) (Figure S9A; Table S9). Previously reported downregulated genes [40], including MTFP1, PXMP2, SNX15, and VWA5A, were all identified as significantly downregulated in our RNA-seq analysis (Table S9). Among the significantly differential genes, 645 were upregulated, whereas 601 were identified as significantly downregulated (Figure 5C; Table S9). When we overlapped identified differential genes with those genes whose promoters were targeted by ZBTB48 in our ChIP-seq data, we found that ∼33% and ∼40% of the upregulated and downregulated genes, respectively, overlapped with promoter-bound genes (Figure S9B). Ten out of twelve of the observed gene expression changes that we selected to test were validated by qRT-PCR utilizing RNA derived from independent knockdown experiments using a different ZBTB48 siRNA (siZBTB48 # 2) (Figure S9C). To confirm that ZBTB48 indeed affects gene expression, LIN28B, which was identified as significantly downregulated by RNA-seq analysis, was further examined to assess corresponding changes at the protein level. The levels of both LIN28B mRNA and protein were indeed reduced in cells treated with ZBTB48 siRNAs (siZBTB48 # 2) (Figure 5D; Figure S9D). While some of the observed expression changes might be an indirect consequence of siZBTB48-mediated telomeric length alterations, these observations are consistent with previous findings [40] indicating that ZBTB48 can function as a transcription activator for a small set of genes.

We next assessed whether RNA-binding has a role in ZBTB48-mediated gene expression changes and observed that ∼50% and ∼34% of the down- and up-regulated genes, respectively, indeed corresponded to transcripts bound by ZBTB48 (Figure S9E, left). Global analysis of gene expression changes revealed that, on average, ZBTB48-bound transcripts had significantly lower expression levels in comparison with non-target transcripts (Figure 5E, p-value 0.0005, ANOVA post hoc Tukey HSD). Subsequently, we examined whether the reduction of ZBTB48 impacts the levels of transcripts containing m6A. Although the overall impact was not large, m6A-containing transcripts had a small, but significant, reduction in their overall abundance in comparison with the non-m6A transcripts in siZBTB48 cells (Figure 5F, p-value 0.005, ANOVA post hoc Tukey HSD test), even though the expression levels of m6A methyltransferase complex subunits, PCIF1, YTH-family proteins, and FTO were not significantly altered in these cells (Figure S9F, also see Figure S7C). Importantly, genes bound by ZBTB48, as detected by ChIP-seq analysis, that produce m6A-modified transcripts did not significantly change in expression in comparison with genes producing transcripts lacking m6A modification (Figure S10A). The observed changes in both directions of m6A-containing mRNA abundance might be explained by the binding of different m6A reader proteins that either stabilize or destabilize similar numbers of target transcripts. For example, YTHDF2 has been reported to predominantly destabilize its targets [21], whereas IGF2BP2 stabilizes m6A-marked transcripts [19].

Since many of the promoter-bound genes are also bound by ZBTB48 at the transcript level (Figure S9E, right), the above-described analysis might be confounded by the transcription-related functions of ZBTB48. To more directly assess effects of ZBTB48 on mRNA stability, we treated ZBTB48- overexpressing cells with the transcription inhibitor Actinomycin D and used qRT-PCR to measure mRNA levels over time. In comparison with GFP-expressing control cells, HOXB6 and KCTD12 transcripts were significantly stabilized, whereas JUN was relatively destabilized in ZBTB48- overexpressing cells (Figures 5G and S10B). Consistently, HOXB6 and KCTD12 mRNAs are targeted by YTHDF2 and have been reported to exhibit increased mRNA half-life in YTHDF2- depleted cells [21]. JUN mRNA, on the other hand, is bound by IGF2BPs [19].

Given that HOXB6 and JAGN1 mRNAs were recognized as targets of ZBTB48 and harbor m6A modification, along with decreased expression in siZBTB48-treated cells (Figure S9C), we evaluated the potential influence of ZBTB48 on their protein levels. Of note, transcripts for both HOXB6 and JAGN1 are targeted by YTHDF2 [21]. In line with our RNA-seq data, HOXB6 protein levels were increased upon ZBTB48 overexpression in comparison with GFP cells (Figure 5H, left). FTO overexpression also led to an increased HOXB6 protein levels (Figure S10C). JAGN1 protein and mRNA levels, on the other hand, were reduced when ZBTB48 was knocked down using two different siRNAs (Figures 5H and S10D). Consistent with their functional interaction, FTO knockdown also decreased JAGN1 protein levels (Figure S10E). We then depleted YTHDF2 using CRISPR-Cas9 mediated gene editing and similarly examined HOXB1 and JAGN1 mRNA and protein levels (Figure S10F). We confirmed binding of YTHDF2 to JAGN1 and HOXB6 transcripts by RIP-qPCR (Figure S10G). Consistently, YTHDF2 depletion substantially elevated JAGN1 and HOXB6 expression at both the transcript and protein levels (Figures 5I and S10H). Although it remains possible that ZBTB48 might regulate target mRNA expression through additional mechanisms, the results presented above are consistent with the idea that ZBTB48 recruits FTO to certain m6A-containing target mRNAs, which would then potentially result in loss of m6A and reader-dependent regulation.

### ZBTB48 inhibits colorectal cancer cell growth

Recent studies have linked FTO-mediated m6A/m6Am demethylation to modulation of the stem-like properties of colorectal cancer (CRC) cells, as well as to CRC metastasis [27,53]. Consistent with that, we found that ZBTB48 depletion significantly increases the proliferation of HEK293 cells, whereas its overexpression results in growth inhibition (Figure 6A, p ≤ 0.01, two-way ANOVA). We then utilized the tumor-derived CRC cell line HCT-116 and examined the impact of FTO or ZBTB48 depletion on cell proliferation. Consistent with previous findings [53], knockdown of FTO significantly accelerated the proliferation of HCT-116 cells (Figure 6B; Figure S11A), and we found that knockdown of ZBTB48 also significantly increased the proliferation of HCT-116 cells (Figure 6B; Figure S11A). When we used ‘The Cancer Genome Atlas’ (TCGA), the ‘Genotype-Tissue Expression’ (GTEx) dataset, and the ‘Clinical Proteomic Tumor Analysis Consortium’ (CPTAC) Data Portal to compare transcriptome and proteome expression between tumor and normal tissues, we found that both ZBTB48 and FTO had lower expression levels in colon adenocarcinoma (COAD) tumor tissues in comparison with normal tissues (Figure S11B). These observations are consistent with the idea that ZBTB48 and FTO might function to supress cellular proliferation in CRC. Since the expression of the key m6A writer, eraser, and reader genes did not significantly change in siZBTB48 cells (Figure S11C), we explored the possibility that the observed increase in cell proliferation could be related to ZBTB48-mediated m6A/m6Am modulation. Remarkably, double knockdown of METTL3 and ZBTB48 (Figure S11D) rescued the observed siZBTB48- mediated cell proliferation phenotype (Figure 6B), suggesting that the siZBTB48-associated enhanced growth phenotype is related, at least partially, to its role (s) in m6A regulation. Although telomere length has been reported not to be significantly altered in ZBTB48-depleted HCT-116 cells [54], we cannot completely rule out the possibility that the observed cellular proliferation phenotype might be partly related to indirect gene expression changes due to transcription-related functions of ZBTB48.

**Figure 6:**
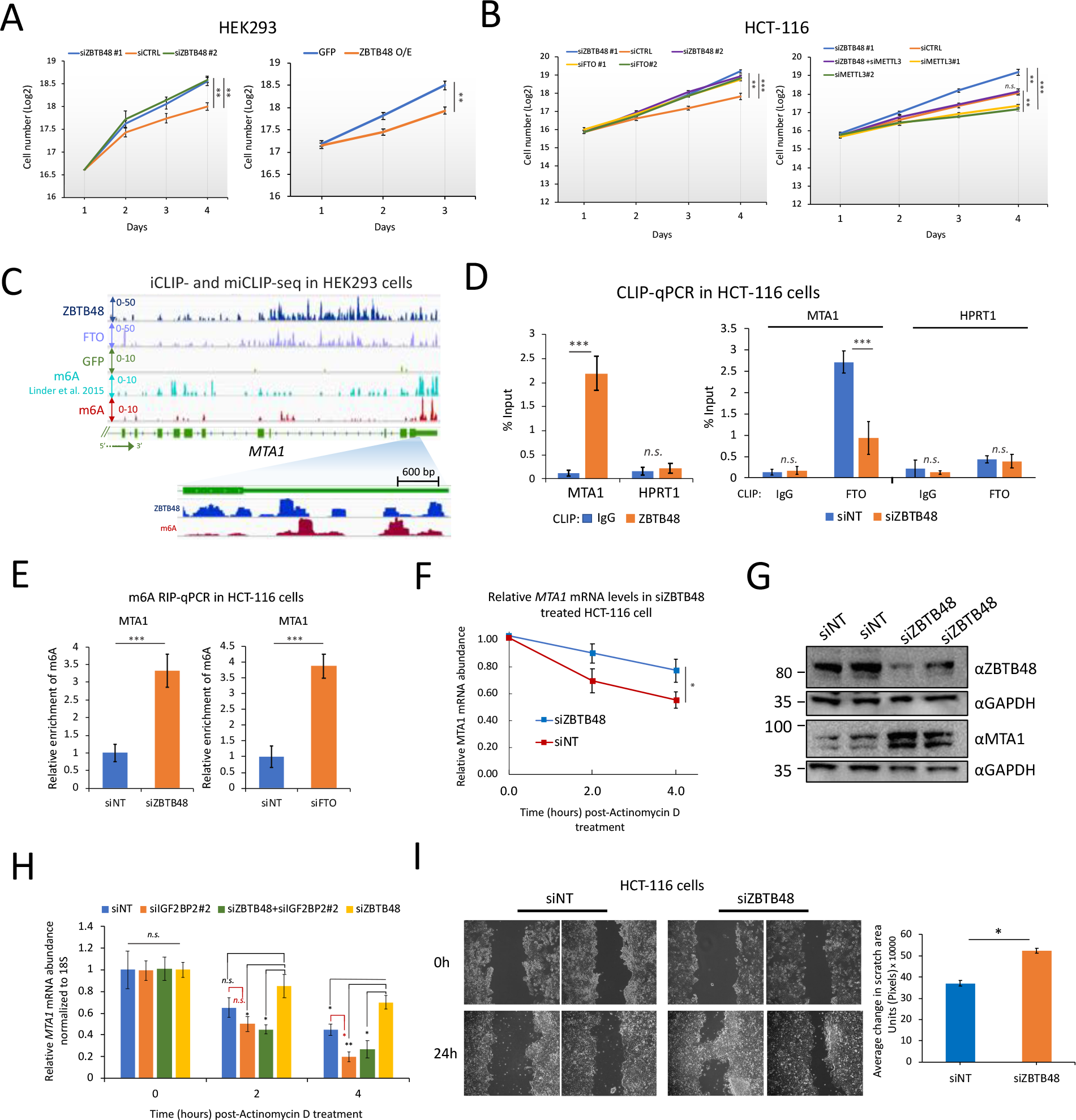
ZBTB48 inhibits cellular proliferation. A: Effects of ZBTB48 KD (left) or overexpression (right) on cell proliferation in HEK293 cells (error bars denote SEM, n = 4, ∗∗p ≤ 0.01, ∗p ≤ 0.05, two-way ANOVA). Cell counts are presented as log2. **B:** Equal numbers of HCT- 116 cells were transfected with the indicated siRNAs in biological triplicates and seeded at the same time in 6-well plates. Cells were separately transfected and seeded for each time point in biological triplicates. Error bars denote SEM. ∗∗∗p ≤ 0.001, ∗∗p ≤ 0.01, ∗p ≤ 0.05 (two-way ANOVA). **C:** Representative genome browser snapshot for MTA1 locus showing the read coverage for ZBTB48 iCLIP-seq, miCLIP-seq (GFP samples), and previously published miCLIP-seq data in HEK293 cells. **D:** Left: CLIP-qRT-PCR to examine binding of ZBTB48 to MTA1 transcripts in HCT-116 cells. Control IPs were performed using IgG. HPRT1 was used as a negative control. Right: CLIP-qPCR to examine binding of FTO to MTA1 transcripts in siNT or siZBTB48 HCT-116 cells. Data are represented as % input. **E:** Bar plot showing the relative m6A levels for MTA1 transcripts estimated using m6A-RIP-qPCR in ZBTB48 knockdown (left), siFTO (right), or siNT-treated HCT-116 cells. Data are represented as % input. **F:** qRT-PCR analysis of MTA1 mRNA in siZBTB48 or siNT-treated HCT-116 cells after treating cells with Actinomycin D for the indicated times. **G:** Western blotting analysis in whole cell lysates prepared from either ZBTB48 KD or control cells. ZBTB48 KD was carried out using two different siRNAs, i.e., siZBTB48 #1 and #2 (third and fourth lanes, respectively). Blots were probed with the indicated antibodies. **H:** qRT-PCR analysis of MTA1 mRNA in cells treated with siNT, siZBTB48 (siRNA#2), siIGF2BP2 (siRNA#2), or a combination of siZBTB48 and siIGF2BP2. For each time point, relative transcript levels were normalized to 18S rRNA and time point “0”. **I:** Wound healing assay using ZBTB48-knockdown or control cells was recorded and quantitatively analyzed (right) (∗p ≤ 0.05, two-way ANOVA). Note: For all qPCR experiments: at least 3 biological replicates, student’s t-test, ∗∗∗p ≤ 0.001, ∗∗p ≤ 0.01, ∗p ≤ 0.05, n.s.: non-significant, error bars denote SEM. See also Figures S11 and S12.

To further delineate the underlying mechanism through which ZBTB48 might influence CRC progression, we examined ZBTB48-bound mRNAs and identified the metastasis-related gene *MTA1* as one of its targets in HEK293 cells (Figure 6C). We prioritized MTA1 for further analysis because: 1) MTA1 has known tumorigenic properties [55], 2) MTA1 mRNA has been shown to be regulated by FTO in an m6A-dependent manner in CRC cells [53], and 3) the MTA1 promoter is not targeted by ZBTB48 in HEK293 cells, thus reducing the possibility of indirect effects related to the function of ZBTB48 as a transcription factor. Since the MTA1 mRNA appears to contain m6A and is targeted by ZBTB48 in HEK293 cells (Figure 6C), we examined whether ZBTB48 could modulate m6A levels in MTA1 mRNA through FTO in HCT-116 cells. First, using co-IPs, we confirmed that FTO interacts directly or indirectly with ZBTB48 in HCT-116 cells (Figure S11E). Consistent with our findings in HEK293 cells, FTO CLIP-qPCR experiments showed significantly reduced signal for MTA1 transcripts in HCT-116 cells treated with siZBTB48, in comparison with control siNT cells (Figure 6D), suggesting that ZBTB48 facilitates targeting of FTO to MTA1 transcripts in CRC cells. When we isolated RNA from ZBTB48-depleted HCT-116 cells and performed m6A-RIP-qPCR experiments, we also observed significantly increased m6A levels in MTA1 mRNA in siZBTB48- treated cells (Figure 6E). FTO knockdown also significantly increased m6A in MTA1 transcripts (Figure 6E, p ≤ 0.001; t-test). These results suggest that, in HCT-116 cells, ZBTB48 modulates m6A in MTA1 mRNA by coordinating the targeting of FTO, consistent with our findings in HEK293 cells.

To assess the downstream effect of ZBTB48 depletion on MTA1 mRNA stability, we performed qRT-PCR experiments after blocking pre-mRNA synthesis by treating HCT-116 cells with Actinomycin D. In comparison with control knockdowns, MTA1 transcripts were significantly stabilized in ZBTB48-depleted cells (Figure 6F). Consistent with these results, MTA1 protein levels were significantly elevated in ZBTB48-depleted cells (Figures 6G and S11F). As reported previously [53], FTO knockdown also enhanced the stability of MTA1 mRNA (Figure S11G). IGF2BP2 is highly expressed in COAD tumor tissues and, among m6A reader proteins implicated in mRNA stability, only IGF2BP2 exhibits a significant positive correlation with MTA1 expression in COAD (Figures S12A and S12B) [53]. Since MTA1 mRNA has been shown to be stabilized by the direct binding of IGF2BP2 in CRC cells [53], we performed RIP-qPCR experiments in HCT-116 cells and observed that IGF2BP2 is bound to MTA1 mRNA and that this binding was significantly increased in ZBTB48-depleted cells, whereas METTL3 knockdown significantly reduced this binding (Figure S12C). Consistently, since IGF2BP2 stabilizes MTA1 mRNA in an m6A-dependent manner, we found that the observed increased stability of MTA1 in ZBTB48 knockdown cells could be reversed by the simultaneous depletion of IGF2BP2 (Figures 6H, S12D, and S12E). We then performed wound healing assays that probe cell migration *in vitro*, and we found that ZBTB48 depletion significantly enhanced wound healing efficiency in comparison with control knockdowns (Figure 6I), consistent with the idea that ZBTB48 inhibits cell migration and proliferation. We conclude that ZBTB48 expression inhibits proliferation of CRC-derived HCT-116 cells, at least partially by regulating the targeting of FTO to certain m6A-containing mRNAs, thus indirectly modulating m6A-reader-dependent mRNA stability.

## Discussion

In this study, we uncovered a previously unknown function of telomere-associated zinc finger protein ZBTB48 in modulating FTO-mediated m6A/m6Am demethylation of mRNAs. We showed that ZBTB48 binds directly to RNA in cultured cells and modulates cellular m6A dynamics through its direct or indirect physical interaction with FTO. Our results suggest that ZBTB48 is important for recruiting FTO to target RNAs and hence modulates cellular m6A/m6Am levels, which impacts mRNA metabolism via the binding of specific reader proteins (Figure 7).

**Figure 7:**
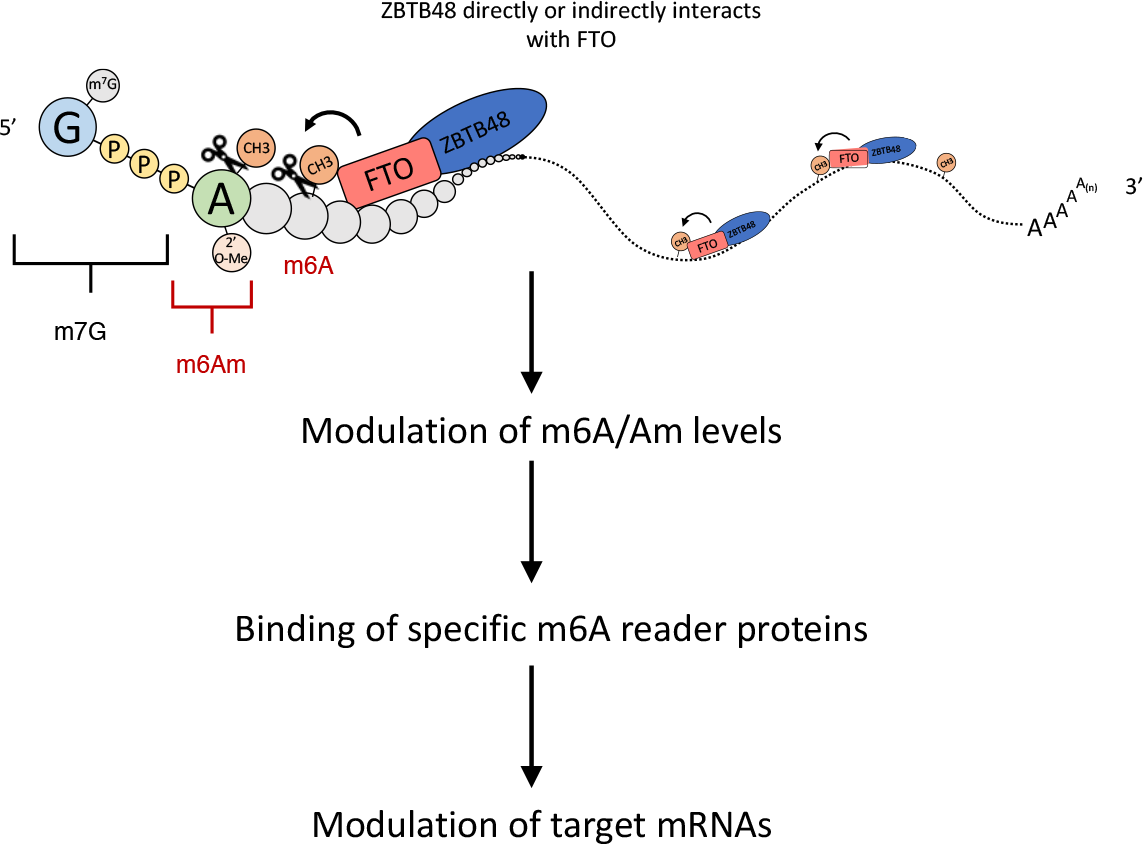
Proposed model for ZBTB48-mediated regulation of m6A/m6Am. ZBTB48 binds to RNA in the vicinity of m6A/m6Am and recruits FTO through either a direct or indirect interaction with it.

FTO can catalyze both m6A and m6Am demethylation of mRNAs [25,32]. How FTO achieves its substrate selectivity is likely complex, and multiple factors, including the sequence and structure of RNA, as well as the subcellular localization of FTO and its protein-protein interactions, have been suggested to play a role in determining FTO’s substrate preferences [34,56,57]. Although recent proteomic studies have attempted to identify the interaction partners of FTO using AP-MS and/or proximity-based Bio-ID approaches, these studies have had limited success. For example, AP-MS analyses of exogenously expressed FTO did not identify any significant interaction partners [57], whereas Bio-ID recovered several highly abundant proteins, including ubiquitous transcription factors and DNA replication/repair proteins [58]. Our results support the idea that the *in vivo* RNA-binding of FTO might be facilitated, at least in part, by protein-protein interactions. We have demonstrated that ZBTB48 facilitates the targeting of FTO to both m6A and m6Am in a transcriptome-wide manner. Considering that FTO can localize to the cytoplasm in certain cell-types [34], it is conceivable that additional tissue and/or context-specific factors, yet to be identified, might also function to facilitate its targeting to cytoplasmic m6A and/or m6Am sites. Although we have shown that ZBTB48 interacts physically with FTO, either directly, or else indirectly in a complex with other proteins, and facilitates its targeting to RNA, we did not investigate the stoichiometry, the strength, and or the structural context of their interaction. Although it is the region containing the zinc fingers of ZBTB48 that interacts with RNA, it is not this region, and may be the BTB domain of ZBTB48, that mediates its interaction with FTO.

It was previously shown that neither FTO nor ALKBH5 exhibits selectivity towards m6A DRACH motifs *in vitro* [36]. Although an m6Am-specific BCA-like motif was identified in our ZBTB48 iCLIP-seq peaks, which also circumstantially supports the recruitment of ZBTB48 to m6Am, our results revealed U-rich sequences to also be significantly overrepresented in ZBTB48 crosslink sites in mRNA. In line with this observation and the interaction of FTO with ZBTB48, previous studies have reported the enrichment of pyrimidine-rich sequences in FTO CLIP-seq experiments [37]. There is also some precedent in the literature for U-rich sequences in the context of m6A modulation. For example, U-rich sequences have been reported to be enriched around m6A sites on RNA [59]. RBM15/RBM15B, two components of the m6A-methylation complex, recognize these U-rich sequences in the vicinity of m6A sites on individual transcripts and recruit the Mettl3.Mettl14- complex [59]. RBM15 and RBM15B both exhibit RNA-binding profiles similar to that of ZBTB48 (i.e., more enriched in 5’UTRs in comparison with 3’UTRs) [59]. Moreover, DDX46 recruits ALKBH5 to certain m6A sites by recognizing the sequence CCGGUU [60]. These studies are consistent with the idea that sequence (or structure)-specific RBPs can modulate m6A/m6Am levels in cells by interacting with and recruiting methyltransferase or demethylase enzymes to specific sites on RNA. Based on our results, we suggest that ZBTB48 binds certain U-rich sequences in the vicinity of m6A sites in target mRNAs and facilitates the recruitment of FTO to these sites. The sequence/structural context of ZBTB48 binding to RNA remains to be examined, however. While our manuscript was in preparation, a recent study demonstrated that RBM33 regulates ALKBH5- mediated m6A demethylation of specific transcripts through its physical interaction with ALKBH5 [61].

Some studies have previously shown that FTO depletion causes only a mild (∼10-20%), albeit significant, increase in global m6A levels in PolyA mRNAs. Although FTO overexpression (or depletion) resulted in ∼2-3-fold change in global m6A levels in total cellular RNAs, we also found that FTO-overexpression causes only a mild decrease in m6A levels in PolyA transcripts, consistent with previous studies. Intriguingly, even though ZBTB48 appears to regulate FTO recruitment to only a subset of m6A targets, we found that the change in global m6A levels upon ZBTB48 overexpression is comparable (or even slightly more) to that of FTO-overexpression, presumably because the m6A signal in miCLIP experiments was dominated by their co-regulatory sites. Consistent with this notion, knockdown of RBM33, which regulates ALKBH5 recruitment to a subset of m6A sites, was found to change global m6A levels similarly to ALKBH5-depletion. Moreover, m6A changes related to indirect effects in ZBTB48 KD (or overexpression) cells remain another possibility. Although FTO expression remained unchanged after knockdown of ZBTB48, we did observe an increase in FTO levels in ZBTB48-overexpressing cells. Considering that ZBTB48 does not appear to target the FTO promoter, it is likely that the observed increase in FTO expression is an indirect effect. FTO is known to negatively regulate its own expression through an auto-regulatory loop [62]. It is conceivable that an increased number of ZBTB48 molecules could sequester FTO and thus disrupt the binding of FTO to a negative regulatory site, which could, in turn, lead to an upregulation of the production of FTO.

FTO has been suggested to function as both an oncogene and a tumor suppressor, depending on the cancer type [63]. We have found that ZBTB48 expression inhibits the growth of colorectal cancer-derived HCT-116 cells. Considering the multifunctional nature of ZBTB48, the underlying mechanism of ZBTB48-mediated growth inhibition and/or its potential role as a tumor suppressor is likely complex. For example, we found that ZBTB48 can regulate gene expression by both acting as a transcription factor and modulating levels of m6A/m6Am, with the latter controlling the expression of the oncoprotein MTA1. As well, telomere length is well known to play a crucial role in cancer cell survival and, recently, m6A was implicated in the regulation of telomere metabolism by downregulating HMBOX1, a telomeric DNA-binding protein [64]. Moreover, telomeric repeat-containing RNA, TERRA, which forms R-loops with telomeric DNA and also functions in telomeric stability, has been shown to be regulated by METTL3-mediated m6A[11,51]. We found that, just as ZBTB48 can modulate mRNA m6A levels by recruiting FTO, it can also modulate TERRA m6A levels by recruiting FTO. ZBTB48 was previously shown to bind telomeric repeat DNA and localize to telomeres [40,41] and, although we found that TERRA is co-immunoprecipitated with ZBTB48, it remains to be determined whether this association occurs in the context of ZBTB48 binding to TERRA, R-loops containing TERRA, or telomeric double-stranded DNA. Considering the known roles of ZBTB48 and TERRA m6A modifications in telomere length regulation, it will also be interesting to explore whether ZBTB48-mediated m6A-modulation of TERRA mediates its impact on telomere homeostasis. Moreover, recent work indicates that FTO mediates demethylation of repeat elements [65]. It will be interesting to explore the role of ZBTB48 in this context.

Finally, although we showed the interaction between ZBTB48 and FTO in HEK293 and HCT-116 cells, we did not examine whether these proteins also interact in other cell types. Some of the observed m6A changes might be indirect, since ZBTB48 can also influence gene expression as a transcription factor. As well, although we showed that ZBTB48 can inhibit cell proliferation in a manner that depends on its role in m6A metabolism, future studies will be needed to fully dissect the contributions of the transcriptional and posttranscriptional roles of ZBTB48 in cell growth and tumorigenesis. It will also be important to interrogate whether this effect is maintained *in vivo*.

## Materials and Methods

### Cell culture and cell counts

The HEK293 Flp-In T-REx cell line was obtained from Life Technologies (Invitrogen catalogue number R780-07). HEK293 and HCT-116 cells (isolated from the colon of an adult male) were acquired from ATCC (CCL-247 and CRL1573, respectively). U2OS cells were a kind gift from Dr. Alexander F. Palazzo. Cell cultures were maintained in Dulbecco’s modified Eagle’s medium (DMEM) (Wisent Bioproducts catalogue number 319-005-CL) supplemented with 10% FBS (Wisent Bioproducts catalogue number 080-705), sodium pyruvate, non-essential amino acids, and penicillin/streptomycin as described.

For cell proliferation assays, equal numbers (∼10^5^) of cells were grown in 6 well plates in biological triplicates. For siRNA knockdown experiments, cells were transfected with the indicated siRNAs in biological triplicates and seeded at the same time in 6-well plates. Moreover, cells were separately transfected and seeded for each time point. Cells were extracted using trypsin (Wisent Bioproducts catalogue number 325-043EL) for each time point and counted using the Invitrogen Countess automated Cell Counter hemocytometer (Invitrogen AMQAX1000).

### Epitope tagging in HEK293 cells

A Gateway™-compatible entry clone for the ZBTB48 ORF was cloned into the pDEST pcDNA5/FRT/TO-eGFP vector according to the manufacturer’s instructions. The vector was co-transfected into HEK293 Flp-In T-REx cells with the pOG44 Flp recombinase expression plasmid. Cells were selected with hygromycin (Life Technologies, 10687010) at 200 μg/mL for FRT site-specific recombination into the genome. Expression was induced by adding Doxycycline (1 μg/mL) into the culture medium 24 hours prior to harvesting the cells.

### Immunoprecipitation (IP) and western blots

IPs were performed essentially as described previously [48,66]. Briefly, cells were lysed in 1 mL of lysis buffer containing 140 mM NaCl, 10 mM Tris pH 7.6–8.0, 1% Triton X-100, 0.1% sodium deoxycholate, 1 mM EDTA, and supplemented with protease inhibitors (Roche catalogue number 05892791001). Cell extracts were incubated with Benzonase (75 units) (Sigma E1014) for 30 minutes in a cold room with end-to-end rotation and clarified in a microcentrifuge at 15,000 g for 30 minutes at 4°C. The supernatant was incubated with 10 μg GFP antibody (Life Technologies G10362) for 2 hours with end-to-end rotation at 4°C. ZBTB48 and FLAG pulldown were performed with 10 μg of ZBTB48 (GeneTex GTX118671) and FLAG (Sigma F3165) antibodies, respectively. 10 μg IgG (Santa Cruz Biotechnology, catalog number: sc-2025) was utilized as a control. Lysates were incubated for an additional 2 hours (Invitrogen catalogue number 10003D) with 100 μL protein A/G Dynabeads at 4°C with end-to-end rotation. Beads were washed (3☓) with lysis buffer containing an additional 1% NP40 and 2% Triton X-100. The samples were boiled in SDS sample buffer and resolved using 4–12% BisTris–PAGE. Proteins were transferred to a nitrocellulose membrane (Thermo Scientific™ catalogue number 77010) using a Gel Transfer Cell (BioRad catalogue number 1703930) as per the manufacturer’s instructions. Primary antibodies were used at 1:1000 dilution, and horseradish peroxidase-conjugated goat anti-mouse (Thermo Fisher 31430) or anti-rabbit secondary (Thermo Fisher 31460) antibodies were used at 1:10,000. Blots were developed using Pierce ECL Western Blotting Substrate (Thermo Scientific™ catalogue number 32106).

### iCLIP-seq procedure

Individual nucleotide resolution UV crosslinking and immunoprecipitation (iCLIP) [67] was performed with the modifications reported in iCLIP-1.5, as detailed in our previous reports [48,68]. Briefly, cells were grown in 15 cm culture plates in independent batches. Protein expression was induced with Doxycycline (1μg/mL) for 24 hours, and cells were UV crosslinked using 0.15 J/cm^2^ at 254 nm in a Stratalinker 1800. Cell lysis was performed in 2.2 mL iCLIP lysis buffer, and 1 mL of the lysate was incubated with Turbo DNase (Life Technologies catalogue number AM2238) and RNase I (1:250; Ambion catalogue number AM2294) for exactly 5 minutes at 37°C with shaking (1400 rpm) in a thermomixer to digest genomic DNA and obtain RNA fragments of an optimal size range. A total of 2% input material was separated to generate size-matched control libraries (SMI). ZBTB48, GFP-ZBTB48, and FLAG-FTO were immunoprecipitated using 10 μg of anti-ZBTB48 (GeneTex GTX118671), anti-GFP antibody (Life Technologies G10362), and anti-FLAG (Sigma F3165) antibodies, respectively. Following stringent washes with iCLIP high salt buffer, dephosphorylation was performed using FastAP and T4 polynucleotide kinase. Pre-adenylated L3 adaptors were ligated to the 3’-ends of RNAs using the enhanced CLIP ligation method, as described [69]. The immunoprecipitated RNA was 5’-end-labeled with ^32^P using T4 polynucleotide kinase (New England Biolabs catalogue number M0201L). Protein-RNA complexes were separated using 4–12% BisTris–PAGE and transferred to a nitrocellulose membrane (Protran). SMI samples, were also resolved on 4–12% BisTris–PAGE and transferred to a nitrocellulose membrane alongside the immunoprecipitated RNA. The membrane corresponding to SMI and IP lanes was excised. To recover RNA from the membrane, proteins were digested with proteinase K (Thermo Fisher catalogue number 25530049). RNA was reverse transcribed into cDNA using barcoded iCLIP primers. The cDNA was size-selected (low: 70 to 85 nt, middle: 85 to 110 nt, and high: 110 to 180 nt), and circularized using CircLigase™ II ssDNA ligase to add the adaptor to the 5’-end. In the CircLigase reaction, we added betaine at a final concentration of 1 M, and the reaction mixture was incubated for 2 hours at 60°C. Circularized cDNA was digested at the internal BamHI site for linearization, then PCR amplified using AccuPrime SuperMix I (Thermo Fisher catalog number 12344040) or Phusion High-Fidelity PCR Master Mix (NEB, M0531S). After agarose gel purification using columns (QIAGEN), the eluted PCR libraries were mixed at a ratio of 1:5:5 from the low, middle, and high fractions and submitted for sequencing on an Illumina NextSeq 500 platform using a full High-Output v2.5 flow cell to generate single-end 51 nucleotide reads with 40M read depth per sample. The barcoded primers used for iCLIP-seq are listed in Table S10.

### miCLIP-seq procedure

miCLIP-seq was performed essentially as previously described [28], with minor modifications as detailed below. Briefly, cells were grown in independent batches to represent biological replicates and expression of the epitope-tagged proteins of interest was induced 24 hours prior to harvesting by adding Doxycycline (1 μg/mL) into the culture medium. Total RNA was extracted from either GFP-ZBTB48 or FTO or GFP-only overexpressing HEK293 cells using TRIzol as per manufacturer’s protocol. RNA was treated with DNase I to eliminate any contaminating genomic DNA. We used 12 μg of total RNA for miCLIP-seq. 2 μL of RNA Fragmentation Reagents (Invitrogen AM8740) was added to the RNA suspended in 20 μL of sterile water. The mixture was incubated in a heat block at 75°C for exactly 12 minutes. 2.2 μL of stop solution (Invitrogen AM8740) was added to stop the reaction. The fragmented RNA was incubated with 10 μg of m6A antibody (MBL RN131P) for 2 hours at 4°C. RNA-antibody complexes were UV crosslinked twice with 0.15 J/cm^2^ at 254 nm in a Stratalinker 1800. Protein G Dynabeads (100 μL) were added, and the samples were rotated for 2 hours at 4°C. Samples were washed three times using iCLIP high salt buffer, and RNA dephosphorylation was performed using FastAP and T4 polynucleotide kinase. Pre-adenylated L3 adaptors were ligated to the 3’-ends of RNAs using the enhanced CLIP ligation method, as detailed above. The immunoprecipitated RNA was 5’-end-labeled with ^32^P using T4 polynucleotide kinase (New England Biolabs catalogue number M0201L), separated using 4–12% BisTris–PAGE and transferred to a nitrocellulose membrane (Protran). The RNA isolation, cDNA preparation, cDNA size selection, circularization, and library amplification step were performed essentially as described above for the iCLIP-seq procedure. The final PCR libraries were agarose gel purified on purification columns (QIAGEN), and the eluted DNA was mixed at a ratio of 1:5:5 from the low, middle, and high fractions and submitted for sequencing on an Illumina NextSeq 500 platform using a full High-Output v2.5 flow cell to generate single-end 51 nucleotide reads with 40M read depth per sample. The barcoded primers used for iCLIP-seq are listed in Table S10.

### Chromatin immunoprecipitation and sequencing (ChIP-seq)

We performed chromatin immunoprecipitation essentially as described in our previous publications [45]. Briefly, ∼20 million HEK293 cells were grown in independent batches in 15cm plates and were crosslinked in 1% formaldehyde for 10 min at room temperature (RT) in crosslinking buffer (50 mM HEPES, 100 mM NaCl, 1 mM EDTA, 0.5 mM EGTA, 1% formaldehyde). The reaction was quenched with 0.125 M glycine for 10 min at room temperature. Cells were washed twice with PBS on plates and harvested using a cell scraper in ∼10 mL PBS. Protein G Dynabead slurry (30 uL beads) was washed twice with 1 mL cold block solution (0.5% BSA in PBS). 250 ul of blocking solution and 2 μg of GFP antibody (Abcam ab290) were added, and beads were rotated at 4°C for at least 4 hrs to bind antibody to beads. Cell pellets were lysed in 3 mL LB1 buffer (50 mM Hepes, 140 mM NaCl, 1 mM EDTA, 10% glycerol, 0.5% NP-40, 0.25% TX-100), supplemented with protease inhibitors (Roche catalogue number 05892791001), by pipetting and rotating vertically at 4°C for 10 min. Samples were spun for 5 min at 1350 x g at 4°C (GH-3.8 rotor = 2400 rpm) and supernatant was removed. Pellets were resuspended in 3 mL LB2 buffer (10 mM Tris, 200 mM NaCl, 1 mM EDTA, 0.5 mM EGTA) by pipetting and were rotated vertically at room temperature for 10 min. Samples were spun for 5 min at 1350 x g at 4°C and supernatant was removed. Pellets were resuspended in 1 mL of LB3 buffer (10 mM Tris, 100 mM NaCl, 1 mM EDTA, 0.5 mM EGTA, 0.1% Na-Deoxycholate, 0.5% N-lauroylsarcosine) by pipetting in 15 mL polypropylene tubes. DNA was fragmented using sonication in a BioRuptor (BioRuptor settings: power = high, “on” interval = 30 sec, “off” interval = 30 sec (10 cycles of “on”/ “off”). Samples were spun for 10 min at 16000 x g (max) at 4°C to pellet cellular debris, and supernatant was transferred to a fresh tube. 10% Triton X-100 was added to final concentration of 1%. 20 ul was set aside as input. Dynabeads coupled to antibody were washed 2-3 times with blocking solution and were added to the sonicated lysate. Samples were rotated at 4°C overnight (12-16 hrs) to bind antibody to chromatin. Beads were washed at least 5X in 1 mL cold RIPA buffer (50 mM Hepes, 500 mM LiCl, 1 mM EDTA, 1% NP-40, 0.7% Na-Deoxycholate). Beads were washed once in 1 mL TE + 50 mM NaCl in the cold room (4°C). Supernatant was removed by spinning at 950 x g at 4°C. Elution was carried out in 200 uL elution buffer (50 mM Tris, 10 mM EDTA, 1% SDS) for 15 min at 65°C in a water bath, with brief vortexing every few minutes. Input samples were thawed, and 300 uL elution buffer was added. Both input and eluted chromatin samples were incubated overnight (12-16 hrs) in a 65°C water bath to reverse crosslinks. One volume of TE buffer was added to each sample to dilute the SDS. RNA was digested for 2 hours at 37°C with RNase A (0.2 mg/mL final concentration). Subsequently, samples were incubated with proteinase K (0.2 mg/mL final concentration) at 55°C for 2 hours to digest proteins. DNA was purified with a DNA purification kit (QIAGEN), and concentration was measured by Qubit. Libraries were prepared using an Illumina TruSeq ChIP Library Preparation Kit as per the manufacturer’s instructions. Libraries were sequenced on an Illumina HiSeq 2500 to a depth of 20 million paired end reads.

### RNA over-digestion assay

The RNA over-digestion assay follows the initial steps of iCLIP, essentially as described above, except that the 3’-end dephosphorylation and L3 adapter ligation steps were omitted. Following IPs and high-salt washes, RNA was 5’-end-labeled with ^32^P using T4 polynucleotide kinase. Beads were divided into two equal halves and incubated with 5 μL of either RNase I or Turbo DNase for 15 mins while shaking (1200 rpm) at 37 °C. Beads were washed once with 900 μL of PNK buffer. Protein-RNA complexes were separated using 4–12% BisTris–PAGE and transferred to a nitrocellulose membrane (Protran), followed by autoradiography.

### siRNA treatment, Actinomycin D treatment, RNA isolation, and qRT-PCR

siRNA knockdowns were performed with 40-50 nM Silencer Select siRNAs with Lipofectamine iMax transfection reagent (Thermo Fisher 13778075) for 72 hours. Silencer Select control siRNA (Thermo Fisher AM4611) and siRNAs against human ZBTB48, FTO, METTL3, and IGF2BP2 were purchased from Thermo Scientific. The siRNA IDs are as follows: ZBTB48: s6568 (siRNA#1) and s6566 (siRNA#1); IGF2BP2: s20922 and s20923; FTO: s35510 and s35511; and METTL3: s32141 and s32143.

Total RNA was extracted using Trizol Reagent (Invitrogen catalogue number 15596018) as per the manufacturer’s instructions. RNA was reverse transcribed into cDNA using either the SuperScript VILO Kit (Invitrogen catalogue number 11754) or Maxima H Minus First Strand cDNA synthesis Kit (Thermo Scientific K1671). TERRA-specific primer used in the RT reaction was 5’-CCCTAACCCTAACCCTAACCCTAACCCTAA-3’. PCR was performed using the AccuPrime Pfx DNA Polymerase (Thermo Scientific catalogue number 12344024), and qPCR was performed with Fast SYBR Green Master qPCR mix (Applied Bioscience 4385617) on an Applied Biosystems 7300 real time PCR System (Thermo Fisher catalogue number 4406984). The qPCR program involved 40 cycles of 95 °C for 15 s and 60 °C for 30 s, and a final cycle (95 °C for 15 s and then 60 °C). Each biological replicate was subjected to qPCR in 3 technical replicates. All experiments were performed in at least 3 biological replicates.

Actinomycin D treatment was carried out with 5 μg/mL Actinomycin D (Sigma-Aldrich catalogue number A1410) for the indicated times prior to RNA extraction. 18S rRNA was used as loading control for the Actinomycin D RT-qPCR. For each time point, relative transcript levels were normalized to 18S rRNA and time point “0”, using the ΔΔCT method [70]. This method uses the threshold cycles (CTs) generated by the qPCR system for calculation of gene expression levels. Primers are listed in Table S11.

### CRISPR-mediated gene knockout

CRISPR-mediated gene knockouts were performed using the LentiCRISPRv2 plasmid, which was kindly provided by the Moffat Lab (Addgene plasmid #52961). Guide RNA sequences from the Toronto Knockout version 3 (TKOv3) library were used to target either AAVS1 (#1 GGGGCCACTAGGGACAGGAT, #2 GTCACCAATCCTGTCCCTAG) or YTHDF2 (#1 AATATAGGTCAGCCAACCCA, #2 ATATAGGTCAGCCAACCCAG, #3 TATGACCGAACCCACTGCCA). Each guide RNA was cloned into the LentiCRISPRv2 vector and transfected into HEK293T cells to produce lentiviral particles. To test knockout efficiency of individual guide RNAs, HEK293 cells were infected with each of the three lentiviruses targeting YTHDF2 or with pooled lentivirus targeting AAVS1 at an MOI of 0.3-0.4 in 6-well plates. 24 hours after infection, cells were subjected to puromycin selection (final concentration 2 ug/mL; BioShop PUR333.25) for 70 hours, and subsequently grown for 3 days before being harvested for Western blot. Lentiviruses for two YTHDF2 guide RNAs (#1 and #2) were pooled together for all subsequent experiments.

### RNA sequencing

Total RNA was extracted from HEK293 cells treated with siRNAs (either siZBTB48 or non-targeting scrambled control) for 72 hours, as described above. 1 μg purified total RNA in biological duplicates was submitted to the Donnelly Sequencing Centre for further processing. Briefly, Poly- A selected, rRNA-depleted libraries were generated using the TruSeq Illumina library preparation kit (Epicenter) as per the manufacturer’s recommendations. The quantified pool was hybridized at a final concentration of 2.21 pM and sequenced using the Illumina NovaSeq platform, 300 cycles (paired-end sequencing) to a target depth of 50 M reads. RNA-seq data have been deposited in GEO with the accession code GSE228608.

### RNA immunoprecipitation (RIP)

RIP experiments were performed as described previously [71]. Briefly, cells grown in 10-cm plates (∼ 80 % confluent) were harvested and lysed in 1 mL lysis buffer containing 25 mM Tris HCl pH 7.5, 150 mM NaCl, 0.5% (v/v) NP-40, 1 mM AEBSF and 1 mM DTT, supplemented with RNase and protease/phosphatase inhibitors. Lysates were clarified by centrifugation (30 min, 20000 g, 4°C). We kept 5% of the lysate as input material. Immunoprecipitation was performed using 5 μg of antibody (either anti-ZBTB48, or anti-GFP antibody (Life Technologies G10362), or anti-FLAG) conjugated with Protein G Dynabeads (Invitrogen). 5 μg of IgG (Santa Cruz Biotechnology, catalog number: sc-2025) was used in control IPs. Cell lysates were incubated with antibodies for 2–3 hours at 4°C with end-to-end rotation. The beads were washed three times with wash buffer containing 25 mM Tris HCl pH 7.5, 150 mM NaCl, 0.05% (v/v) NP-40, 1 mM AEBSF and 1 mM DTT (5 minutes rotation at 4°C for each wash). RNA from the input and IP samples was extracted using Trizol Reagent (Invitrogen catalogue number 15596018) according to the manufacturer’s instructions. cDNA was synthesized using the First-strand cDNA synthesis kit (Thermo Fisher Scientific) with random hexameric primers.

### CLIP-qRT-PCR

CLIP was performed following the initial steps of iCLIP, essentially as described above. Lysates were treated with 2 μL Turbo DNase and 10 μL RNase I (1:250) for 5 min at 37°C while shaking in a thermomixer (1200 rpm). RNA-protein complexes were immunoprecipitated using 5 μg antibody (either anti-ZBTB48, or anti-GFP antibody (Life Technologies G10362), or anti-FLAG) conjugated with 50 μL Protein A/G Dynabeads. 5 μg IgG (Santa Cruz Biotechnology, catalog number: sc-2025) was used as a control. Following washes (5☓) with CLIP high salt buffer, proteins were digested with proteinase K and RNA extracted using Trizol. The isolated RNA was subjected to qRT-PCR analysis. Input RNA was reverse transcribed into cDNA and used to calculate the percent enrichment in the immunoprecipitated samples. qPCR primers are listed in Table S11.

### m6A dot blot assay

Total RNA was isolated from cells using Trizol (Invitrogen, 15596-018) according to the manufacturer’s instructions. Two rounds of Dynabeads® mRNA DIRECT™ kit (Thermofisher Scientific, #61006) were used to purify mRNA. The m6A-dot-blot was performed following previously published procedures [72]. Briefly, total RNA or mRNA was crosslinked to a charged nylon-based membrane using UV light and was incubated with primary rabbit anti-m6A antibody (Synaptic Systems, 202003) for 1 hour at room temperature. After washing three times with TBE, the membrane was incubated with HRP-conjugated Goat anti-rabbit IgG (DakoCytomation, p0448) for another 1 hour. After washing three times with TBE, blots were developed with enhanced chemiluminescence (GE Healthcare, RPN2232). The signal densities for m6A and blotted mRNA (Methylene blue staining) were quantified by ImageJ software (Media Cybernetics). m6A signal values were normalized to the Methylene blue values (input RNA), and the data were represented in bar graphs with standard error of mean (n=3). For CLIP-m6A-dot blots, the initial steps of the iCLIP were performed as described above, and the isolated RNA was subjected to the m6A dot bot procedure.

### m6A-qRT-PCR

Total RNA was purified using Trizol (Invitrogen, 15596-018) according to the manufacturer’s instructions. RNA was fragmented to ∼300 nt by RNA Fragmentation Reagents, and immunoprecipitation was performed using anti-m6A antibody (Synaptic Systems, 202003), essentially as described above. The enrichment of m6A was measured using quantitative Reverse Transcription Polymerase Chain Reaction (qRT-PCR). Primers for m6A-qRT-PCR are listed in Supplemental Table S11.

### Nucleoside mass-spectrometry analysis

Nucleoside mass-spectrometry (MS) analysis was performed following published protocols [25,32]. Briefly, total RNA or polyadenylated RNA was treated with DNase to ensure the removal of DNA. RNA was cleaned with an RNA clean up kit (Zymo kit, R1013, Cedarlane) and resuspended in 20 μl H2O. RNA concentration was measured, and 500 to 1000 ng RNA was used for digestion. mRNA was de-capped with 0.5 U Cap-Clip enzyme (Cell Script, C-CC15011H) for one hour at 37°C. Total RNA and mRNA samples were further digested with 2 U of Nuclease P1 (N8630, Sigma) overnight at 42 °C in NH4OAc buffer (10 mM, pH 5.3). 1 U of Alkaline phosphatase (Sigma, P4252) and 0.05M MES was added to the reaction and the reaction was incubated for 6 hours at 55 °C. The samples were filtered through 10,000 MW cut off spin filter (AMicon, UFC501024) for 10 min at 16,000 g at 4C. The final filtrate was injected into the MS.

A reverse phase micro-capillary liquid trap column and analytical column were used for molecular separation using the EASY-nLC 1200 system. A 25mm x 75 μm silica micro-capillary trap column was packed with 2.5μm Synergy Hydro RP (Phenomenex). A tip for a 100mm × 75μm silica micro-capillary analytical column was created with a column puller (Sutter Instruments) and packed with 2.5 μm Synergy Hydro RP stationary phase (Phenomenex). The EASY-nLC system was used to drive a 30 min organic gradient using buffers A (0.1% formic acid) and B (80% acetonitrile with 0.1% formic acid). The molecules were separated at a flow rate of 250 nL/min (gradient buffer B: 0% to 100% - 2 min, 100% - 23 minutes, 100% to 0% - 1 min, 0% - 2 min). Eluates were sprayed into an Q Exactive HF mass spectrometer (ThermoFisher Scientific). The full scan was performed in 60000 resolution with 3E6 AGC.

The nucleosides were quantified by using the nucleoside-to-base ion mass transitions of 282.1 to 150.1 (m6A), 268 to 136 (A), 296 to 150 (m6Am), and 282 to 136 (Am). The m6A and m6Am were quantified as the ratio of m6A and m6Am to A, respectively.

### Wound healing assay

The wound healing assay was performed essentially as previously described [73]. Horizontal lines were drawn using a marker pen on the back of 6-well plates. Cells were grown in these 6-well plates to an optimal density and a pipette tip was used to scratch a wound through the centre of the well, perpendicular to the marked horizontal lines. Cells were washed with PBS to remove cellular debris and serum-free media was added. Wound healing was observed after 24 hours with an inverted microscope (OLYMPUS CKX 41). We used *Wound Healing Size Tool*, an ImageJ/Fiji® plugin, to calculate the healing area of scratches [74].

### Immunofluorescence

HEK293 cells were seeded on poly-L-lysine-coated and acid-washed coverslips. Cells were washed three times with PBS. Cells were fixed in 4% Paraformaldeyde for 15 minutes, permeabilized with 0.2% Triton X-100 in PBS for 5 minutes, and incubated with block solution (1% goat serum, 1% BSA, 0.5% Tween-20 in PBS) for 1 hour. ZBTB48 antibody (Invitrogen PA5- 56467) or FTO antibody (Santa Cruz Biotechnology, catalog number: sc-271713) was used for staining at 1:100 concentration in block solution for 2 hours at room temperature (RT). Cells were incubated with Goat anti-rabbit (or anti-mouse for FTO) secondary antibody and Hoescht stain in block solution for 1 hour at room temperature. Cells were fixed in Dako Fluorescence Mounting Medium (S3023). Imaging was performed using a Zeiss confocal spinning disc AxioObserverZ1 microscope equipped with an Axiocam 506 camera using Zen software.

### Quantification and statistical analysis iCLIP-seq analysis

iCLIP- and miCLIP-seq analyses were performed as detailed in our previous reports [48,75]. 51- nt iCLIP-seq and miCLIP-seq raw reads consist of 3 random positions, a 4-nt multiplexing barcode, and another 2 random positions, followed by the cDNA sequence. Reads were de- duplicated based on the first 45 nt. After de-multiplexing, we removed the random positions, barcodes, and any 3′-bases matching Illumina adaptors. Reads shorter than 25nt were discarded, and the remaining reads were trimmed to 35nt. These steps were carried out using Trimmomatic [76]. Reads were mapped to the human genome/transcriptome (Ensembl annotation hg19) using Tophat [77] with default settings. Reads with a mapping quality < 3 were removed from further analysis, which also removes multi-mapping reads. Data for iCLIP-seq analyses have been deposited in GEO with the accession code GSE228608.

### Crosslink induced truncations (CITS)

CLIP Tool Kit (CTK) was used to identify RNA-binding sites at single-nucleotide resolution [78]. iCLIP-seq primarily captures the crosslink-induced truncation sites (CITS) during cDNA preparation. We thus called CITS peaks on individual CLIP replicates as well as SMI samples. CITS with FDR ≤ 0.01 were considered as significant. The identified significant CITS were utilized to extract underlying reads from each dataset and the resulting CLIP reads were normalized to SMI reads (iCLIP/SMI). Moreover, we filtered out any CITS that were present in GFP-only iCLIP-seq samples [49]. Remaining CITS peaks from individual replicates were merged for downstream analysis. miCLIP-seq data were processed similarly to the iCLIP-seq data, and putative anti-m6A mediated CITS were identified (FDR ≤ 0.01). Subsequently, m6A/m6Am sites were identified using the following criteria: m6A CITS from 2 replicates (p-value 0.01) were merged at a maximum gap of 1 bp. Each peak was extended 2 bp both upstream and downstream of the identified CITS, and from these extended peaks, only those with a DRACH motif were selected. The coordinates of the base ‘A’ in the motif were designated as m6A sites. For m6Am sites, each peak was extended 2 bp both upstream and downstream, and only the extended peaks with at least one base ‘A’ were selected, with consecutive As merged, and the coordinates of A were recorded, producing m6Am candidate sites. The m6Am candidate sites were first filtered by selecting those falling into the first quarter of the entire length of the 5’ UTRs of annotated genes, and then by excluding those that overlapped with the m6A sites identified above. Differential methylation analysis for m6A was performed using DEseq2 [79].

Metagene plots along the transcript and peak distribution across genomic regions (analyzing 5’ and 3’UTRs separately) were generated using the R package GenomicPlot (URL: https://github.com/shuye2009/GenomicPlot). To examine overlap of iCLIP and ChIP peaks, iCLIP CITS were extended to 50 nt on each side and a minimum of 1nt overlap between peaks from two datasets was set as a threshold. Raw GFP control and MKRN2 iCLIP-seq data were acquired from our previous report (GEO: GSE136399) [49], whereas iCLIP-seq data for PTBP1 and U2AF1 have been deposited as a part of our large-scale study submitted as a companion paper elsewhere (GEO: GSE230846).

### RNA-seq analysis

For differential expression analysis, reads were aligned to human genome GRCh38 using STAR (version 2.7.6.a) [80], gene-level read counts were quantified using RSEM (version 1.3.3) [81] and normalized by variance-stabilizing transformation using DESeq2 [79]. Adjusted p-value ≤ 0.05 and absolute log2fold change ± 1 were used to determine significantly regulated genes.

### ChIP-seq analysis

ChIP-seq analysis was performed essentially as described in our previous studies [45]. Briefly, Illumina adaptor sequences were removed from the 3′ ends of 51-nt reads, and the reads were mapped to the human genome hg19 using Bowtie 2 with default settings. Duplicate reads were discarded, and peaks were called jointly on the immunoprecipitated and input samples with MACS2 (version 2.1.2) [82]. ChIP-seq data have been deposited in GEO with the accession code GSE228608

### Data and code availability

- ChIP-seq, iCLIP-seq, miCLIP, and RNA-seq data generated in this paper can be found online at Gene Expression Omnibus (https://www.ncbi.nlm.nih.gov/geo/) with the accession code GSE228608. Data can be accessed with reviewers’ token ‘sxqjqecgxlsrjid’. (https://www.ncbi.nlm.nih.gov/geo/query/acc.cgi?acc=GSE228608)
- All original code has been deposited at Github (https://github.com/shuye2009/GenomicPlot).
- Any additional information required to reanalyze the data reported in this paper is available from the lead contact upon request.

## Acknowledgments

We thank Drs. Thomas Gonatopoulos-Pournatzis, Esha Sharma, Andrew Best, Zuyao Ni, Ernest Radovani, and Eric Wolf for helpful discussions, and Dave O’Hanlon for technical help. Dr. Eliza S. Lee is acknowledged for her help in IF analysis. Alexandra Nitoiu is gratefully acknowledged for her help in qPCR experiments. Sherin Shibin and team members at the Donnelly Sequencing Center are gratefully acknowledged for their assistance with next-generation sequencing.

## Author contributions

S.N.-S and J.F.G. conceived and designed the study. S.N.-S. performed all experiments (with support from others when stated below), participated in the computational analyses, and co- ordination of the project. S.P. performed computational analyses related to iCLIP- and RNA-seq with feedback from S.N.-S., U.B., and H.L. G.L.B. participated in co-IPs, knockdowns and qPCRs.

N.A. participated in iCLIP and qPCR experiments. S.F. performed IF experiments and edited the manuscript. M.U. participated in data analysis. G.Z., H.T. and E.M. took part in AP-MS and RNA-MS. Z.Z. supervised H.L. B.J.B. participated in data analysis and edited the manuscript. J.F.G. supervised S.N.-S., S.P., N.A., G.L.B., G.Z., H.T., and E.M., supervised the project, and participated in data analysis. S.N.-S. wrote the manuscript with feedback/edits from J.F.G. All authors have read and approved the final manuscript.

## References

1. Zhao BS, Roundtree IA, He C. Post-transcriptional gene regulation by mRNA modifications. Nat Rev Mol Cell Biol. 2017;18:31–42.

2. Roundtree IA, Evans ME, Pan T, He C. Dynamic RNA Modifications in Gene Expression Regulation. Cell. Cell Press; 2017. p. 1187–200.

3. Zhao BS, Wang X, Beadell A V., Lu Z, Shi H, Kuuspalu A, et al. m6A-dependent maternal mRNA clearance facilitates zebrafish maternal-to-zygotic transition. Nature 2017 542:7642. 2017;542:475–8.

4. Chen T, Hao YJ, Zhang Y, Li MM, Wang M, Han W, et al. m(6)A RNA methylation is regulated by microRNAs and promotes reprogramming to pluripotency. Cell Stem Cell. 2015;16:289–301.

5. Zhao X, Yang Y, Sun BF, Shi Y, Yang X, Xiao W, et al. FTO-dependent demethylation of N6-methyladenosine regulates mRNA splicing and is required for adipogenesis. Cell Res. 2014;24:1403.

6. Jia G, Fu Y, Zhao X, Dai Q, Zheng G, Yang Y, et al. N6-methyladenosine in nuclear RNA is a major substrate of the obesity-associated FTO. Nat Chem Biol. 2011;7:885–7.

7. Jiang X, Liu B, Nie Z, Duan L, Xiong Q, Jin Z, et al. The role of m6A modification in the biological functions and diseases. Signal Transduction and Targeted Therapy 2021 6:1. 2021;6:1–16.

8. Liu J, Yue Y, Han D, Wang X, Fu Y, Zhang L, et al. A METTL3-METTL14 complex mediates mammalian nuclear RNA N6-adenosine methylation. Nat Chem Biol. 2014;10:93–5.

9. Zheng G, Dahl JA, Niu Y, Fedorcsak P, Huang CM, Li CJ, et al. ALKBH5 is a mammalian RNA demethylase that impacts RNA metabolism and mouse fertility. Mol Cell. 2013;49:18–29.

10. Zaccara S, Ries RJ, Jaffrey SR. Reading, writing and erasing mRNA methylation. Nature Reviews Molecular Cell Biology 2019 20:10. 2019;20:608–24.

11. Chen L, Zhang C, Ma W, Huang J, Zhao Y, Liu H. METTL3-mediated m6A modification stabilizes TERRA and maintains telomere stability. Nucleic Acids Res. 2022;50:11619–34.

12. Chen XY, Zhang J, Zhu JS. The role of m6A RNA methylation in human cancer. Mol Cancer. BioMed Central Ltd.; 2019.

13. He L, Li H, Wu A, Peng Y, Shu G, Yin G. Functions of N6-methyladenosine and its role in cancer. Mol Cancer. BioMed Central Ltd.; 2019.

14. Dominissini D, Moshitch-Moshkovitz S, Schwartz S, Salmon-Divon M, Ungar L, Osenberg S, et al. Topology of the human and mouse m6A RNA methylomes revealed by m6A-seq. Nature. 2012;485:201–6.

15. Meyer KD, Saletore Y, Zumbo P, Elemento O, Mason CE, Jaffrey SR. Comprehensive analysis of mRNA methylation reveals enrichment in 3′ UTRs and near stop codons. Cell. 2012;149:1635–46.

16. Shi R, Ying S, Li Y, Zhu L, Wang X, Jin H. Linking the YTH domain to cancer: the importance of YTH family proteins in epigenetics. Cell Death & Disease 2021 12:4. 2021;12:1– 14.

17. Alarcón CR, Goodarzi H, Lee H, Liu X, Tavazoie S, Tavazoie SF. HNRNPA2B1 Is a Mediator of m(6)A-Dependent Nuclear RNA Processing Events. Cell. 2015;162:1299–308.

18. Zarnack K, König J, Tajnik M, Martincorena I, Eustermann S, Stévant I, et al. Direct competition between hnRNP C and U2AF65 protects the transcriptome from the exonization of Alu elements. Cell. 2013;152:453–66.

19. Huang H, Weng H, Sun W, Qin X, Shi H, Wu H, et al. Recognition of RNA N6- methyladenosine by IGF2BP Proteins Enhances mRNA Stability and Translation. Nat Cell Biol. 2018;20:285.

20. Liu N, Dai Q, Zheng G, He C, Parisien M, Pan T. N6-methyladenosine-dependent RNA structural switches regulate RNA-protein interactions. Nature. 2015;518:560.

21. Wang X, Lu Z, Gomez A, Hon GC, Yue Y, Han D, et al. N6-methyladenosine-dependent regulation of messenger RNA stability. Nature. 2014;505:117–20.

22. Keith JM, Ensinger MJ, Moss B. HeLa cell RNA (2’-O-methyladenosine-N6-)- methyltransferase specific for the capped 5’-end of messenger RNA. J Biol Chem. 1978;253:5033–41.

23. Wei CM, Gershowitz A, Moss B. Methylated nucleotides block 5’ terminus of HeLa cell messenger RNA. Cell. 1975;4:379–86.

24. Ben-Haim MS, Pinto Y, Moshitch-Moshkovitz S, Hershkovitz V, Kol N, Diamant-Levi T, et al. Dynamic regulation of N6,2’-O-dimethyladenosine (m6Am) in obesity. Nat Commun. 2021;12.

25. Mauer J, Luo X, Blanjoie A, Jiao X, Grozhik A v., Patil DP, et al. Reversible methylation of m6Am in the 5′ cap controls mRNA stability. Nature. 2017;541:371–5.

26. Sendinc E, Valle-Garcia D, Dhall A, Chen H, Henriques T, Navarrete-Perea J, et al. PCIF1 Catalyzes m6Am mRNA Methylation to Regulate Gene Expression. Mol Cell. 2019;75:620–630.e9.

27. Relier S, Ripoll J, Guillorit H, Amalric A, Achour C, Boissière F, et al. FTO-mediated cytoplasmic m6Am demethylation adjusts stem-like properties in colorectal cancer cell. Nat Commun. 2021;12.

28. Linder B, Grozhik A V., Olarerin-George AO, Meydan C, Mason CE, Jaffrey SR. Single-nucleotide-resolution mapping of m6A and m6Am throughout the transcriptome. Nat Methods. 2015;12:767–72.

29. Akichika S, Hirano S, Shichino Y, Suzuki T, Nishimasu H, Ishitani R, et al. Cap-specific terminal N 6-methylation of RNA by an RNA polymerase II-associated methyltransferase. Science. 2019;363.

30. Boulias K, Toczydłowska-Socha D, Hawley BR, Liberman N, Takashima K, Zaccara S, et al. Identification of the m6Am Methyltransferase PCIF1 Reveals the Location and Functions of m6Am in the Transcriptome. Mol Cell. 2019;75:631–643.e8.

31. Sun H, Zhang M, Li K, Bai D, Yi C. Cap-specific, terminal N6-methylation by a mammalian m6Am methyltransferase. Cell Res. 2019;29:80–2.

32. Wei J, Liu F, Lu Z, Fei Q, Ai Y, He PC, et al. Differential m6A, m6Am, and m1A Demethylation Mediated by FTO in the Cell Nucleus and Cytoplasm. Mol Cell. 2018;71:973- 985.e5.

33. Li Y, Wu K, Quan W, Yu L, Chen S, Cheng C, et al. The dynamics of FTO binding and demethylation from the m6A motifs. RNA Biol. 2019;16:1179.

34. Relier S, Rivals E, David A. The multifaceted functions of the Fat mass and Obesity-associated protein (FTO) in normal and cancer cells. RNA Biol. 2022;19:132–42.

35. Zhang X, Wei LH, Wang Y, Xiao Y, Liu J, Zhang W, et al. Structural insights into FTO’s catalytic mechanism for the demethylation of multiple RNA substrates. Proc Natl Acad Sci U S A. 2019;116:2919–24.

36. Zou S, Toh JDW, Wong KHQ, Gao YG, Hong W, Woon ECY. N(6)-Methyladenosine: a conformational marker that regulates the substrate specificity of human demethylases FTO and ALKBH5. Sci Rep. 2016;6.

37. Bartosovic M, Molares HC, Gregorova P, Hrossova D, Kudla G, Vanacova S. N6- methyladenosine demethylase FTO targets pre-mRNAs and regulates alternative splicing and 3′- end processing. Nucleic Acids Res. 2017;45:11356–70.

38. Edupuganti RR, Geiger S, Lindeboom RGH, Shi H, Hsu PJ, Lu Z, et al. N6-methyladenosine (m6A) recruits and repels proteins to regulate mRNA homeostasis. Nat Struct Mol Biol. 2017;24:870–8.

39. Yoon JH, Choi W il, Jeon BN, Koh DI, Kim MK, Kim MH, et al. Human Kruppel-related 3 (HKR3) is a novel transcription activator of alternate reading frame (ARF) gene. J Biol Chem. 2014;289:4018–31.

40. Jahn A, Rane G, Paszkowski-Rogacz M, Sayols S, Bluhm A, Han C, et al. ZBTB48 is both a vertebrate telomere-binding protein and a transcriptional activator. EMBO Rep. 2017;18:929–46.

41. Li JSZ, Fusté JM, Simavorian T, Bartocci C, Tsai J, Karlseder J, et al. TZAP: A telomere-associated protein involved in telomere length control. Science. 2017;355:638–41.

42. Maris JM, Jensen J, Sulman EP, Beltinger CP, Allen C, Biegel JA, et al. Human Krüppel- related 3 (HKR3): a candidate for the 1p36 neuroblastoma tumour suppressor gene? Eur J Cancer. 1997;33:1991–6.

43. Maris JM, Jensen J, Sulman EP, Beltinger CP, Allen C, Biegel JA, et al. Human Krüppel- related 3 (HKR3): a candidate for the 1p36 neuroblastoma tumour suppressor gene? Eur J Cancer. 1997;33:1991–6.

44. Bauer A, Savelyeva L, Claas A, Praml C, Berthold F, Schwab M. Smallest region of overlapping deletion in 1p36 in human neuroblastoma: a 1 Mbp cosmid and PAC contig. Genes Chromosomes Cancer. 2001;31:228–39.

45. Schmitges FW, Radovani E, Najafabadi HS, Barazandeh M, Campitelli LF, Yin Y, et al. Multiparameter functional diversity of human C2H2 zinc finger proteins. Genome Res. 2016;26:1742–52.

46. Teo G, Liu G, Zhang J, Nesvizhskii AI, Gingras A-C, Choi H. SAINTexpress: improvements and additional features in Significance Analysis of INTeractome software. J Proteomics. 2014;100:37–43.

47. Mellacheruvu D, Wright Z, Couzens AL, Lambert J-P, St-Denis NA, Li T, et al. The CRAPome: a contaminant repository for affinity purification-mass spectrometry data. Nat Methods [Internet]. 2013 [cited 2019 Feb 22];10:730–6. Available from: http://www.ncbi.nlm.nih.gov/pubmed/23921808

48. Song J, Nabeel-Shah S, Pu S, Lee H, Braunschweig U, Ni Z, et al. Regulation of alternative polyadenylation by the C2H2-zinc-finger protein Sp1. Mol Cell. 2022;82:3135–3150.e9.

49. Wolf EJ, Miles A, Lee ES, Nabeel-Shah S, Greenblatt JF, Palazzo AF, et al. MKRN2 Physically Interacts with GLE1 to Regulate mRNA Export and Zebrafish Retinal Development. Cell Rep. 2020;31.

50. Sigova AA, Abraham BJ, Ji X, Molinie B, Hannett NM, Guo YE, et al. Transcription factor trapping by RNA in gene regulatory elements. Science. 2015;350:978–81.

51. Vaid R, Thombare K, Mendez A, Burgos-Panadero R, Djos A, Jachimowicz D, et al. m6A modification of TERRA RNA is required for telomere maintenance and is a therapeutic target for ALT positive Neuroblastoma. bioRxiv [Internet]. 2022 [cited 2023 Feb 22];2022.12.09.519591. Available from: https://www.biorxiv.org/content/10.1101/2022.12.09.519591v1

52. Abid HZ, McCaffrey J, Raseley K, Young E, Lassahn K, Varapula D, et al. Single-molecule analysis of subtelomeres and telomeres in Alternative Lengthening of Telomeres (ALT) cells. BMC Genomics. 2020;21:1–17.

53. Ruan DY, Li T, Wang YN, Meng Q, Li Y, Yu K, et al. FTO downregulation mediated by hypoxia facilitates colorectal cancer metastasis. Oncogene. 2021;40:5168–81.

54. Jung SJ, Seo YR, Park WJ, Heo YR, Lee YH, Kim S, et al. Clinicopathological Characteristics of TZAP Expression in Colorectal Cancers. Onco Targets Ther. 2020;13:12933.

55. Sen N, Gui B, Kumar R. Role of MTA1 in cancer progression and metastasis. Cancer Metastasis Rev. 2014;33:879–89.

56. Ontiveros, Jordan R, Shen H, Stoute J, Yanas A, Cui Y, Zhang Y, et al. Coordination of mRNA and tRNA methylations by TRMT10A. Proc Natl Acad Sci U S A. 2020;117:7782–91.

57. Song H, Wang Y, Wang R, Zhang X, Liu Y, Jia G, et al. SFPQ Is an FTO-Binding Protein that Facilitates the Demethylation Substrate Preference. Cell Chem Biol. 2020;27:283–291.e6.

58. Covelo-Molares H, Obrdlik A, Poštulková I, Dohnálková M, Gregorová P, Ganji R, et al. The comprehensive interactomes of human adenosine RNA methyltransferases and demethylases reveal distinct functional and regulatory features. Nucleic Acids Res. 2021;49:10895–910.

59. Patil DP, Chen CK, Pickering BF, Chow A, Jackson C, Guttman M, et al. m(6)A RNA methylation promotes XIST-mediated transcriptional repression. Nature. 2016;537:369–73.

60. Zheng Q, Hou J, Zhou Y, Li Z, Cao X. The RNA helicase DDX46 inhibits innate immunity by entrapping m6A-demethylated antiviral transcripts in the nucleus. Nature Immunology 2017 18:10. 2017;18:1094–103.

61. Yu F, Zhu AC, Liu S, Gao B, Wang Y, Khudaverdyan N, et al. RBM33 is a unique m6A RNA-binding protein that regulates ALKBH5 demethylase activity and substrate selectivity. Mol Cell. 2023;83:2003–2019.e6.

62. Liu SJ, Tang HL, He Q, Lu P, Fu T, Xu XL, et al. FTO is a transcriptional repressor to auto- regulate its own gene and potentially associated with homeostasis of body weight. J Mol Cell Biol [Internet]. 2019 [cited 2023 Jan 11];11:118. Available from: https://www.ncbi.nlm.nih.gov/pmc/articles/PMC6734146/

63. Li Y, Su R, Deng X, Chen Y, Chen J. FTO in cancer: functions, molecular mechanisms, and therapeutic implications. Trends Cancer. 2022;8:598–614.

64. Lee JH, Hong J, Zhang Z, de la Peña Avalos B, Proietti CJ, Deamicis AR, et al. Regulation of telomere homeostasis and genomic stability in cancer by N6-adenosine methylation (m6A). Sci Adv. 2021;7:7073–101.

65. Wei J, Yu X, Yang L, Liu X, Gao B, Huang B, et al. FTO mediates LINE1 m6A demethylation and chromatin regulation in mESCs and mouse development. Science. 2022;376.

66. Nabeel-Shah S, Lee H, Ahmed N, Burke GL, Farhangmehr S, Ashraf K, et al. SARS-CoV-2 nucleocapsid protein binds host mRNAs and attenuates stress granules to impair host stress response. iScience. 2022;25.

67. Huppertz I, Attig J, D’Ambrogio A, Easton LE, Sibley CR, Sugimoto Y, et al. iCLIP: Protein–RNA interactions at nucleotide resolution. Methods. 2014;65:274–87.

68. Nabeel-Shah S, Greenblatt J. Revised iCLIP-seq Protocol for Profiling RNA-protein Interaction Sites at Individual Nucleotide Resolution in Living Cells. Bio Protoc. 2023;13.

69. Nostrand EL Van, Pratt GA, Shishkin AA, Gelboin-burkhart C, Fang MY, Sundararaman B, et al. Robust transcriptome-wide discovery of RNA-binding protein binding sites with enhanced CLIP (eCLIP). Nat Methods. 2016;13:508–514.

70. Livak KJ, Schmittgen TD. Analysis of relative gene expression data using real-time quantitative PCR and the 2-ΔΔCT method. Methods. 2001;25:402–8.

71. Martindale JL, Gorospe M, Idda ML. Ribonucleoprotein Immunoprecipitation (RIP) Analysis. Bio Protoc. 2020;10.

72. Shen L, Liang Z, Yu H. Dot Blot Analysis of N6-methyladenosine RNA Modification Levels. Bio Protoc. 2017;7.

73. Cormier N, Yeo A, Fiorentino E, Paxson J. Optimization of the Wound Scratch Assay to Detect Changes in Murine Mesenchymal Stromal Cell Migration After Damage by Soluble Cigarette Smoke Extract. J Vis Exp. 2015;2015:53414.

74. Suarez-Arnedo A, Figueroa FT, Clavijo C, Arbeláez P, Cruz JC, Muñoz-Camargo C. An image J plugin for the high throughput image analysis of in vitro scratch wound healing assays. PLoS One. 2020;15.

75. Han H, Braunschweig U, Gonatopoulos-pournatzis T, Wrana JL, Moffat J, Blencowe BJ, et al. Multilayered Control of Alternative Splicing Regulatory Networks by Transcription Factors Resource Multilayered Control of Alternative Splicing Regulatory Networks by Transcription Factors. Mol Cell. 2017;65:539–553.e7.

76. Bolger AM, Lohse M, Usadel B. Trimmomatic: A flexible trimmer for Illumina sequence data. Bioinformatics. 2014;30:2114–20.

77. Trapnell C, Pachter L, Salzberg SL. TopHat: Discovering splice junctions with RNA-Seq. Bioinformatics. 2009;25:1105–11.

78. Shah A, Qian Y, Weyn-Vanhentenryck SM, Zhang C. CLIP Tool Kit (CTK): A flexible and robust pipeline to analyze CLIP sequencing data. Bioinformatics. 2017;33:566–7.

79. Love MI, Huber W, Anders S. Moderated estimation of fold change and dispersion for RNA-seq data with DESeq2. Genome Biol. 2014;15:550.

80. Dobin A, Davis CA, Schlesinger F, Drenkow J, Zaleski C, Jha S, et al. STAR: Ultrafast universal RNA-seq aligner. Bioinformatics. 2013;29:15–21.

81. Li B, Dewey CN. RSEM: accurate transcript quantification from RNA-Seq data with or without a reference genome. BMC Bioinformatics. 2011;12.

82. Feng J, Liu T, Qin B, Zhang Y, Liu XS. Identifying ChIP-seq enrichment using MACS. Nat Protoc. 2012;7:1728–40.

