## Supplemental Figures for "Recruitment of the m^6^A/Am demethylase FTO to target RNAs by the telomeric zinc finger protein ZBTB48"

### Figure S1

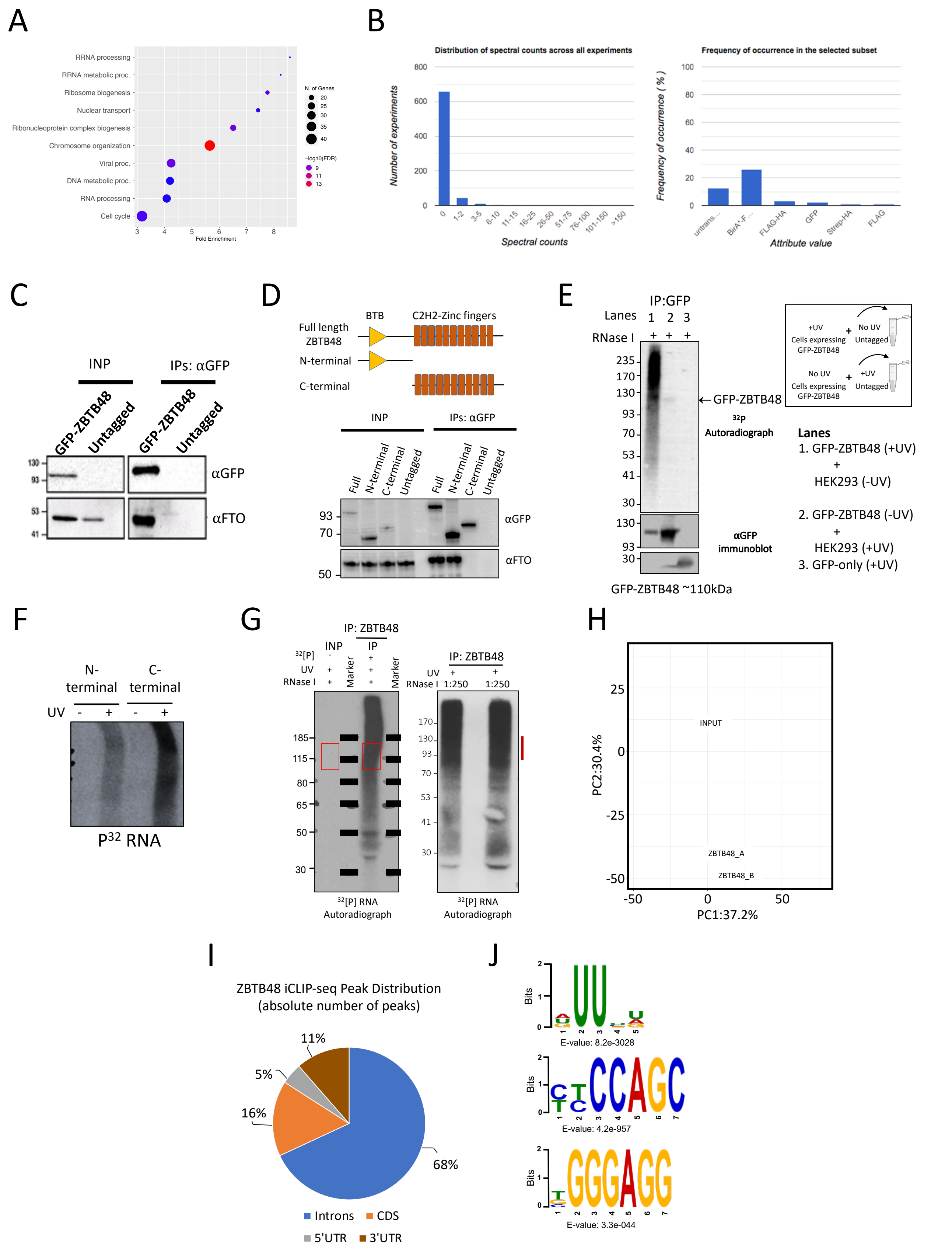

A
RRNA processing
RRNA metabolic proc.
Ribosome biogenesis
Nuclear transport
Ribonucleoprotein complex biogenesis
Chromosome organization
Viral proc.
DNA metabolic proc.
RNA processing
Cell cycle
Fold Enrichment
N. of Genes
−log10(FDR)
B
Distribution of spectral counts across all experiments
Number of experiments
Spectral counts
Frequency of occurrence in the selected subset
Frequency of occurrence ( % )
Attribute value
C
INP
IPs: αGFP
GFP-ZBTB48
Untagged
αGFP
αFTO
D
BTB
C2H2-Zinc fingers
Full length ZBTB48
N-terminal
C-terminal
Full
Untagged
E
IP:GFP
Lanes 1 2 3
RNase I + + +
← GFP-ZBTB48
32P Autoradiograph
αGFP immunoblot
GFP-ZBTB48 ~110kDa
+UV Cells expressing GFP-ZBTB48
No UV Untagged
No UV Cells expressing GFP-ZBTB48
+UV Untagged
Lanes
1. GFP-ZBTB48 (+UV) + HEK293 (-UV)
2. GFP-ZBTB48 (-UV) + HEK293 (+UV)
3. GFP-only (+UV)
F
N-terminal
C-terminal
UV - + - +
P32 RNA
G
IP: ZBTB48
INP
IP
32[P]
UV
RNase I
Marker
1:250
32[P] RNA Autoradiograph
H
INPUT
ZBTB48_A
ZBTB48_B
PC1:37.2%
PC2:30.4%
I
ZBTB48 iCLIP-seq Peak Distribution (absolute number of peaks)
68%
16%
5%
11%
Introns
CDS
5'UTR
3'UTR
J
Bits
E-value: 8.2e-3028
E-value: 4.2e-957
E-value: 3.3e-044

**Figure S1: ZBTB48 interacts with FTO and binds to RNA in cells. A:** GO enrichment analysis related to biological processes using ZBTB48 interaction partners as reported previously in Schmitges et al. 2016. Plot legend indicating the p-values and gene count for each GO term is provided. **B:** Bar plot showing the distribution of FTO spectral counts across AP-MS experiments (left) and frequency of FTO occurrence for various epitope tags (right). The analysis was performed using the CRAPome database (https://reprint-apms.org/?q=chooseworkflow). **C:** Interaction of ZBTB48 with FTO detected by co-immunoprecipitation. IPs were performed with GFP antibody (10 ug) using whole-cell lysates prepared from HEK293 cells expressing or not expressing GFP-tagged ZBTB48. Cell lysates were treated with Benzonase (nuclease) prior to IPs. Blots were probed with the indicated antibodies. **D:** Top: Schematic representation of GFP-tagged ZBTB48 full-length, N-terminal, and C-terminal regions. Bottom: Interaction of GFP-ZBTB48 truncation mutants expressed in HEK293 cells with FTO detected by co-immunoprecipitation. IPs were performed with GFP antibody. Blots were probed with the indicated antibodies. **E:** CLIP-autoradiography of ZBTB48 using mixed cells. Equal numbers of +UV GFP-ZBTB48 and no-UV wildtype HEK293, as well as no-UV GFP-ZBTB48 cells and +UV wildtype HEK293 cells, were mixed prior to CLIP-autoradiography. GFP-only cells were used as an additional control. Bottom panel indicates the recovery of GFP-tagged proteins. Schematic of the mixed cells is shown on the right side. **F:** Autoradiographs of $^{32}$P-labeled immunopurified complexes containing ZBTB48 truncation mutants and their bound RNAs. Cells expressing GFP-tagged N-terminal or C-terminal ZBTB48 truncation mutants were UV crosslinked and IPs were performed using anti-GFP antibody after partial RNase I digestion. Samples without UV crosslinking were used as controls. **G:** Autoradiographs of input (INP) and $^{32}$P-labeled immunopurified (IP) ZBTB48-RNA complexes after partial RNase I digestion (RNase I dilution: 1:250). Complexes were resolved on 4-12% Bis-Tris gels and transferred to nitrocellulose membranes. Size matched input (2% of the starting material) was also resolved along with the IP. Red box and vertical red line on the right indicate the excised areas for iCLIP-seq library preparations. **H:** Principal component analysis (PCA) of ZBTB48 iCLIP-seq replicates. **I:** Pie chart showing the distribution of ZBTB48 iCLIP-seq peaks across protein-codong genes. Absolute number of peaks is used. **J:** RNA motifs identified in ZBTB48 iCLIP-seq. The iCLIP-seq CITS were used for *de novo* motif discovery. p-value was calculated against randomly assorted sequences.

### Figure S2

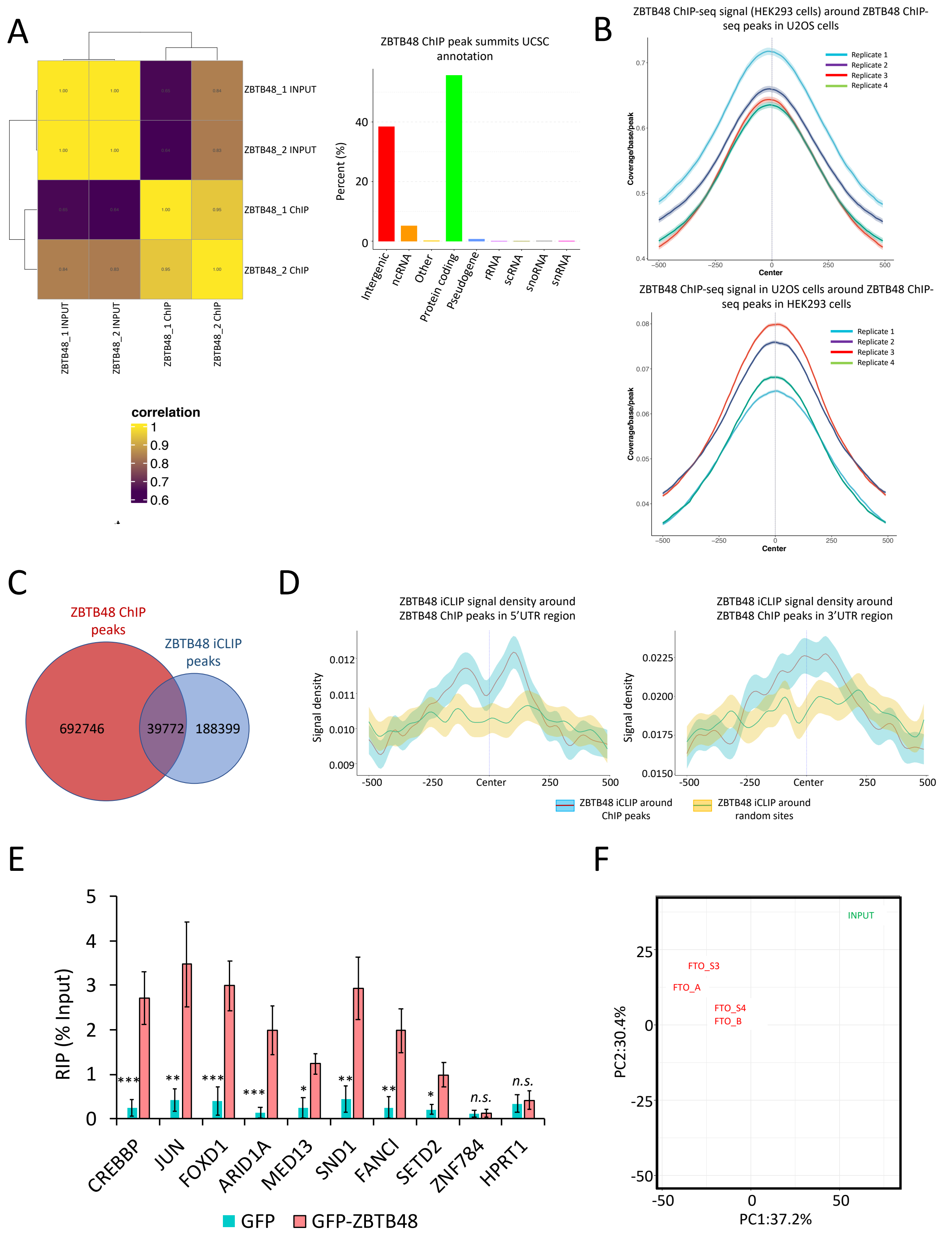
A
ZBTB48_1 INPUT
ZBTB48_2 INPUT
ZBTB48_1 ChIP
ZBTB48_2 ChIP
correlation
1
0.9
0.8
0.7
0.6
ZBTB48 ChIP peak summits UCSC annotation
Percent (%)
Intergenic
ncRNA
Other
Protein coding
Pseudogene
rRNA
scRNA
snoRNA
snRNA
B
ZBTB48 ChIP-seq signal (HEK293 cells) around ZBTB48 ChIP-seq peaks in U2OS cells
Replicate 1
Replicate 2
Replicate 3
Replicate 4
Coverage/base/peak
Center
ZBTB48 ChIP-seq signal in U2OS cells around ZBTB48 ChIP-seq peaks in HEK293 cells
C
ZBTB48 ChIP peaks
ZBTB48 iCLIP peaks
692746
39772
188399
D
ZBTB48 iCLIP signal density around ZBTB48 ChIP peaks in 5'UTR region
ZBTB48 iCLIP signal density around ZBTB48 ChIP peaks in 3'UTR region
Signal density
ZBTB48 iCLIP around ChIP peaks
ZBTB48 iCLIP around random sites
E
RIP (% Input)
CREBBP
JUN
FOXD1
ARID1A
MED13
SND1
FANCI
SETD2
ZNF784
HPRT1
n.s.
GFP
GFP-ZBTB48
F
INPUT
FTO_S3
FTO_A
FTO_S4
FTO_B
PC2:30.4%
PC1:37.2%

**Figure S2: ZBTB48 has distinct RNA- and DNA-binding patterns. A:** Left, Heatmap showing the correlation between ZBTB48 ChIP-seq replicates. Pearson co-efficient of correlation values are indicated. Right, Bar plot showing the distribution of the localization of ZBTB48 ChIP-seq peaks. ChIP-seq was performed in cells overexpressing GFP-tagged ZBTB48. **B:** GFP-ZBTB48 ChIP-seq signal density in HEK293 cells around endogenous ZBTB48 ChIP peaks identified in U2OS cells (top) and vice versa (bottom). Different colors represent 4 biological replicates in U2OS cells (Jahn et al., 2017; PMID: 28500257). **C:** Venn diagram showing the overlap of ZBTB48 ChIP and iCLIP peaks. iCLIP-seq CITS and ChIPseq peaks were extended by 100nt (50nt on each side of the identified peak center) and an overlap of at least 1nt was used as a threshold to be considered as an overlapping peak. **D:** Metagene profiles showing the iCLIP-seq signal density (CITS) of ZBTB48 around either ZBTB48 ChIP-seq peaks or random sites. Plots for the 5'UTR and 3'UTR are shown separately. Shaded area indicates the standard error of mean (SEM) for iCLIP-seq replicates. **E:** Bar graph representation of GFP-ZBTB48 RIP-qPCR showing binding of ZBTB48 to the indicated transcripts (biological replicates n = 4, student's t test, $***p \leq 0.001$, $**p \leq 0.01$, $*p \leq 0.05$, error bars: standard error of mean (SEM)). **F**: Principal component analysis (PCA) of FTO iCLIP-seq replicates.

### Figure S3

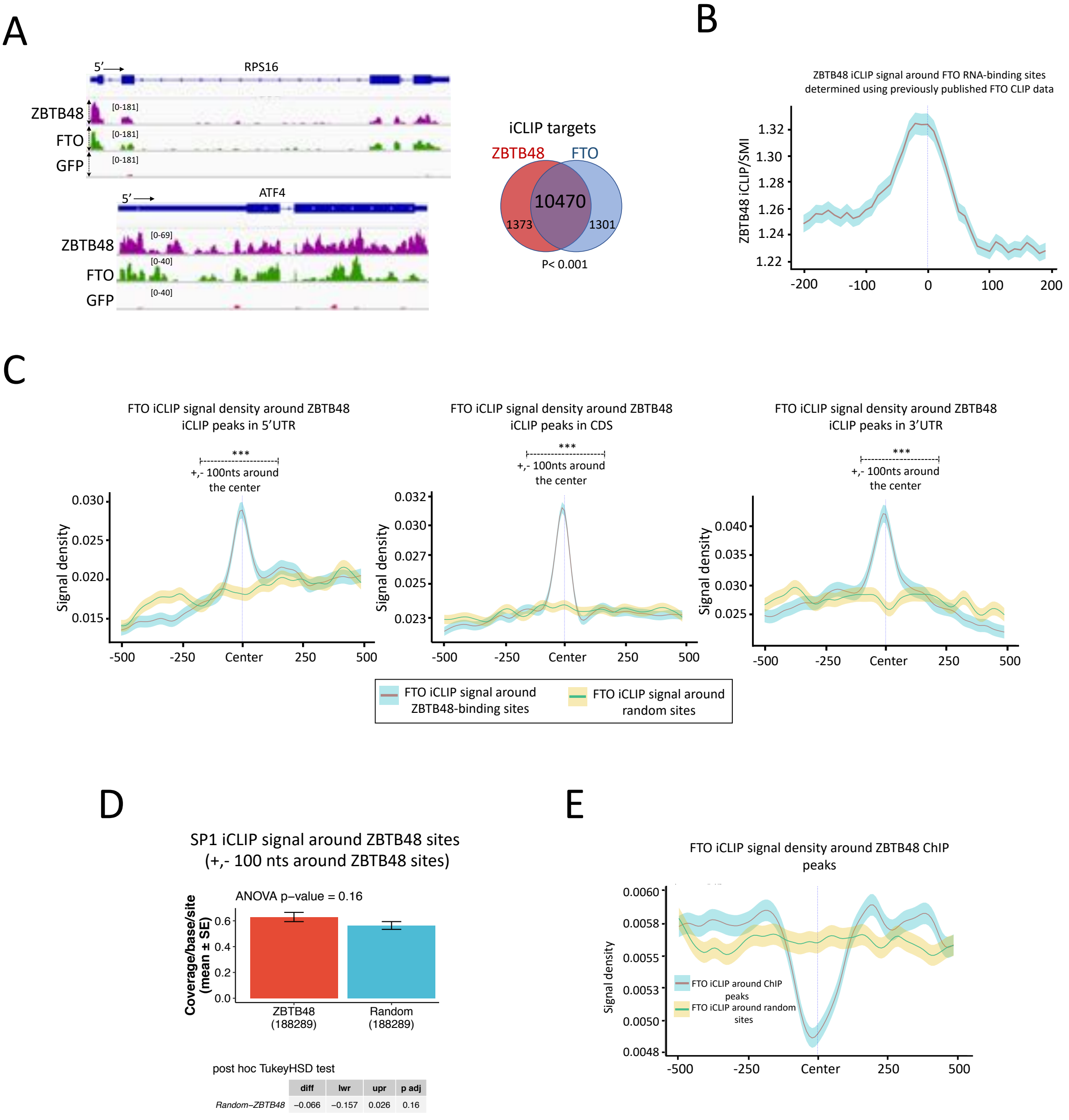

A
5′
RPS16
ZBTB48
FTO
GFP
[0-181]
[0-181]
[0-181]
5′
ATF4
ZBTB48
FTO
GFP
[0-69]
[0-40]
[0-40]
iCLIP targets
ZBTB48
FTO
10470
1373
1301
P< 0.001
B
ZBTB48 iCLIP signal around FTO RNA-binding sites determined using previously published FTO CLIP data
ZBTB48 iCLIP/SMI
1.32
1.30
1.28
1.26
1.24
1.22
-200
-100
0
100
200
C
FTO iCLIP signal density around ZBTB48 iCLIP peaks in 5′UTR
***
+,- 100nts around the center
Signal density
0.030
0.025
0.020
0.015
-500
-250
Center
250
500
FTO iCLIP signal density around ZBTB48 iCLIP peaks in CDS
***
+,- 100nts around the center
Signal density
0.032
0.030
0.027
0.025
0.023
-500
-250
Center
250
500
FTO iCLIP signal density around ZBTB48 iCLIP peaks in 3′UTR
***
+,- 100nts around the center
Signal density
0.040
0.035
0.030
0.025
-500
-250
Center
250
500
FTO iCLIP signal around ZBTB48-binding sites
FTO iCLIP signal around random sites
D
SP1 iCLIP signal around ZBTB48 sites (+,- 100 nts around ZBTB48 sites)
ANOVA p−value = 0.16
Coverage/base/site (mean ± SE)
0.6
0.4
0.2
0.0
ZBTB48 (188289)
Random (188289)
post hoc TukeyHSD test
diff
lwr
upr
p adj
Random−ZBTB48
−0.066
−0.157
0.026
0.16
E
FTO iCLIP signal density around ZBTB48 ChIP peaks
Signal density
0.0060
0.0058
0.0055
0.0053
0.0050
0.0048
FTO iCLIP around ChIP peaks
FTO iCLIP around random sites
-500
-250
Center
250
500

**Figure S3: FTO and ZBTB48 RNA-binding sites coincide. A:** Left, Genome browser snapshots of the indicated genes showing the locations and coverage of ZBTB48, FLAG-FTO, and GFP-only iCLIP-seq signals. The 5' end of each gene is indicated with an arrow. Right, Venn diagram showing the overlap of ZBTB48 and FLAG-FTO iCLIP-seq identified target RNAs (hypergeometric test). **B:** Metagene plot showing the size-matched input (SMI) normalized iCLIP-seq signal (iCLIP/SMI) for ZBTB48 around FTO binding sites (indicated as '0') acquired from previously published FTO CLIP-seq data in HEK293 cells (Bartosovic et al. 2017; PMID: 28977517). **C:** Metagene plots indicate the FLAG-FTO iCLIP-seq peaks density around ZBTB48-binding sites in the 5'UTR (left), CDS (middle), and 3'UTR (right) (***$p \leq 0.001$, Wilcoxon (Mann-Whitney) test). **D:** Bar plot shows the mean signal intensity of SP1 iCLIP-seq peaks around either ZBTB48 iCLIP-seq peaks or random sites. **E:** Metagene plot shows the binding density of FLAG-FTO (iCLIP-seq peaks) around ZBTB48 ChIP-seq peaks.

### Figure S4

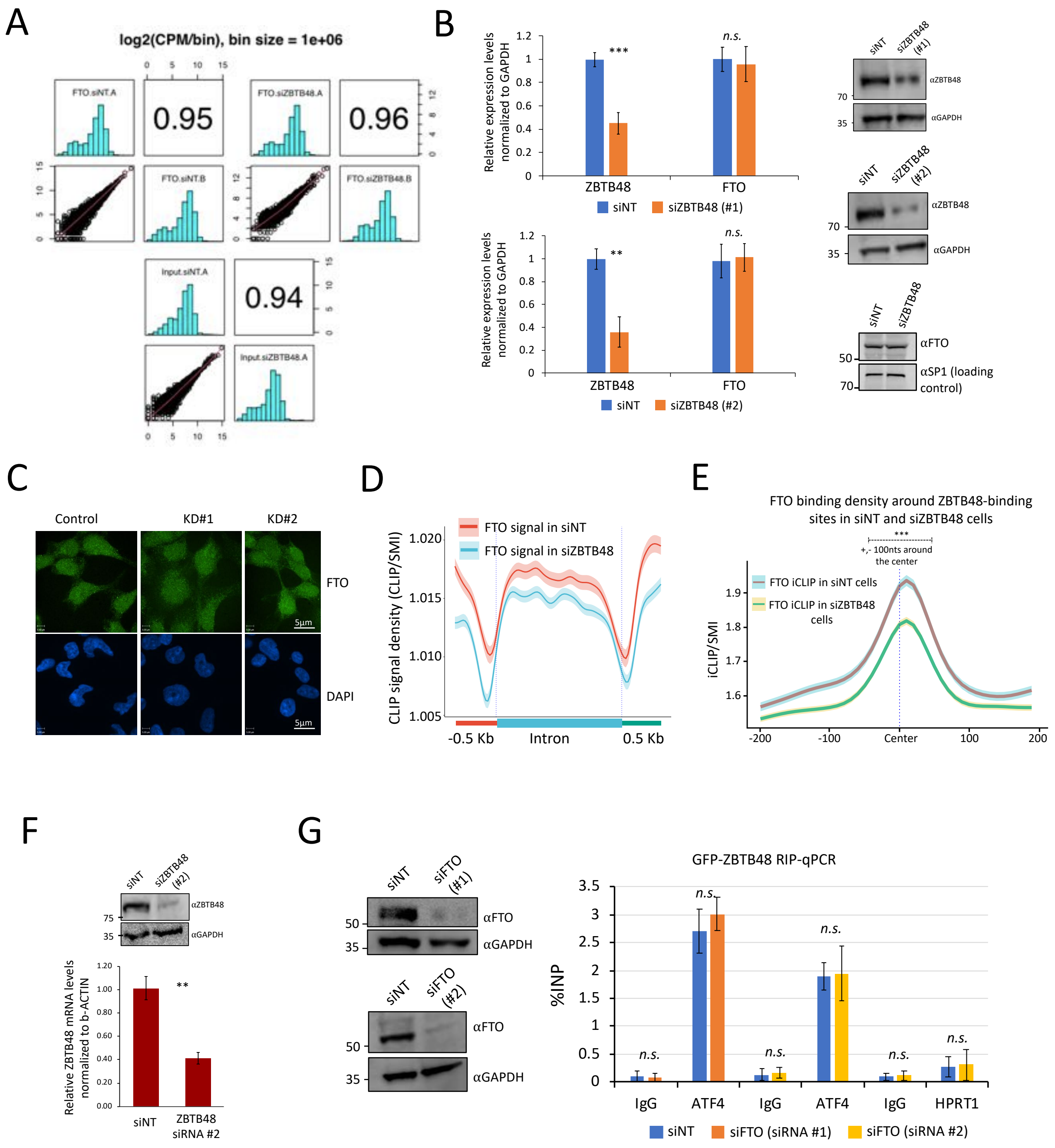
A
log2(CPM/bin), bin size = 1e+06
FTO.siNT.A
0.95
FTO.siZBTB48.A
0.96
FTO.siNT.B
FTO.siZBTB48.B
Input.siNT.A
0.94
Input.siZBTB48.A
B
Relative expression levels normalized to GAPDH
ZBTB48
FTO
***
n.s.
siNT
siZBTB48 (#1)
**
siZBTB48 (#2)
siNT
siZBTB48 (#1)
αZBTB48
αGAPDH
70
35
siZBTB48 (#2)
siZBTB48
αFTO
αSP1 (loading control)
50
C
Control
KD#1
KD#2
FTO
DAPI
5μm
D
FTO signal in siNT
FTO signal in siZBTB48
CLIP signal density (CLIP/SMI)
-0.5 Kb
Intron
0.5 Kb
E
FTO binding density around ZBTB48-binding sites in siNT and siZBTB48 cells
***
+,- 100nts around the center
FTO iCLIP in siNT cells
FTO iCLIP in siZBTB48 cells
iCLIP/SMI
-200
-100
Center
100
200
F
αZBTB48
αGAPDH
75
Relative ZBTB48 mRNA levels normalized to b-ACTIN
**
siNT
ZBTB48 siRNA #2
G
siNT
siFTO (#1)
siFTO (#2)
αFTO
αGAPDH
GFP-ZBTB48 RIP-qPCR
%INP
n.s.
IgG
ATF4
HPRT1
siNT
siFTO (siRNA #1)
siFTO (siRNA #2)

**Figure S4: ZBTB48 modulates FTO's RNA binding on target transcripts. A:** Correlation for FTO iCLIP-seq replicates in siNT and siZBTB48 cells. Pearson co-efficient of correlation values are indicated. CPM: crosslinks per million **B:** Left, Bar graphs represent qRT-PCR results to examine the expression of ZBTB48 and FTO in ZBTB48 knockdown cells (siZBTB48 # 1 and siZBTB48 # 2). The experiments were performed in biological triplicates, and p-values were calculated using the student's t-test ($***p \leq 0.001$, $**p \leq 0.01$, $*p \leq 0.05$, n.s.: non-significant). Right, Western blots showing the ZBTB48 protein levels in ZBTB48 knockdown cells (siZBTB48 # 1 and siZBTB48 # 2). FTO protein levels in whole cell extracts prepared from siNT or siZBTB48-treated cells are also shown. Blots were probed with the indicated antibodies. **C:** Representative immunofluorescence (IF) images showing the localization of FTO in control (siNT) or ZBTB48 KD HEK293 cells. ZBTB48 KDs were performed using two different siRNAs. Experiment was performed using an anti-FTO antibody. For nuclear counterstaining, DAPI was used. **D:** Standardized metaplot profiles showing the binding density (iCLIP/size-matched input) in introns and their surrounding exons of FTO in either siZBTB48 or siNT HEK293 cells. **E:** Metagene plots indicate the effect of ZBTB48 knockdown on FTO RNA-binding density around ZBTB48 RNA-binding sites ($***p \leq 0.001$, Wilcoxon (Mann-Whitney) test). **F:** Bottom, qRT-PCR results to examine the expression of ZBTB48 in HEK293 cells treated with siNT or siZBTB48 # 2. The experiments were performed in biological triplicates, and p-values were calculated using the student's t-test ($**p \leq 0.01$). Top panel, Western blotting analysis in whole cell lysates prepared from cells treated with siZBTB48 # 2. Blots were probed with the indicated antibodies. **G:** Left, Western blotting analysis in whole cell lysates prepared from cells treated either with siFTO #1 or siFTO #2. Blots were probed with the indicated antibodies. Right, Bar graph representation of GFP-ZBTB48 RIP-qPCR showing binding of ZBTB48 to ATF4 transcripts in cells treated with either siFTO #1 or siFTO #2 (biological replicates $n = 3$, student's t test, $***p \leq 0.001$, $**p \leq 0.01$, error bars: SEM).

### Figure S5

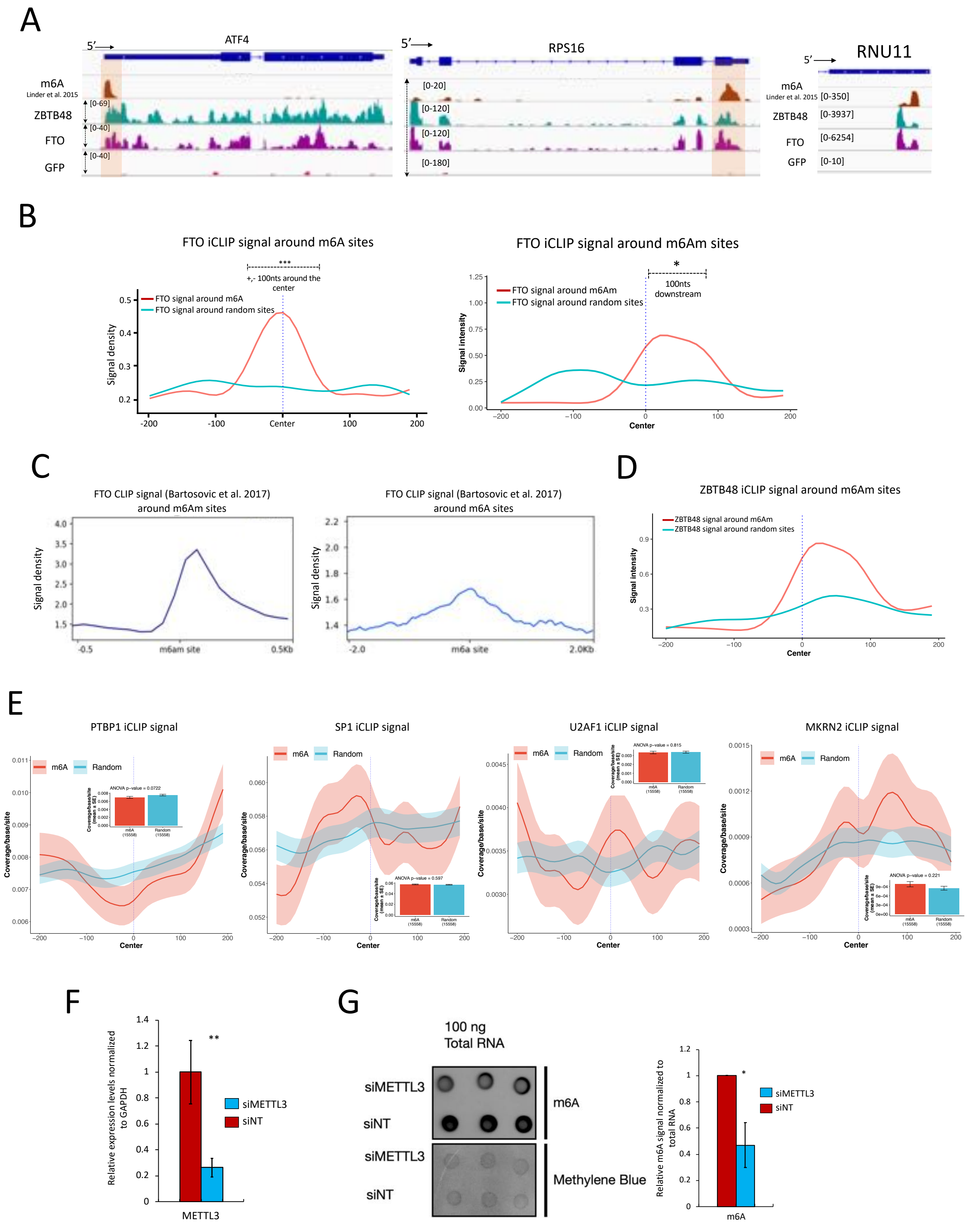

A
ATF4
RPS16
RNU11
m6A
Linder et al. 2015
ZBTB48
FTO
GFP
B
FTO iCLIP signal around m6A sites
FTO signal around m6A
FTO signal around random sites
+,- 100nts around the center
Signal density
FTO iCLIP signal around m6Am sites
FTO signal around m6Am
FTO signal around random sites
100nts downstream
Signal intensity
Center
C
FTO CLIP signal (Bartosovic et al. 2017) around m6Am sites
FTO CLIP signal (Bartosovic et al. 2017) around m6A sites
Signal density
m6am site
m6a site
D
ZBTB48 iCLIP signal around m6Am sites
ZBTB48 signal around m6Am
ZBTB48 signal around random sites
Signal intensity
E
PTBP1 iCLIP signal
SP1 iCLIP signal
U2AF1 iCLIP signal
MKRN2 iCLIP signal
m6A
Random
Coverage/base/site
ANOVA p-value = 0.0722
ANOVA p-value = 0.597
ANOVA p-value = 0.815
ANOVA p-value = 0.221
F
Relative expression levels normalized to GAPDH
siMETTL3
siNT
METTL3
G
100 ng Total RNA
siMETTL3
siNT
m6A
Methylene Blue
Relative m6A signal normalized to total RNA

**Figure S5: ZBTB48 and FTO bind around m6A/m6Am sites. A:** Genome browser shots showing the miCLIP-seq signal (Linder et al. 2015; PMID: 26121403), as well as the iCLIP-seq signals for ZBTB48, FTO, and GFP-only for the indicated genes. **B:** FTO iCLIP-seq peak densities around m6A (left) and m6Am (right) sites in comparison with random sites. m6A/m6Am sites were identified using previously published miCLIP-seq data (Linder et al. 2015; PMID: 26121403) ($***p \leq 0.001$, $*p \leq 0.01$, Wilcoxon (Mann-Whitney) test). **C:** Metagene plots showing FTO CLIP-seq signal densities around m6A and m6Am sites. Previously published FTO CLIP-seq data was used (Bartosovic et al. 2017; PMID: 28977517). **D:** ZBTB48 iCLIP-seq peak density around m6Am or random sites. **E:** iCLIP-seq peak densities around m6A or random sites for PTBP1, SP1, U2AF1, and MKRN2. Note: For C, D, and E, m6A/m6Am sites were identified using previously published miCLIP-seq data (Linder et al. 2015; PMID: 26121403). **F:** qRT-PCR showing the expression of METTL3 in HEK293 cells treated with either siMETTL3 or control siNT siRNAs. The experiments were performed in biological triplicates, and p-values were calculated using the student's t-test ($**p \leq 0.01$). Error bars represent SEM. **G:** m6A dot blot assay indicates a reduction in m6A signal in siMETTL3 treated cells in comparison with siNT control cells. Top panel was probed using anti-m6A antibody, whereas bottom panel was stained with Methylene blue. Quantification of the m6A signals is represented as a bar graph on the right ($n=3$, $*p \leq 0.05$, student's t-test). Error bars represent SEM.

### Figure S6

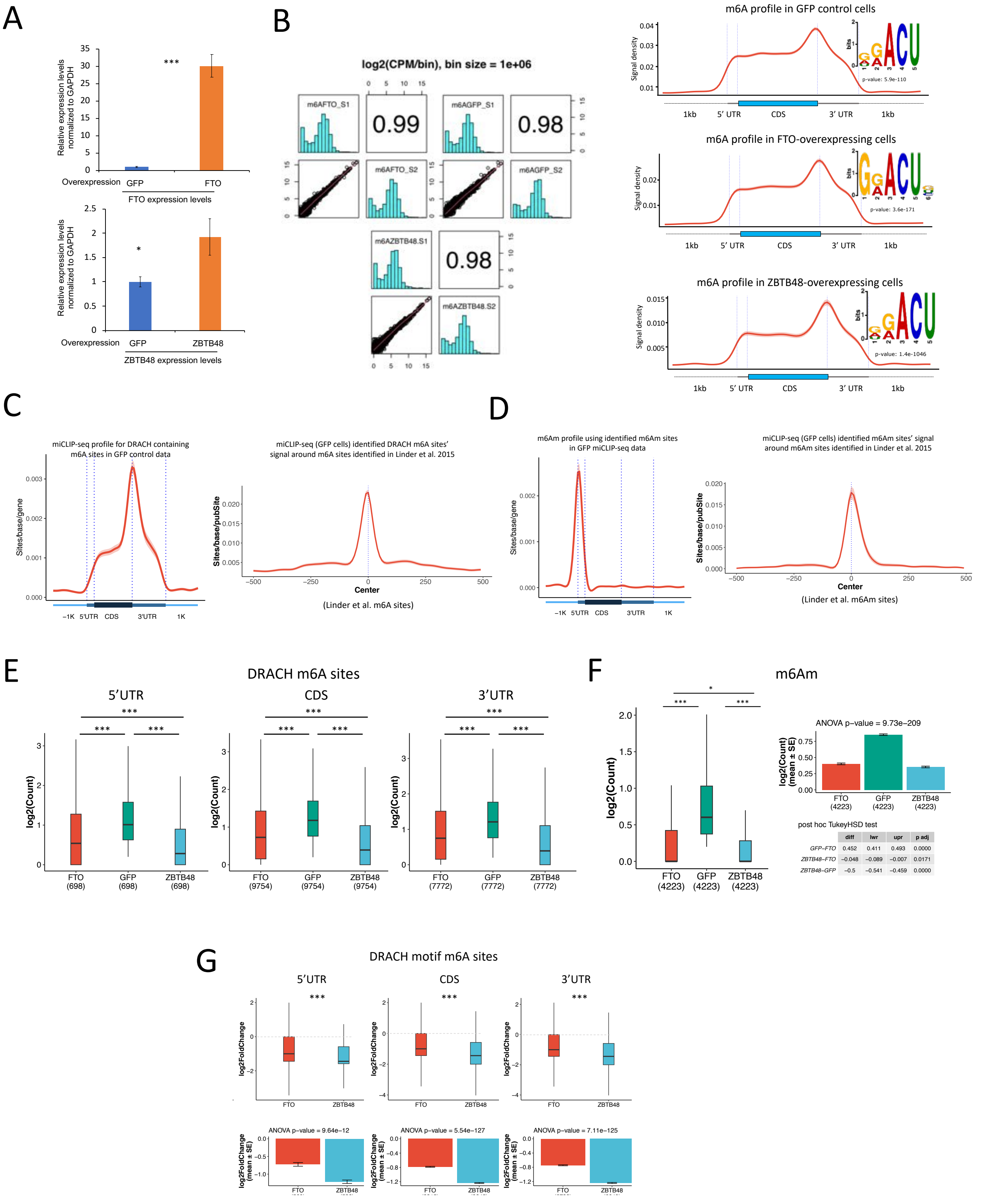


| | diff | lwr | upr | p adj |
|---|---|---|---|---|
| GFP–FTO | 0.452 | 0.411 | 0.493 | 0.0000 |
| ZBTB48–FTO | −0.048 | −0.089 | −0.007 | 0.0171 |
| ZBTB48–GFP | −0.5 | −0.541 | −0.459 | 0.0000 |

**Figure S6: ZBTB48 affects cellular m6A/m6Am. A:** qRT-PCR showing the expression levels for FTO (left) and ZBTB48 (right) in HEK293 cells overexpressing either FTO or ZBTB48, respectively. Cells expressing GFP alone were used as a control. Expression was induced using doxycycline 24 hours prior to harvesting ($***p \leq 0.001$, $*p \leq 0.05$, student's t-test, n=3). Error bars represent SEM. **B:** Left, Correlation analysis of miCLIP-seq replicates for the indicated samples. Pearson correlation co-efficient values are shown. Right, Standardized metaplot profiles showing the miCLIP-seq signal for HEK293 cells overexpressing either ZBTB48 or FTO or GFP-alone. Enriched motifs identified in each miCLIP-seq are shown above the metagene plots. The miCLIP-seq CITS were used for *de novo* motif discovery. p-value was calculated against randomly assorted sequences. **C:** Left, Metagene plot shows the miCLIP-seq profile generated using only DRACH motif containing CITS peaks in representative GFP-alone samples. Right, Signal density of DRACH motif containing CITS peaks in GFP-alone samples around previously published miCLIP-seq identified m6A sites (Linder et al. 2015; PMID: 26121403). **D:** Left, Metagene plot generated using the putative m6Am sites identified for our GFP miCLIP-seq data. Right, Signal density of m6Am sites identified for our GFP miCLIP-seq data around previously reported m6Am sites (Linder et al. 2015; PMID: 26121403). **E:** Boxplots of read counts for DRACH-motif containing m6A sites across different mRNA regions, as indicated (Mann–Whitney test, $***p \leq 0.001$; CDS: coding region). **F:** Boxplots of miCLIP read counts for m6Am sites for the indicated samples. ($***p \leq 0.001$, * $p \leq 0.05$; CDS: coding region). **G:** Box plot showing the differential methylation analysis using miCLIP data generated in either GFP-ZBTB48-overexpressing or FTO-overexpressing cells versus GFP control cells. Analysis was performed with the DESeq2 package, which accounts for the read depth in each sample, using reads corresponding to DRACH-motif-containing m6A sites. Outliers are not shown in the box plots. Dotted lines indicate the base line signal (i.e., m6A in GFP cells). Bottom bar plots indicate the mean values for the indicated groups along with p-values (ANOVA, post hoc Tukey HSD).

### Figure S7

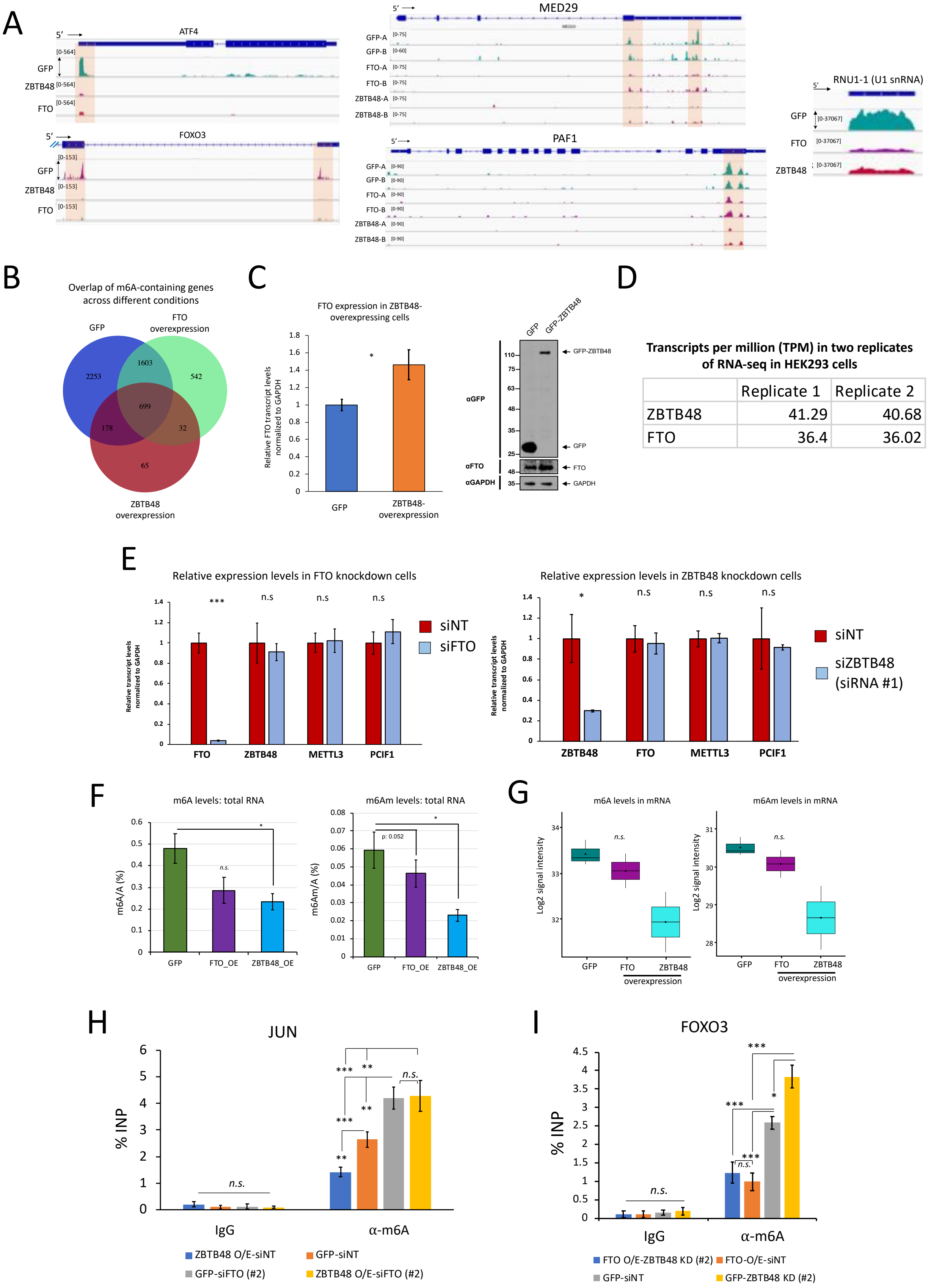


**Transcripts per million (TPM) in two replicates of RNA-seq in HEK293 cells**

| | Replicate 1 | Replicate 2 |
|---|---|---|
| ZBTB48 | 41.29 | 40.68 |
| FTO | 36.4 | 36.02 |

**Figure S7: ZBTB48 modulates cellular m6A/m6Am through FTO. A:** Genome browser snapshots of the indicated genes to show the signal density of miCLIP-seq data for either ZBTB48-overexpressing, or FTO-overexpressing, or GFP-alone samples. The locations of putative m6A sites are highlighted on each gene track. **B:** Venn diagram showing the overlap of m6A-containing genes identified in various samples, as indicated. **C:** Left, qRT-PCR shows the expression levels of FTO in cells overexpressing either GFP-ZBTB48 or GFP-alone. The expression of GFP-ZBTB48 or GFP was induced 24 hours prior to harvesting. The experiments were performed in biological triplicates, and p-values were calculated using the student's t-test ($*p \leq 0.05$). Error bars represent SEM. Right, FTO protein levels were determined using Western blotting analysis in whole cell extracts prepared from either ZBTB48-overexpressing cells or GFP-alone. Blots were probed with the indicated antibodies. **D:** ZBTB48 and FTO mRNA expression levels in HEK293 cells represented as transcripts per million. Analysis is based on RNA-seq data from siNT-treated cells. **E:** Bar graphs representing qRT-PCR results to examine the differential expression of selected genes in either siFTO (left) or siZBTB48 (right) cells in comparison with siNT controls. The experiments were performed in biological triplicates, and p-values were calculated using the student's t-test ($***p \leq 0.001$, $**p \leq 0.01$, $*p \leq 0.05$, n.s.: non-significant). Error bars represent SEM. **F:** LC-MS/MS analysis shows that overexpression of GFP-ZBTB48 or FTO leads to a decrease in m6A/A (left) and m6Am/A (right) levels in total RNA, in comparison with the control GFP cells. Error bars show SEM ($*p \leq 0.05$; student's t-test). **G:** LC-MS/MS shows that overexpression of GFP-ZBTB48 or FTO leads to substantially decreased m6A/A (left) and m6Am/A (right) levels in polyadenylated mRNAs, in comparison with the control GFP cells. The difference in m6A and m6Am levels was not significant (P-values > 0.05) due to variation in the biological replicates; however, the overall trend indicates a substantial reduction in m6A/m6Am levels in both ZBTB48- and FTO-overexpressing cells. **H-I**: Bar plots showing the m6A levels for JUN (H) and FOXO3 (I) transcripts estimated using m6A-RIP-qPCR experiments performed under the indicated conditions. Data are represented as % input (biological replicates n = 5, student's t-test, $***p \leq 0.001$, $**p \leq 0.01$, $*p \leq 0.05$, n.s.: non-significant, error bars denote SEM). Note: In figures H-I, FTO and ZBTB48 KD were performed using siFTO #2 and siZBTB48#2, respectively.

### Figure S8

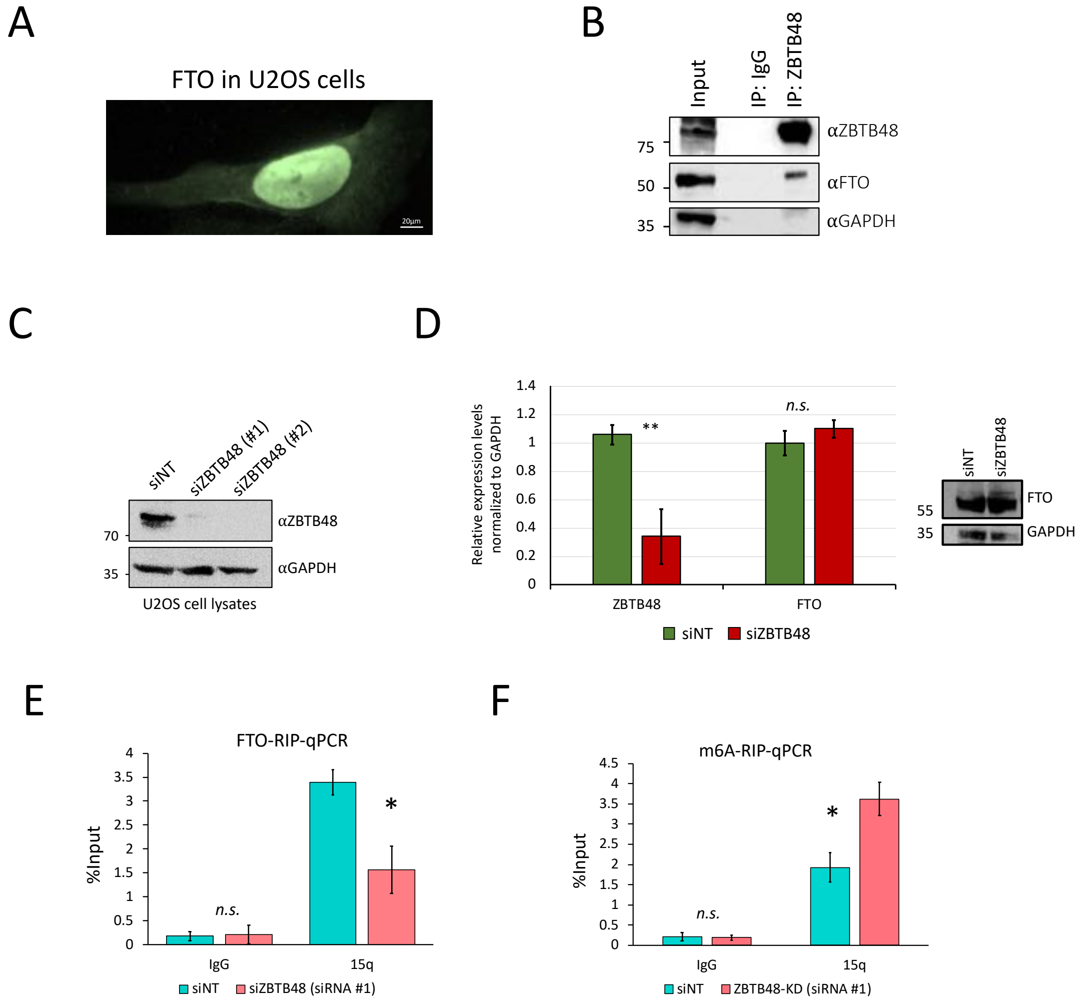
A
FTO in U2OS cells
20µm
B
Input
IP: IgG
IP: ZBTB48
75
50
35
αZBTB48
αFTO
αGAPDH
C
siNT
siZBTB48 (#1)
siZBTB48 (#2)
70
35
αZBTB48
αGAPDH
U2OS cell lysates
D
Relative expression levels normalized to GAPDH
1.4
1.2
1
0.8
0.6
0.4
0.2
0
**
n.s.
ZBTB48
FTO
siNT
siZBTB48
55
35
FTO
GAPDH
E
FTO-RIP-qPCR
%Input
4
3.5
3
2.5
2
1.5
1
0.5
0
n.s.
*
IgG
15q
siNT
siZBTB48 (siRNA #1)
F
m6A-RIP-qPCR
%Input
4.5
4
3.5
3
2.5
2
1.5
1
0.5
0
n.s.
*
IgG
15q
siNT
ZBTB48-KD (siRNA #1)

**Figure S8: ZBTB48 recruits FTO to TERRA in U2OS cells. A:** IF analysis to examine the localization of FLAG-FTO in U2OS cells. Experiment was performed using an anti-FLAG antibody. **B:** Interaction of ZBTB48 with FTO detected by co-immunoprecipitation in U2OS cells. IPs were performed with ZBTB48 antibody (10 ug) using whole-cell lysates. Cell lysates were treated with Benzonase (nuclease) prior to IPs. Blots were probed with the indicated antibodies. **C:** Western blotting analysis in whole cell lysates prepared from U2OS cells treated either with siZBTB48 #1 or siZBTB48 #2. Blots were probed with the indicated antibodies. **D:** Left, Bar graphs represent qRT-PCR results to examine the expression of ZBTB48 and FTO in ZBTB48 knockdown U2OS cells (siZBTB48 # 1). The experiments were performed in biological triplicates, and p-values were calculated using the student's t-test ($***p \leq 0.001$, $**p \leq 0.01$, $*p \leq 0.05$, n.s.: non-significant, Error bars represent SEM). Right, Western blotting analysis in whole cell lysates prepared from U2OS cells treated with siZBTB48 #1 to examine FTO protein levels. Blots were probed with the indicated antibodies. **E:** Bar graph showing FTO RIP-qPCR in siNT- or siZBTB48-treated (siZBTB48 #1) U2OS cells. TERRA was detected using specific primers against the 15q chromosome. IPs with IgG served as a negative control. **F:** Bar plot showing the m6A levels for TERRA transcripts (15q) estimated using m6A-RIP-qPCR in ZBTB48 knockdown (siZBTB48 #1) and siNT-treated U2OS cells. Data are represented as % input (biological replicates $n = 5$, student's t-test, $***p \leq 0.001$, $**p \leq 0.01$, $*p \leq 0.05$, n.s.: non-significant, error bars denote SEM).

### Figure S9

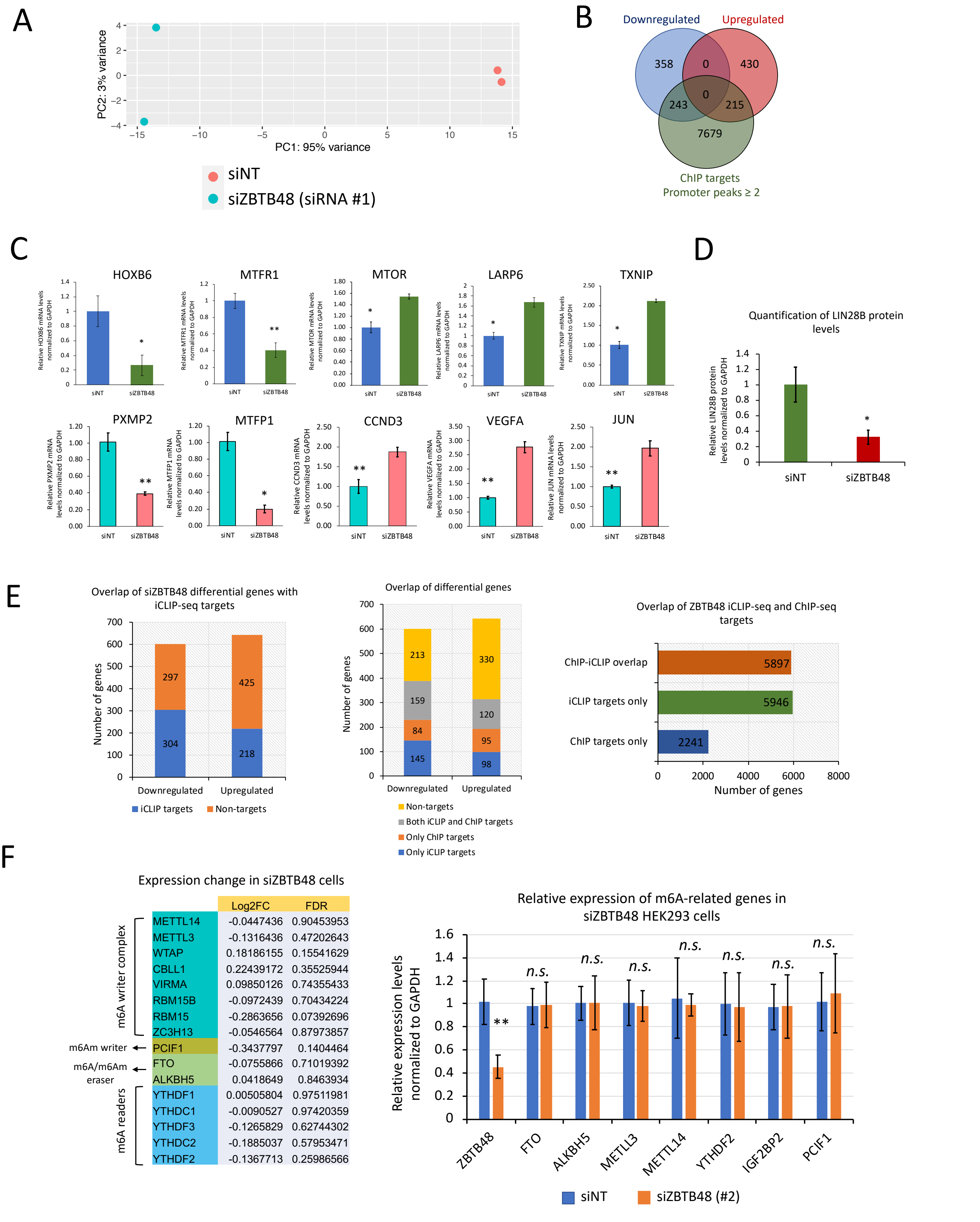


| | Log2FC | FDR |
|---|---|---|
| METTL14 | -0.0447436 | 0.90453953 |
| METTL3 | -0.1316436 | 0.47202643 |
| WTAP | 0.18186155 | 0.15541629 |
| CBLL1 | 0.22439172 | 0.35525944 |
| VIRMA | 0.09850126 | 0.74355433 |
| RBM15B | -0.0972439 | 0.70434224 |
| RBM15 | -0.2863656 | 0.07392696 |
| ZC3H13 | -0.0546564 | 0.87973857 |
| PCIF1 | -0.3437797 | 0.1404464 |
| FTO | -0.0755866 | 0.71019392 |
| ALKBH5 | 0.0418649 | 0.8463934 |
| YTHDF1 | 0.00505804 | 0.97511981 |
| YTHDC1 | -0.0090527 | 0.97420359 |
| YTHDF3 | -0.1265829 | 0.62744302 |
| YTHDC2 | -0.1885037 | 0.57953471 |
| YTHDF2 | -0.1367713 | 0.25986566 |

**Figure S9: ZBTB48 depletion alters gene expression. A:** Principal component analysis (PCA) of ZBTB48 knockdown RNA-seq replicates (siZBTB48 # 1). **B:** Venn diagram depicts the overlap between differentially expressed genes in siZBTB48 HEK293 cells and ZBTB48 ChIP targets. For ChIP targets, only those genes that had at least two peaks in the promoter region (1kB upstream and 100nt downstream of TSS) were considered. **C:** Bar graphs representing qRT-PCR results to examine the differential expression of selected ZBTB48 RNA targets in ZBTB48 knockdown cells (student's t-test, n=3, $***p \leq 0.001$, $**p \leq 0.01$, $*p \leq 0.05$). Error bars represent SEM. **D:** Bar graph representation for the quantification of LIN28B protein levels in siZBTB48-treated cells (representative blots shown in Figure 5D). Quantification was performed using ImageJ software (https://imagej.nih.gov/ij/download.html; last accessed June 2023) (student's t-test, n=3, $*p \leq 0.05$). **E:** Left, Bar graph depicts the overlap of ZBTB48 target genes (as identified from ZBTB48 iCLIP-seq) with differentially expressed genes in siZBTB48. Middle, Bar graph depicts the overlap of ZBTB48 target genes at either iCLIP, or ChIP, or both ChIP and iCLIP level, or non-targets with siZBTB48 differential genes. Right, Bar graph indicates the overlap between ZBTB48 iCLIP-seq and ChIP-seq targets. For ChIP-seq targets, only those genes that had at least two peaks in the promoter region (1kB upstream and 100nt downstream of TSS) were considered. **F:** Left, Log2fold change values of m6A-related genes based on differential expression analysis of RNA-seq data (siZBTB48 vs control siNT) in HEK293 cells. Right, Bar graphs show qRT-PCR results examining the differential expression of selected genes in siZBTB48 (siRNA#2) HEK293 cells in comparison with siNT controls ($**p \leq 0.01$, n.s.: non-significant, student's t-test, biological replicates (n) =3).

### Figure S10

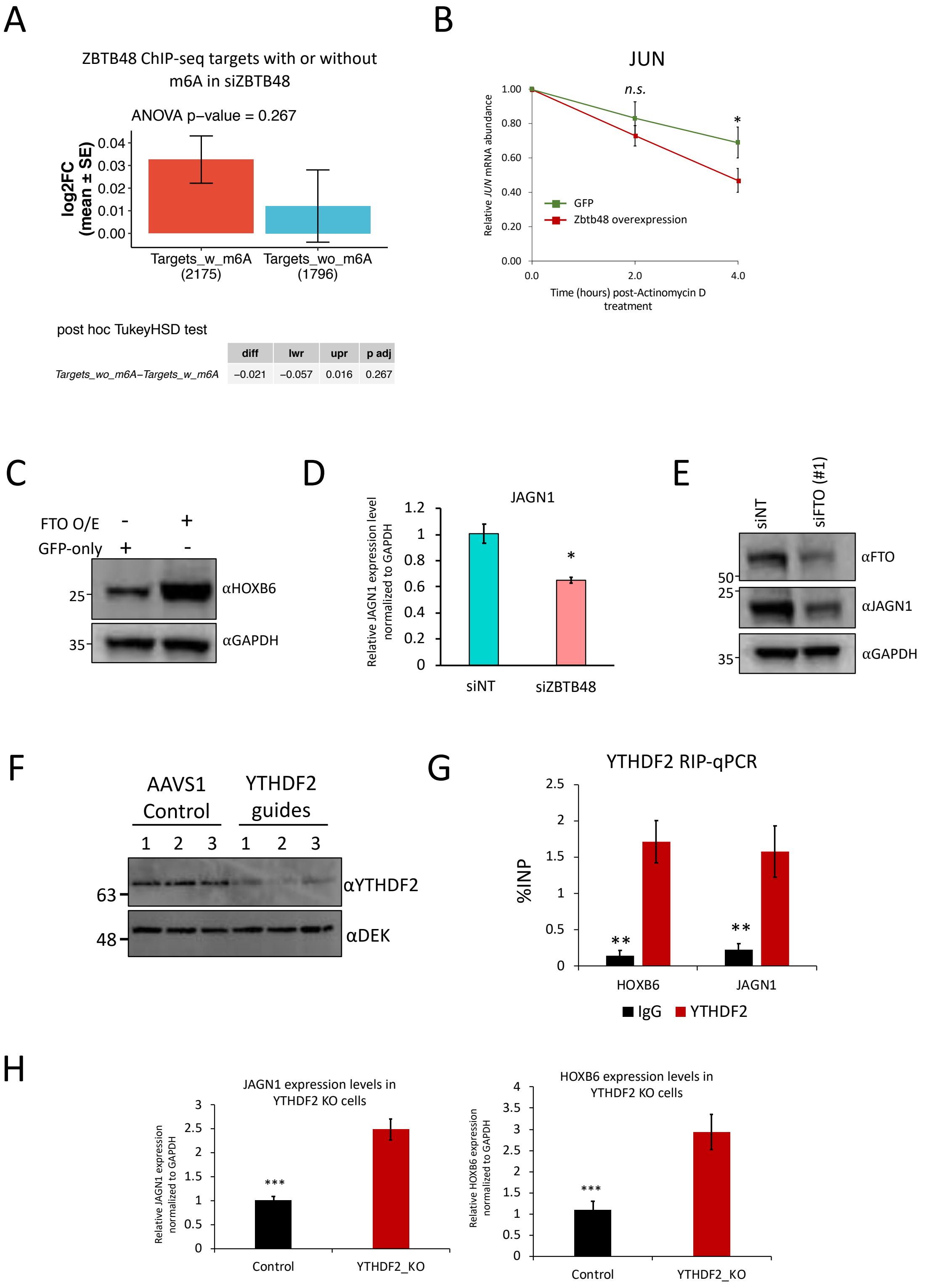


| | diff | lwr | upr | p adj |
|---|---|---|---|---|
| Targets_wo_m6A–Targets_w_m6A | −0.021 | −0.057 | 0.016 | 0.267 |

**Figure S10: ZBTB48 mediated regulation of m6A through FTO affects gene expression via YTHDF2. A:** Bar graph depicts mean expression change in siZBTB48 cells for ZBTB48 ChIP targets with or without m6A (ANOVA, post hoc Tukey HSD p-value is shown). **B:** qRT-PCR analysis of m6A-containing JUN mRNAs after treating GFP or GFP-ZBTB48 overexpressing cells with Actinomycin D for the indicated times (student's t-test, n=3, $*p \leq 0.05$, n.s.: non-significant). **C:** Western blotting analysis in whole cell lysates to show the effect of FTO overexpression, as indicated. Blots were probed with the indicated antibodies. **D:** Bar graph shows the qRT-PCR results examining the expression of JAGN1 mRNA in siZBTB48 (siRNA#2) HEK293 cells in comparison with siNT controls ($*p \leq 0.05$, n.s.: non-significant, student's t-test, biological replicates (n) =3). **E:** Western blotting analysis in whole cell lysates prepared from cells treated with either siFTO or siNT. Blots were probed with the indicated antibodies. **F:** Western blotting analysis in whole cell extracts prepared from HEK293T cells transfected with lentivirus with either control AAVS1 or YTHDF2 targeting CRISPR guides. Blots were probed with the indicated antibodies. **G:** Bar graph representation of YTHDF2 RIP-qPCR showing binding of YTHDF2 to the indicated transcripts (biological replicates n = 3, student's t test, $**p \leq 0.01$, error bars: SEM). **H:** Bar graphs show the qRT-PCR results examining the expression of JAGN1 and HOXB6 mRNAs in YTHDF2 KO HEK293 cells in comparison with controls ($***p \leq 0.001$, student's t-test, biological replicates (n) =3).

### Figure S11

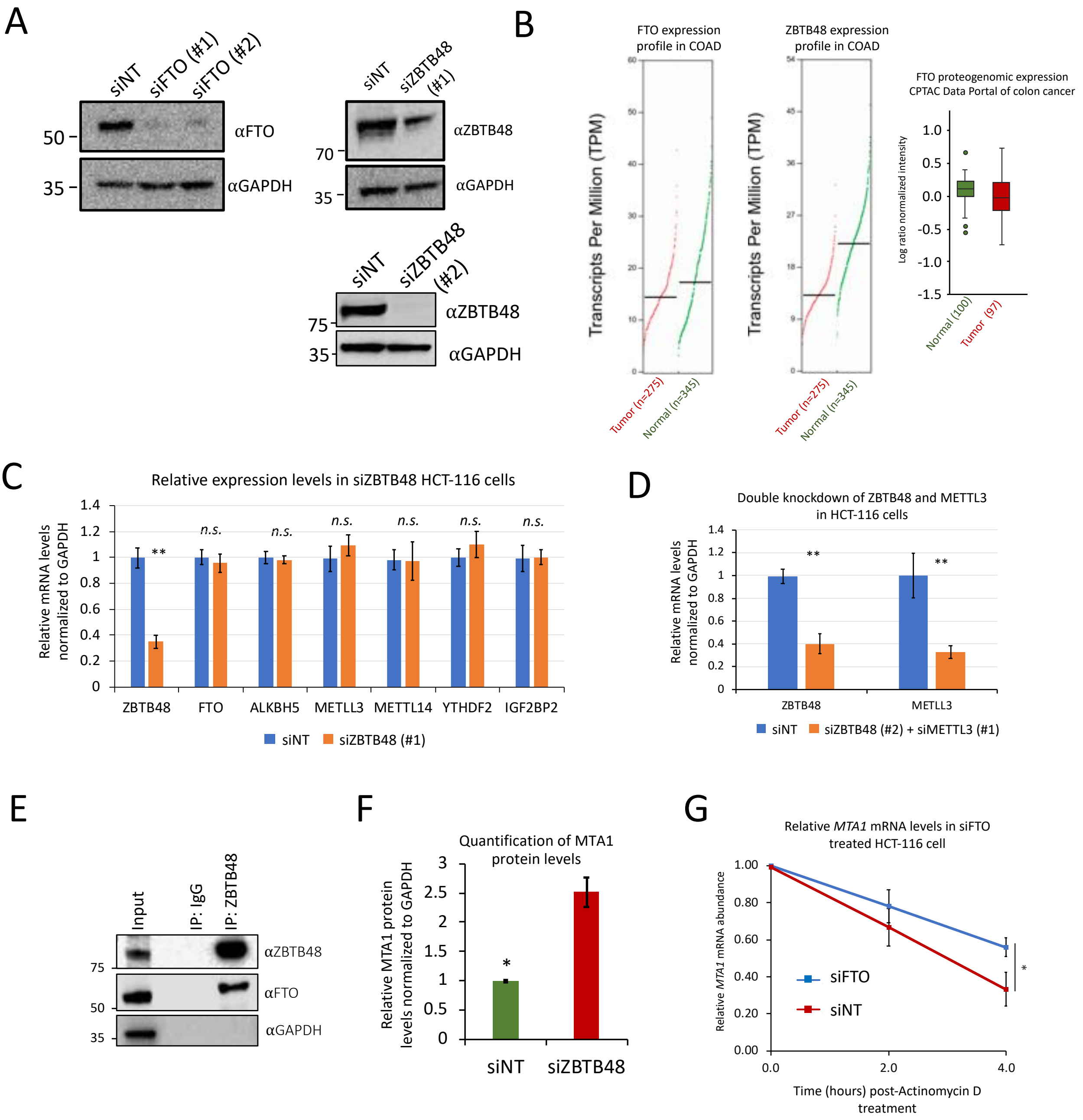

A
siNT
siFTO (#1)
siFTO (#2)
αFTO
αGAPDH
50
35
siNT
siZBTB48 (#1)
αZBTB48
αGAPDH
70
35
siNT
siZBTB48 (#2)
αZBTB48
αGAPDH
75
35
B
FTO expression profile in COAD
ZBTB48 expression profile in COAD
Transcripts Per Million (TPM)
Tumor (n=275)
Normal (n=345)
FTO proteogenomic expression
CPTAC Data Portal of colon cancer
Log ratio normalized intensity
1.0
0.5
0.0
-0.5
-1.0
-1.5
Normal (100)
Tumor (97)
C
Relative expression levels in siZBTB48 HCT-116 cells
Relative mRNA levels normalized to GAPDH
1.4
1.2
1
0.8
0.6
0.4
0.2
0
**
n.s.
ZBTB48
FTO
ALKBH5
METTL3
METTL14
YTHDF2
IGF2BP2
siNT
siZBTB48 (#1)
D
Double knockdown of ZBTB48 and METTL3 in HCT-116 cells
Relative mRNA levels normalized to GAPDH
1.4
1.2
1
0.8
0.6
0.4
0.2
0
**
ZBTB48
METTL3
siNT
siZBTB48 (#2) + siMETTL3 (#1)
E
Input
IP: IgG
IP: ZBTB48
αZBTB48
αFTO
αGAPDH
75
50
35
F
Quantification of MTA1 protein levels
Relative MTA1 protein levels normalized to GAPDH
3
2.5
2
1.5
1
0.5
0
*
siNT
siZBTB48
G
Relative MTA1 mRNA levels in siFTO treated HCT-116 cell
Relative MTA1 mRNA abundance
1.00
0.80
0.60
0.40
0.20
0.00
siFTO
siNT
*
0.0
2.0
4.0
Time (hours) post-Actinomycin D treatment

**Figure S11: ZBTB48 is deregulated in colorectal cancer. A:** Western blotting analysis in whole cell lysates prepared from HEK293 cells treated with the indicated siRNAs. Blots were probed with the indicated antibodies. **B:** The Cancer Genome Atlas (TCGA) RNA-seq expression values for FTO (left) and ZBTB48 (middle) in normal tissues (green) versus primary tumors (red) (Note that COAD stands for colon adenocarcinoma). TCGA analysis was performed using the online tool GEPIA (http://gepia.cancer-pku.cn/index.html). Right, comparison of the proteogenomic expression of FTO in tumor (red) and normal tissues (green) from the CPTAC Data Portal of colon cancer (https://pdc.cancer.gov/pdc/browse/filters/pdc_study_id:PDC000109%7CPDC000116%7CPDC000117). **C:** Bar graphs show qRT-PCR results examining the differential expression of selected genes in siZBTB48 HCT-116 cells in comparison with siNT controls ($**p \leq 0.01$, n.s.: non-significant, student's t-test, n=3). Error bars represent SEM. **D:** qRT-PCR to evaluate the knockdown of ZBTB48 and METTL3 in HCT-116 cells treated with a combination of siRNAs (siZBTB48#2+siMETTL3). Error bars represent SEM ($***p \leq 0.001$, $**p \leq 0.01$, n.s.: non-significant, student's t-test, n=3). **E:** Interaction of ZBTB48 with FTO detected by co-immunoprecipitation in HCT116 cells. IPs were performed with ZBTB48 antibody (10 ug) using whole-cell lysates. Cell lysates were treated with Benzonase (nuclease) prior to IPs. Blots were probed with the indicated antibodies. **F:** Bar graph representation for the quantification of MTA1 protein levels in siZBTB48-treated cells (representative blots shown in Figure 6G). Quantification was performed using ImageJ software (https://imagej.nih.gov/ij/download.html; last accessed June 2023) (student's t-test, n=3, $*p \leq 0.05$). **G:** qRT-PCR analysis of *MTA1* expression in siFTO or siNT-treated HCT-116 cells after treating cells with Actinomycin D for the indicated times ($*p \leq 0.05$, student's t test, n=3). Error bars represent SEM.

### Figure S12

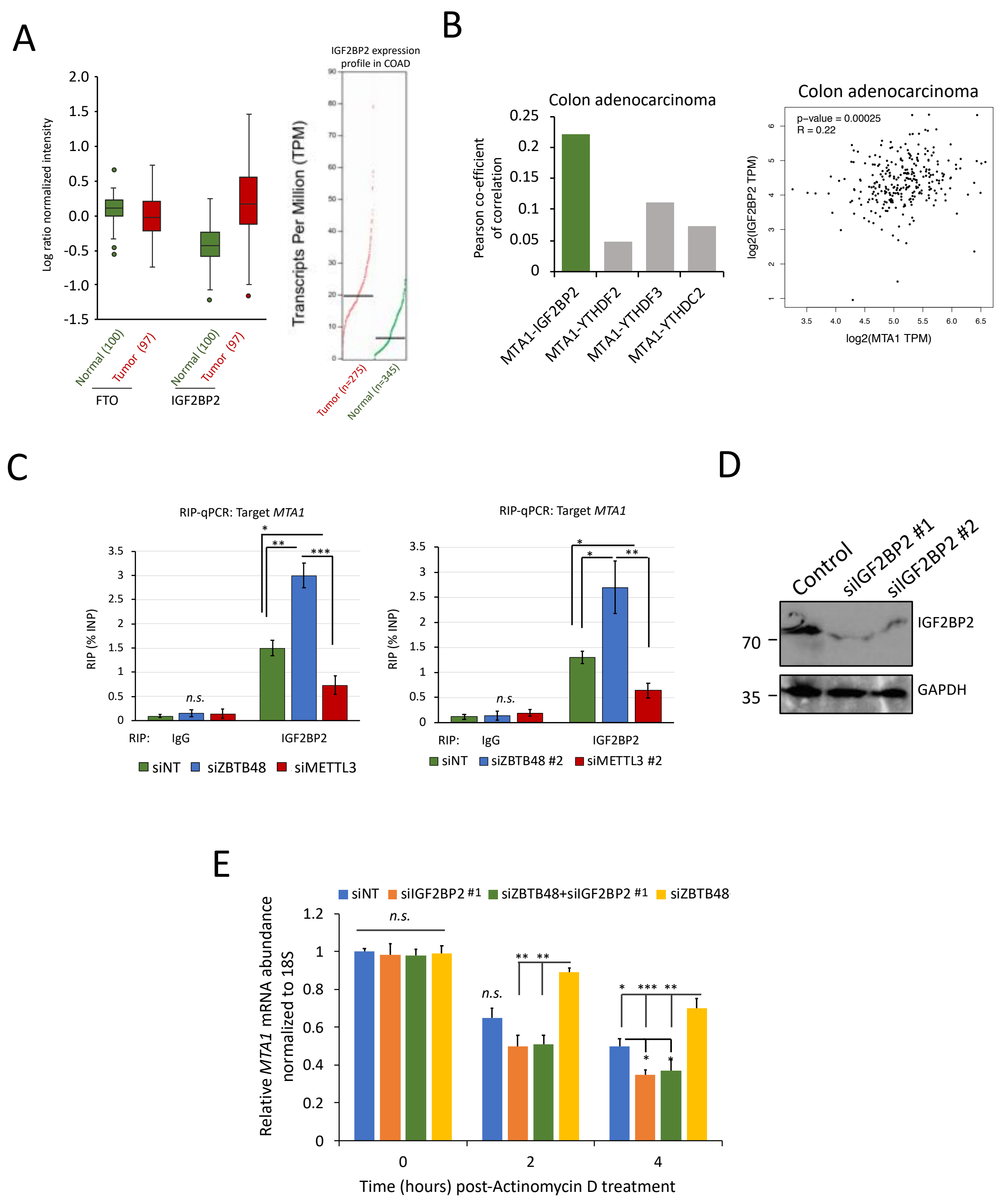

A
Log ratio normalized intensity
Normal (100)
Tumor (97)
FTO
IGF2BP2
IGF2BP2 expression profile in COAD
Transcripts Per Million (TPM)
Tumor (n=275)
Normal (n=345)
B
Colon adenocarcinoma
Pearson co-efficient of correlation
MTA1-IGF2BP2
MTA1-YTHDF2
MTA1-YTHDF3
MTA1-YTHDC2
Colon adenocarcinoma
p-value = 0.00025
R = 0.22
log2(IGF2BP2 TPM)
log2(MTA1 TPM)
C
RIP-qPCR: Target MTA1
RIP (% INP)
n.s.
RIP: IgG IGF2BP2
siNT siZBTB48 siMETTL3
RIP-qPCR: Target MTA1
siNT siZBTB48 #2 siMETTL3 #2
D
Control
siIGF2BP2 #1
siIGF2BP2 #2
70
35
IGF2BP2
GAPDH
E
siNT siIGF2BP2 #1 siZBTB48+siIGF2BP2 #1 siZBTB48
Relative MTA1 mRNA abundance normalized to 18S
n.s.
Time (hours) post-Actinomycin D treatment

**Figure S12: ZBTB48 regulates MTA1 expression through IGF2BP2. A:** Left, Boxplot showing the proteogenomic expression of FTO and IGF2BP2 in tumor (red) and normal tissues (green) from the CPTAC Data Portal of colon cancer. Right, TCGA RNA-seq expression values for IGF2BP2 in normal tissues (green) vs primary tumors (red). **B:** Left, Bar plot shows the Pearson co-efficient of correlation (r) values for the indicated pairs based on the TCGA RNA-seq data for COAD. Right, Scatter plot for IGF2BP2-to-MTA1 expression correlations. Each dot represents a single TCGA sample. 'R' denotes the Pearson correlation coefficient. TPM: Transcripts per million. **C:** Bar graph representation of IGF2BP2 RIP-qPCR for MTA1 in HCT-116 cells treated with the indicated siRNAs. The experiments were performed in biological triplicates, and p-values were calculated using the student's t test (∗∗∗$p \leq 0.001$, ∗∗$p \leq 0.01$, ∗$p \leq 0.05$, n.s.: non-significant). Error bars represent SEM. **D:** Western blotting analysis in whole cell lysates prepared from HCT-116 cells treated with the indicated siRNAs. Blots were probed with the indicated antibodies. **E:** qRT-PCR analysis of MTA1 in cells treated with siNT, siZBTB48 (siRNA#1), siIGF2BP2 (siRNA#1), or a combination of siZBTB48 and siIGF2BP2. For each time point, relative transcript levels were normalized to 18S rRNA and time point "0".
